# Unscrambling cancer genomes via integrated analysis of structural variation and copy number

**DOI:** 10.1101/2020.12.03.410860

**Authors:** Charles Shale, Jonathan Baber, Daniel L. Cameron, Marie Wong, Mark J. Cowley, Anthony T. Papenfuss, Edwin Cuppen, Peter Priestley

## Abstract

Complex somatic genomic rearrangement and copy number alterations (CNA) are hallmarks of nearly all cancers. Whilst whole genome sequencing (WGS) in principle allows comprehensive profiling of these events, biological and clinical interpretation remains challenging. We have developed LINX, a novel algorithm which allows interpretation of short-read paired-end WGS derived structural variant and CNA data by clustering raw structural variant calls into distinct events, predicting their impact on the local structure of the derivative chromosome, and annotating their functional impact on affected genes. Novel visualisations facilitate further investigation of complex genomic rearrangements. We show that LINX provides insights into a diverse range of structural variation events including single and double break-junction events, mobile element insertions, complex shattering and high amplification events. We demonstrate that LINX can reliably detect a wide range of pathogenic rearrangements including gene fusions, immunoglobulin enhancer rearrangements, intragenic deletions and duplications. Uniquely, LINX also predicts chained fusions which we demonstrate account for 13% of clinically relevant oncogenic fusions. LINX also reports a class of inactivation events we term homozygous disruptions which may be a driver mutation in up to 8.8% of tumors including frequently affecting *PTEN*, *TP53* and *RB1*, and are likely missed by many standard WGS analysis pipelines.

## Introduction

Copy number alteration (CNA) and structural variation (SV) are two of the key classes of somatic mutations in cancer with approximately 90% of tumor genomes undergoing significant rearrangements (Zack et al. 2013). However, the mechanisms driving and the consequences of genomic rearrangements in cancer are less well understood than for point mutation events. This is due both to the relative paucity of whole genome sequencing data which is required for comprehensive SV analysis, and also to the fact that genomic rearrangements have significant diversity. Many rearrangements involve a high degree of complexity with individual events resulting in multiple or even hundreds of breaks (Stephens et al. 2011; Garsed et al. 2014). Interpretation of these highly rearranged genomes is challenging but simultaneously highly relevant for the identification of driver events that may function as biomarkers or druggable targets.

LINX is a novel SV interpretation tool, which integrates CNA and SV calling derived from WGS data and comprehensively clusters, chains and classifies genomic rearrangements. The motivation for this is twofold: first from a biological perspective accurate interpretation can allow better insight into distinct mechanisms of rearrangements in tumorigenesis and second from a clinical perspective in accurately predicting the functional impact of structural rearrangements in both gene fusions and disruptions. A number of previous tools have been developed to analyse the roles of certain rearrangement event types in tumorigenesis such as chromothripsis (Stephens et al. 2011), chromoplexy (Baca et al. 2013), LINE insertions (Rodriguez-Martin et al. 2020), or amplification mechanisms (Deshpande et al. 2019). Clustering methodologies have also been used previously to propose signatures of structural rearrangement (Nik-Zainal et al. 2016; Li et al. 2020). LINX goes further than just integrating the functionality of each of these previous tools both by classifying all classes of rearrangements in each tumor genome, and by predicting the local chained structure of the derivative chromosome as well as the functional impact of the rearrangement in a single application.

## Results

### LINX algorithm

The input for LINX is a base-pair consistent segmented copy number and SV callset from the previously described tools PURPLE (Priestley et al. 2019) and GRIDSS (Cameron et al., n.d.). A docker image is available which allows GRIDSS2, PURPLE & LINX to be run in a single pipeline from BAM input.

There are 4 key steps in the LINX algorithm (FIGURE 1A-B, SUPP FILE 1). First, LINX annotates individual breakends derived from GRIDSS with several basic geometric and genomic properties which are important to the clustering and chaining algorithm. This includes whether each breakend is part of a foldback inversion, flanks a region of loss of heterozygosity, or is in a well known fragile site region (Priestley et al. 2019; Dillon, Burrow, and Wang 2010). LINX also uses a combination of previously known line element source information (Rodriguez-Martin et al. 2020) and a procedure to identify novel suspected mobile LINE source elements based on both the local breakpoint structure and signals of poly-A sequence insertions.

**FIGURE 1:**
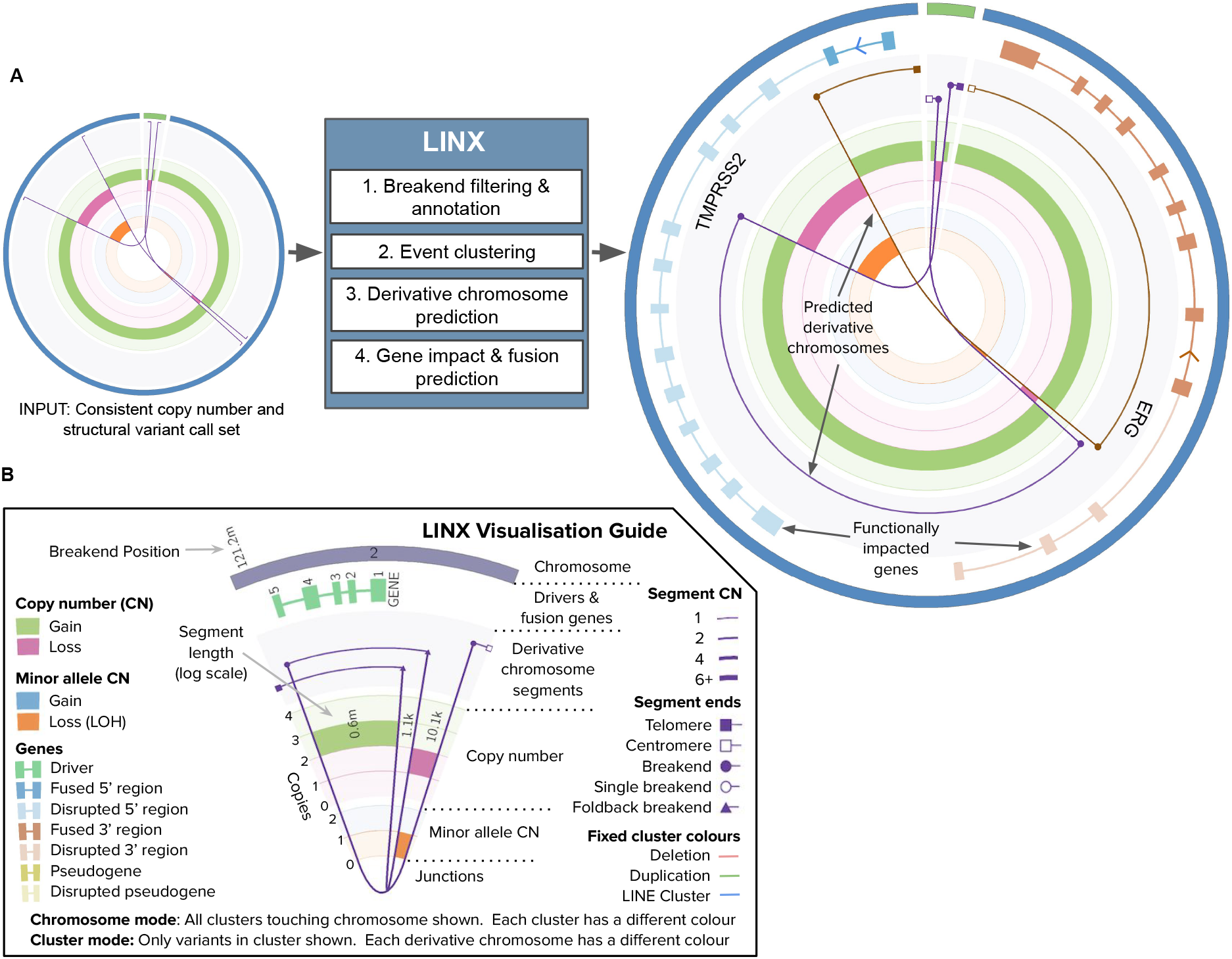
LINX schematic and visualisations a) The LINX algorithm works in 4 steps to annotate, cluster, chain and determine the functional impact of an integrated copy number. The circos on the left represents the input of LINX and shows 3 structural variants (purple lines) affecting 2 chromosomes (outer track in green and blue) with consistent copy number breakpoints (middle track showing green for gain and red for loss). The circos on the right shows example output of LINX including the chaining of the variants into 2 continuous predicted derivative chromosomes (lines in brown and purple), and a canonical TMPRSS2_ERG fusion (genes depicted in blue and light brown on 2nd outer circle with fused exons showing darker shading) on one of the 2 predicted chromosomes. b) A detailed guide to the visualisations produced by LINX

Second, LINX performs a clustering routine to group raw structural variants into distinct rearrangement ‘events’. LINX defines a rearrangement event as 1 or more junctions which likely occurred proximately in time and transformed the genome from one stable configuration to another. Events can range from a simple deletion or tandem duplication to a complex event including chromothripsis event or breakage fusion bridge (McClintock 1941) cascade. The fundamental principle for clustering in LINX is to join breakpoints where it is highly unlikely that they could have occurred independently. Rather than a single rule such as clustering variants into events based solely on proximity (Hadi et al., n.d.) or variants which form a ‘deletion bridge’ (Baca et al. 2013), LINX relies on a set of 11 independent rules (SUPP FILE 1). These include clustering variants which are very close in proximity (<5kb between breakends); clustering breakends which together delimit an LOH event, homozygous deletion or region of high major allele copy number; clustering translocations which share common arms at both ends; clustering inversions, long deletion & long tandem duplication variants which directly overlap each other; and clustering all foldback inversions which occur on the same chromosome arm.

Third, after resolving all variants into clusters, LINX determines the derivative chromosome structures via a chaining algorithm. To do this, LINX considers all pairs of facing breakends on each chromosomal arm within each cluster and iteratively prioritises which pair is most likely to be joined. The chaining logic imposes allele specific copy number constraints at all points on each chromosome and also the biological constraint that chromosomes are not permitted without a centromere unless strict criteria relating to detection of extrachromosomal DNA (ecDNA or double minutes) are met. Foldback and complex replication-based duplication events are explicitly modelled to allow chaining of clusters of variable junction copy number and high amplification and heuristics allow classification of the likely amplification mechanism (see detailed methods) as ecDNA or breakage fusion bridge. Overall, the chaining prioritisation scheme is designed to be error tolerant and aims to maximise the chance that each individual breakend is linked correctly to the next breakend on the derivative chromosome. However, due to multiple possible paths, upstream sources of error and missing information, the prediction is representative only and unlikely to be completely correct in all details, especially in the case of highly complex clusters.

The fourth and final step in LINX is to annotate the gene impact of breakends to predict gene disruptions and fusions. Any breakend overlapping or in the upstream region of an Ensembl transcript (Yates et al. 2020) is annotated with its position and orientation relative to the strand of the gene and flag if it is disruptive to the gene (ie. if it impacts an exonic region). Gene fusions are called by searching for correctly oriented splice acceptor and donor pairs on the derivative chromosomes predicted by LINX, including fusions which may span multiple break junctions (Anderson et al. 2018). To meet the fusion calling criteria, the breakends must also connect to viable contexts in each gene and not be terminated by further breakends in the chain on either 5’ or 3’ partner end (see methods for full details). As complex rearrangements can result in multiple unexpressed gene fusions, LINX streamlines clinical interpretation by maintaining curated lists of known pathogenic fusion gene pairs, 5’ and 3’ promiscuous fusion genes and marks such fusions as reportable. Finally, LINX leverages the catalog of mutated cancer drivers determined by PURPLE across a panel of oncogenes and tumor suppressor genes [SUPP TABLE 1] (Priestley et al. 2019) and determines which SV clusters contributed to each amplification, deletion and LOH event in the sample.

### Pan-cancer analysis

To demonstrate the functionality of LINX we have run it on a pan cancer cohort of 4,358 paired tumor-normal whole genome sequenced (median of 106x and 38x paired-end sequencing coverage, respectively) adult metastatic cancer samples from Hartwig Medical Foundation (referred to as HMF cohort) (Priestley et al. 2019) [SUPP TABLE 2]. 1,924 of these had matched whole transcriptome sequencing data based on RNAs-seq and were used for orthogonal validation where appropriate. Overall we found a mean of 324 rearrangement junctions per sample with the highest rates in esophagus (mean = 753) and stomach (mean = 647) tumors and lowest rates in thyroid (mean = 102) and neuroendocrine (mean = 109) tumors [FIGURE 2A,SUPP TABLE 3]. Event classification by LINX highlighted the diversity and tumor type specificity of rearrangement mechanisms with deletions, tandem duplications, LINE insertions and complex events (defined as events with 3 or more junctions) found to be the largest classes of rearrangements in agreement with previous pan-cancer whole genome analysis(Li et al. 2020). We examined each of these event classifications in detail as follows.

**FIGURE 2:**
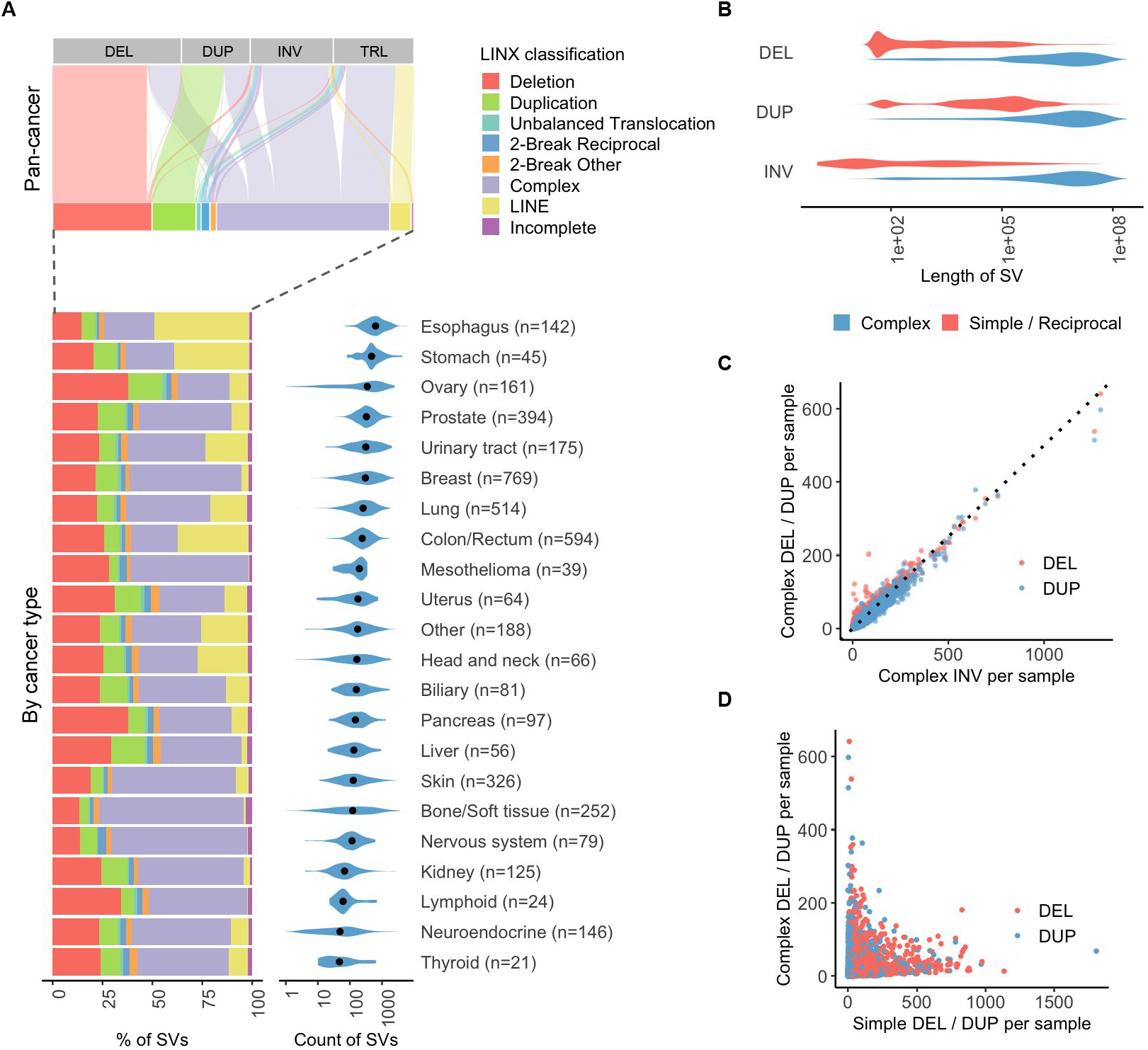
Landscape of genomic rearrangements a) Top panel shows an alluvial plot depicting the proportional assignment of each of the raw structural variant types (TRL = translocation,INV=inversion,DEL=deletion, DUP=duplication) to LINX classification. The LINX classifications are further broken down by tumor type in a relative bar chart in the left lower panel. The right lower panel shows the distribution of the number of structural variants per sample grouped by tumour type with the black dots indicating the median values. b) Length distribution of notional deletion, duplications and non foldback inversions for both simple rearrangements and complex clusters (containing 3 or more variants). Note that foldback inversions have a distinct length distribution and are shown separately in extended figure 4A. c) Counts of deletions and duplications in complex clusters per sample both closely follow a 1:2 ratio (indicated by dotted line) compared to inversions as expected by random rearrangements following catastrophic events d) Counts of simple deletions and duplications per sample are not correlated to counts of deletions and duplications in complex clusters

#### Classification of simple and complex rearrangement events

Classification of event types in LINX can considerably simplify interpretation of a cancer genome. An important use case is to distinguish simple events driven by a single break resulting in deletions and duplications [EXT FIGURE 1] from variants which are notionally called as deletions and duplications by the SV caller but may be part of a more complex event. Clean mutational profiles for simple deletions and duplications are important for downstream applications such as signature analysis (Li et al. 2020) and in particular HR deficiency classification (Davies et al. 2017)),(Nguyen et al. 2020) which is associated with both short deletions and tandem duplications and may be relevant to treatment.

In the HMF cohort, we find that lengths of deletions and duplications classified as simple events are notably shorter than those clustered in complex events [FIGURE 2B]. Moreover, the simple deletions and duplications show distinct characteristic length peaks, some of which have been previously shown to be associated with *BRCA1*, *BRCA2*, and *CDK12* inactivation or *CCNE1* amplification (Menghi et al. 2018), as well as a short DUP signature that we have recently shown to be associated with colorectal tumors (Cameron et al., n.d.). On the other hand, the deletions and duplications involved in complex events have length distributions closely resembling that of inversions clustered in complex events. We also find that the per sample counts of deletions, duplications and inversions in complex events closely follows a 1:1:2 ratio as expected from random rearrangements following a catastrophic event[FIGURE 2C], whereas the counts of simple deletion and duplication junctions per sample were only very weakly positively correlated to those for deletions or duplication that are categorised as part of complex events (deletions r = 0.156, duplications r = 0.13)[FIGURE 2D]. Taken together these observations suggest LINX is able to accurately distinguish between simple and complex rearrangements.

LINX also annotates every cluster involving two directly related break junctions (further referred to as 2-break junction events) with a resolved type where they can be consistently chained [EXT FIGURE 2] or marks as ‘incomplete’ where they cannot form a consistent set of derivative chromosomes [EXT FIGURE 3]. Consistent 2-break junction clusters fall into two major categories - reciprocal events (eg. reciprocal inversions or translocations) or events with insertions of a templated sequence either in a chain or cycle (Li et al. 2020). 2-break junction events with insertion sequences most frequently involve very short templated sequences <1kb in length, previously referred to as ‘genomic shards’ (Bignell et al. 2007), which we find to be pervasive in cancer constituting 14% of somatic breakpoints. Genomic shards can confound classification of otherwise simple variant types, since a short templated insertion from another chromosome appears notionally as 2 translocations and can easily be misinterpreted as a reciprocal translocation or more complex event. LINX classifies events that can be resolved as a simple deletion, tandem duplication or translocation event with one or more inserted shards as a ‘synthetic’ event, under the assumption that the structure is most likely created by the disruption of a simple event with the insertion of the templated sequence during repair without affecting the donor locus. In support of this hypothesis, we find that samples with high counts of simple deletion and duplications have significantly higher (p<1*10^−60^ for both) counts of synthetic deletion and duplications respectively [EXT FIGURE 4A-B], and furthermore we observe the lengths of synthetic deletion and duplications are highly consistent with the respective lengths of simple deletions and duplications [EXT FIGURE 4C]. Synthetic deletion and duplication events can have many different breakend topological rearrangements depending on the source and orientation of the inserted shard [EXT FIGURE 1]. Insertion of genomic shards is by no means unique to simple deletion and duplication events, as we also see frequent short templated insertion sequences in breaks of more complex events including foldback inversion and chromothripsis events. Synthetic foldback inversions also show the same length distribution as simple foldbacks [EXT FIGURE 4C].

Reciprocal events are the other major category of 2-break junction events formed from 2 concurrent double stranded breaks, either a pair of inversions if both breaks occur on the same chromosome or a pair of translocations when the breaks occur on different chromosomes. Reciprocal inversions and translocations are well known rearrangements but are present in 65% of samples in the HMF cohort, but are infrequent relative to other events in cancer making up 0.8% and 0.5% of all clusters, respectively. In addition to these classical reciprocal events we also find other configurations of reciprocal events that likely involved 2 breaks and were found in 40% of samples [RECIP_TRANS_DUPS and RECIP_INV_DUPS in EXT FIGURE 2]. One prominent configuration we term ‘reciprocal duplication’ and involves a pair of reciprocal translocations or inversions between 2 distant loci but with breakends facing each other at each end often with multiple kilobase or even megabase overlap [EXT FIGURE 4D]. Reciprocal duplications are significantly enriched (p<1*10^−60^) in samples with strong tandem duplication signatures [EXT FIGURE 4E]. Moreover the length distribution of reciprocal duplications matches the length distribution of the signature for samples with drivers known to cause tandem duplication phenotypes, ie. *BRCA1*, *CCNE1* or *CDK12* drivers [EXT FIGURE 4F]. This suggests that these reciprocal duplication events may arise from the same process that forms tandem duplication events, likely when multiple tandem duplications occur simultaneously in a cell and instead of repairing locally that they may crossover and create a reciprocal duplication. This observation places constraints on the mechanism by which tandem duplications may form, since it requires duplication of DNA at both loci prior to breakage, and is consistent with a replication restart-bypass model (Willis et al. 2017) but not microhomology-mediated break-induced replication model (Hastings, Ira, and Lupski 2009).

#### Mobile element and pseudogene insertion detection

Somatic integration of long interspersed nuclear elements (LINE) are common features in many types of cancer particularly esophagus and head and neck cancers (Rodriguez-Martin et al. 2020). A LINE insertion may involve either the transposition of a full or partial LINE source element or the transduction of a partnered or orphaned genomic region within 5kb downstream of the LINE element. Whilst LINE insertions are typically simple events in themselves, correct classification of these break junctions is important to accurate interpretation of the genome as they can otherwise be mistaken as translocations and other complex events.

LINE integrations can be difficult to resolve with short read technology, since the inserted sequence is frequently not uniquely mappable in the genome and typically includes a Poly-A tail (Tubio et al. 2014) making assembly difficult. LINX circumvents both these issues by leveraging GRIDSS’s single breakend calling capability (Cameron et al., n.d.) to identify LINE insertion sites with breakend evidence for either repetitive LINE sequence, PolyA sequence or a list of known recurrently active LINE source elements. To validate LINX’s detection of mobile element insertions we ran LINX on 75 samples from the PCAWG pan-cancer cohort and compared LINX’s LINE insertion calls to those from traffic-mem (Rodriguez-Martin et al. 2020). Overall, 339 of 564 (60%) LINX LINE insertions calls were also detected by traffic-mem, with traffic-mem calling an additional 270 insertions not found by LINX. The concordance in total LINE insertion count was very strong on a per sample basis [EXT FIGURE 5A,SUPP TABLE 4] with most of the private calls in both pipelines being found in the high LINE mutational burden samples [EXT FIGURE 5B] suggesting that many of the private calls from both pipelines may be genuine LINE insertions.

Across the full HMF cohort, LINX found 76% of tumors have at least 1 LINE insertion event. Some tumors suffer extreme deregulation, with 6.7% of tumors having over 100 insertions and 2241 insertions found in a single Esophagus tumor sample [FIGURE 3A,3B]. The most frequently inserted LINE source elements in the Hartwig cohort were all amongst the top 6 reported previously in the PCAWG pan-cancer cohort (Rodriguez-Martin et al. 2020): chr22:29,059,272-29,065,304, chrX:11,725,366-11,731,400, chr14:59,220,385-59,220,402,chr9:115,560,408-115,566,440 and chr6:29,920,213-29,920,223. Analysis of the precise breakend locations at these sites reveals highly recurrent site specific patterns of transduction [FIGURE 3C, EXT FIGURE 5C], where the 3’ ends of the transduced sequences are normally sourced from a handful of specific downstream sites (presumably polyadenylation sequences of alternative transcription endpoints for the LINE source element) whereas the location of 5’ side of the transduction appears to be relatively randomly distributed.

**FIGURE 3:**
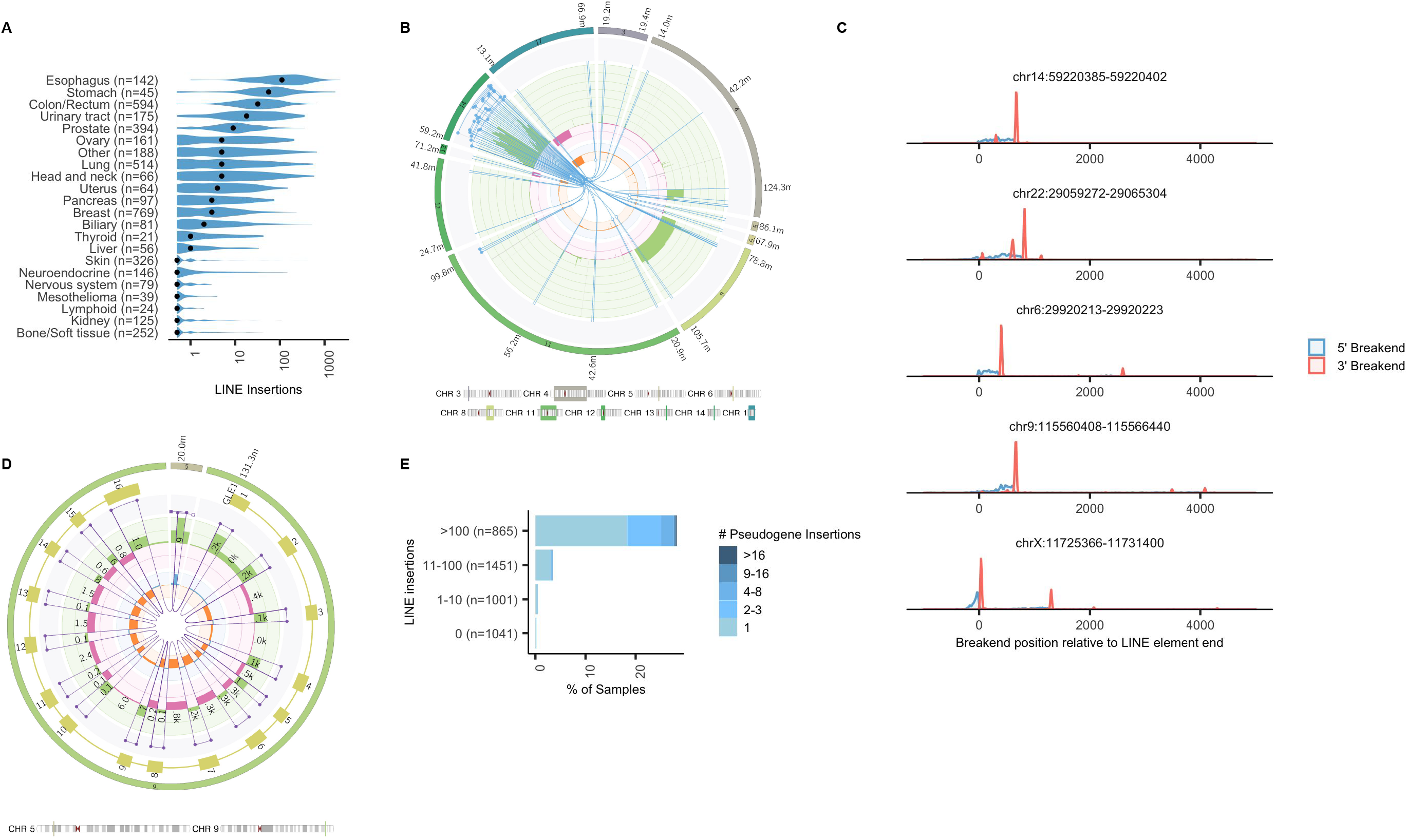
Mobile element insertions a) Violin plot showing the distribution of the number of LINE insertions per sample grouped by tumour type. Black dots indicate the median values for each tumour type b) Complex LINE cluster in HMF002232B, a colorectal cancer. Overlapping segments from the LINE source element from chr 14:59.2M has been inserted in at least 20 independent locations scattered throughout the genome. c) Histogram showing frequency breakends positions for all mobile element transductions in HMF cohort originating from the 5 most active LINE source elements relative to the last base of the LINE source element. d) Pseudogene insertion of *GLE1* into an overlapping break junction on chromosome 5 in HMF002165A, a non-small cell lung cancer. All 16 exons of the *GLE1* canonical transcript are inserted, but part of the first and last exons are lost. e) Samples with high numbers of LINE insertions also have high numbers of pseudogene insertions

At the LINE insertion site, accurate breakpoint determination can also give insight into potential biological mechanisms. LINX finds frequent target site duplication (Ostertag and Kazazian 2001), but intriguingly finds 2 peaks in the distance between the insertion breakends, one at an overlap of 16 bases, but also a 2nd peak with no overlap suggesting the possibility of two distinct breakage mechanisms for the 2nd strand after LINE invasion [EXT FIGURE 5D]. Furthermore for the 20% of insertions where LINX observed a 5’ inversion in the insertion sequence (due to twin priming (Ostertag and Kazazian 2001), only a single peak with target site duplication of 16 bases is found.

LINX also detects somatic pseudogene insertion resulting from the activated reverse transcriptase activity associated with deregulated LINE activity in tumors (Rodriguez-Martin et al. 2020). LINX annotates any group of deletions that matches the exact boundaries of annotated introns as pseudogene insertions [FIGURE 3D]. We find 577 pseudogene insertions in our cohort, exclusively in samples with somatically activated LINE mechanisms and enriched in the samples with the most deregulated LINE activity [FIGURE 3E].

#### Complex events

LINX classifies any cluster which has 3 or more junctions and is not resolved as a LINE insertion as ‘COMPLEX’. Previous tools, notably ChainFinder ((Baca et al. 2013), have been developed to systematically search for complex rearrangement patterns in tumors. We compared LINX and ChainFinder across 1479 samples and found that whilst in 22% of cases LINX and ChainFinder produced near identical clusters, the majority of junctions clustered by LINX are left unclustered by ChainFinder, while few SVs were exclusively clustered by ChainFinder [EXT FIGURE 6A,6B]. We find this to be due to two main reasons: first, ChainFinder fails to cluster a large number of junctions that are highly proximate (<5kb between breakends) [EXT FIGURE 6C] and second, LINX employs a variety of clustering techniques to link distant junctions on the same chromosome arm which are not captured by ChainFinder [EXT FIGURE 6D]. The additional variants clustered by LINX compared to ChainFinder share a strikingly similar length distribution to the variants clustered by both tools [EXT FIGURE 6E], including deletions, duplications and inversions with lengths greater than 1Mb which are not normally found in simple events.

Conversely, in a small proportion (1.8%) of cases, junctions are clustered by ChainFinder and not by LINX. 95% of these are deletions and tandem duplications <1Mb in length which may also have occurred as independent events and be inadvertently clustered by ChainFinder [EXT FIGURE 6E]. In line with this, we find that 20% of the deletions clustered by ChainFinder but not by LINX are in known fragile sites [EXT FIGURE 6F], and often are phased in trans suggesting that they likely occurred in different events (Hadi et al. 2020).

Across the HMF cohort we found at least 1 COMPLEX event in 95% of tumors, and at least 1 event of 20 or more junctions in 60% of tumors [FIGURE 4A]. Whilst there are relatively few complex events in any given tumor, they account for more than half of junctions overall. Complex clusters with >100 junctions were found in all cancer types, with breast cancer having the highest median maximum cluster size of 62 [FIGURE 4B]. We observe that complex events with a higher number of junctions are more likely to disrupt or amplify a putative cancer driver gene. Overall, 12.7% of all complex clusters in the cohort contributed to a LOH, amplification deletion or disruption driver, but this rises to 39.1% for events with 20 or more junctions and 77% for events with more than 20 junctions and high amplification (JCN >=8) [FIGURE 4C]

**FIGURE 4:**
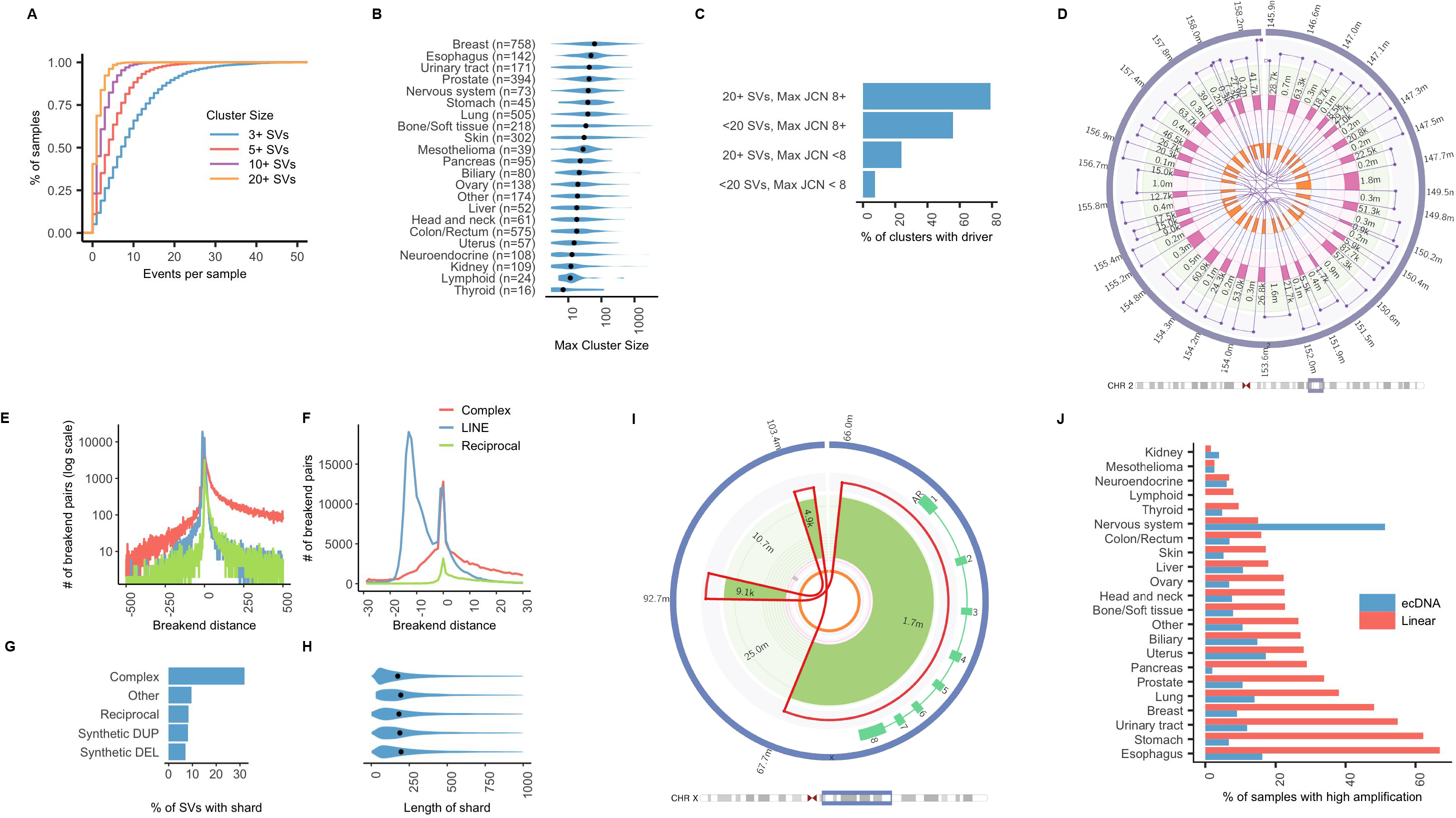
Complex rearrangements and high amplification a) Cumulative distribution function plot of count of complex rearrangement clusters per sample with at least 3,5,10 and 20 variants b) Violin plot showing the distribution of the maximum number of variants in a single complex rearrangement cluster per sample grouped by tumour type. Black dots indicate the median values for each tumour type c) Proportion of clusters contributing to at least 1 amplification, deletion, homozygous disruption or LOH driver in a panel of cancer genes by complexity of cluster and maximum JCN d) Fully resolved chromothripsis event consisting of 31 structural variants affecting a 13mb region of chromosome 2 in HMF001571A, a prostate tumor. e) &f) Counts of occurrences of trans-phased breakends by distance between the breakends for complex events, LINE insertions and 2-break reciprocal clusters in the range of −500 to 500 bases (log scale) and zoomed in to −30 to 30 bases respectively. Negative distances indicate overlapping breakends and duplication at the rearrangement site. g) Proportion of variants with at least one breakend joining a shard of less than 1kb in length by resolved type for selected resolved types. h) Violin plot showing the distribution of shard length by resolved type. i) Double minute formed by 3 junctions in HMF003969A, a prostate tumor, and which amplifies known oncogene, *AR*, to a copy number of approximately 23x. j) Proportion of samples with ecDNA and linear amplifications by cancer type

LINX goes further than other clustering tools in that it allows not only for complex clusters to be identified, but in many of cases is able to completely resolve such events into a consistent set of derivative chromosomes, including in chains with up to 33 or more junctions [FIGURE 4D]. Uniquely, and critically for accurate chaining in these complex structures, LINX utilises phased assembly output from GRIDSS to determine whether proximate facing breakends are cis or trans-phased(Cameron et al., n.d.). Trans-phased facing breakends, causing local duplication, are common in complex events and can often extend up to several hundred bases, but only rarely extend beyond 30 bases in reciprocal events and mobile insertions [FIGURE 4E,4F], suggesting a fundamentally different breaking mechanism in complex events which may cause double stranded breaks with significant overlap. Proximate cis-phased breakends are even more common than trans-phased accounting for 16.4% of all junctions in complex events, and resemble in length distribution the shards detected in simple events but with higher frequency in complex clusters [FIGURE 4G-H]. We frequently observe localised regions of scarring with multiple distinct shards sourced from the same location sometimes with overlapping templates sequences.

#### Amplification mechanisms

Regions of high amplification are amongst the most complex events in tumors as they require iterative and repeated cycles of synthesis or unequal segregation to form. Two key distinct biological mechanisms are well known which create highly amplified rearrangements: repeated cycles of breakage fusion bridge (BFB) and stochastic amplification of circular extrachromosomal DNA by asymmetric segregation during cell division (ecDNA). ecDNA may arise from any event which creates simultaneous multiple double stranded breaks on the same chromosomal arm with one or more chromosomal segments repairing to form a circular structure without a centromere. BFB, on the other hand, is triggered by the formation via translocation or foldback inversion of a chromosome with 2 centromeres, arising from either multiple concurrent double stranded breaks or telomere erosion, and leads to duplication of chromosomal segments within a linear chromosome.

Despite these significant differences in mechanism, distinguishing between ecDNA and BFB is non trivial based on short-read sequencing data, but is essential in order to understand the diversity of amplification drivers in tumors and may be relevant to the prognosis or treatment of certain tumors (Kim et al. 2020). The key difficulties in discrimination are that both mechanisms can leave a similar footprint as both may arise out of complex shattering events, and are highly shaped by the same selection processes, both positive (amplification of key oncogenes) and negative (constraints on amplifications of other proximate genes).

LINX employs a set of heuristics to identify subsets of clusters as likely ecDNA [FIGURE 4,EXT FIGURE 7A]. The key principle used to identify ecDNA is to look for high junction copy number (JCN) structural variants adjacent to low copy number regions which can be chained into a closed or predominantly closed loop. LINX also checks that the high JCN cannot be explained by compounding linear amplification mechanisms, by comparing the JCN of the candidate ecDNA junctions to the maximal amplification impact of foldback inversions (hallmarks of BFB) as well as junctions that link closed segments of the ecDNA to other regions of the genome and may cause part of the amplification (see methods). We validated the ecDNA calling on a set of 13 WGS neurosphere cultured glioblastoma samples that had been previously analysed (deCarvalho et al. 2018) for ecDNA with Amplicon Architect (Deshpande et al. 2019). LINX and Amplicon Architect called ecDNA for an identical set of 19 oncogenes across the 13 samples [SUPP TABLE 5] including the 11 samples which were orthogonally validated by FISH.

Applying the ecDNA heuristic to the HMF cohort, we find that ecDNA to be a relatively uncommon event present in 9.9% of all tumors, with the highest frequency in CNS tumors (51%) [FIGURE 4J]. This is lower than found in a large recent pan-cancer cohort analysis of WGS using AmpliconArchitect (Kim et al. 2020) which found a pan-cancer prevalence 14%. We find that overall 12% of putative amplification drivers identified in the HMF cohort are associated with ecDNA events [EXT FIGURE 7B], but that this rate increases for more highly amplified events to greater than 40% for events with maximum JCN>32. The relative rate of ecDNA is the highest for *EGFR* [EXT FIGURE 7C] but this appears to be highly specific to CNS tumors (where 87% of *EGFR* amplifications are associated with ecDNA) whereas for lung tumors (where EGFR amplification is also common) and other cancer types the rates of ecDNA are only 11% and 21% respectively similar to that of other well known oncogenes [EXT FIGURE 7D].

The high amplification events that do not meet the ecDNA criteria are assumed to be formed via linear amplification. Whilst we find that 76% of these events have at least 1 foldback inversion which may suggest a breakage fusion bridge process [EXT FIGURE 7E], in many events the foldback JCN does not explain the full amplification and in the remaining events LINX identifies no foldback events at all [EXT FIGURE 7F]. The majority of these are unlikely to be ecDNA, however, since there is no obvious set of junctions and segments which can be closed into a circle with a consistent copy number. Some events, such as the exceptionally complex amplifications of *MDM2* & *CDK4* common in liposarcoma (Garsed et al. 2014) may not fall neatly into either an ecDNA or breakage fusion bridge classification [EXT FIGURE 7G].

### Detection of clinically relevant pathogenic rearrangements

LINX calls a diverse and comprehensive range of fusions and pathogenic rearrangements [FIGURE 5A, EXT FIGURE 8 A-D]. We orthogonally validated LINX’s pathogenic fusion predictions, by comparing them to fusions predicted from RNA-seq data taken from the same samples. For the RNA comparison, we used Arriba, one of the best performing RNA fusion callers (Haas et al. 2019).

**FIGURE 5:**
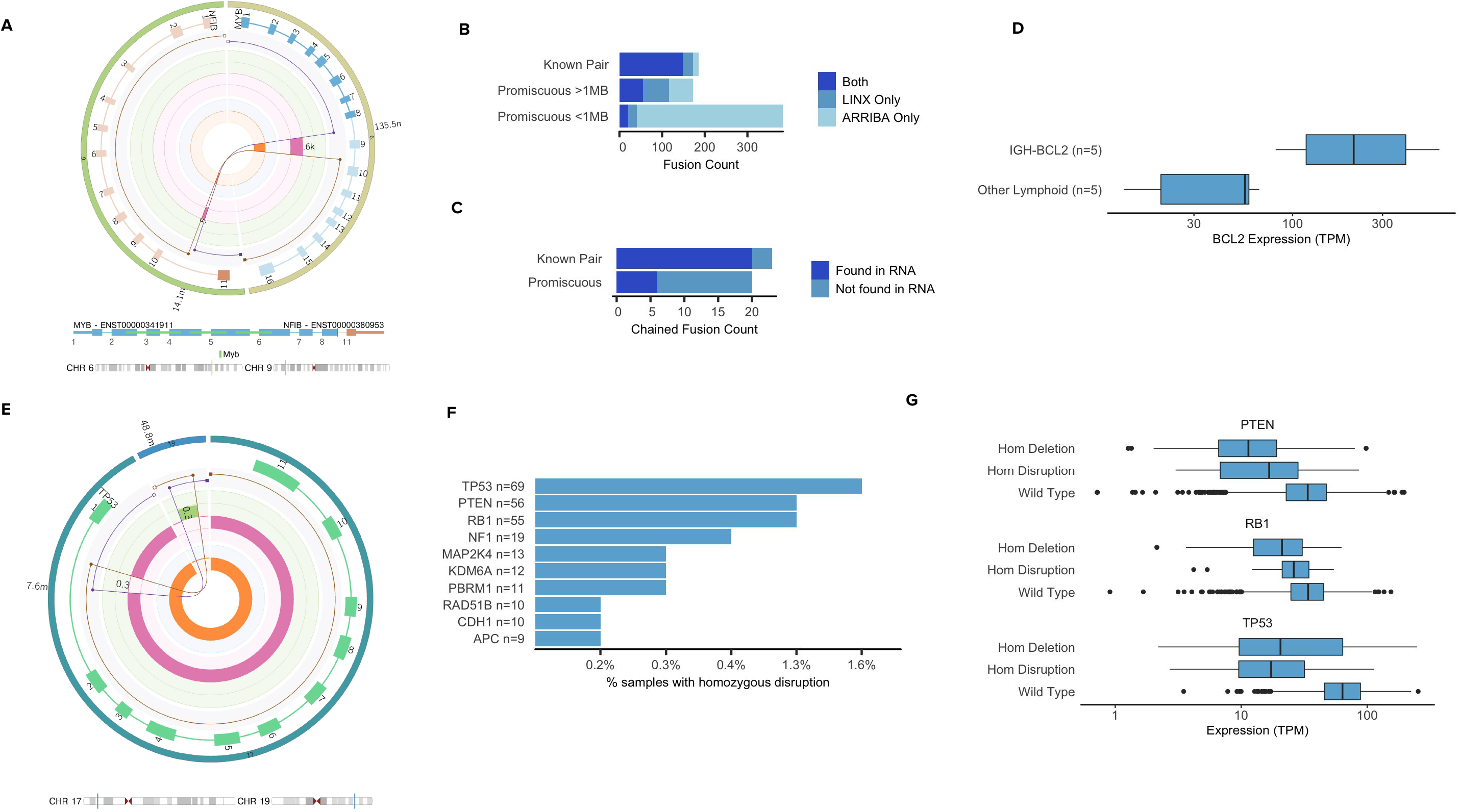
Clinically relevant rearrangements a) A *MYB*-*NFIB* fusion caused by a reciprocal translocation in HMF000780A, a salivary gland tumor. The translocation links exons 1-8 in MYB to exon 11 in NFIB. b) Comparison of LINX fusion predictions in HMF cohort to Arriba fusion predictions from orthogonal RNA sequencing for known pairs and promiscuous fusion partners. Promiscuous fusions of less than 1Mb length are shown separately as they may occur from read through transcripts and not be associated with a genomic rearrangement. c) Count of LINX chained fusion predictions for known and promiscuous fusion and whether they are also found to be expressed in RNA by Arriba d) Distribution of *BCL2* expression in Lymphoid samples with and without a predicted pathogenic *IGH-BCL2* rearrangement. box: 25th-75th percentile; whiskers: data within 1.5 times the IQR. P-values indicate significance from a two-tailed Mann–Whitney U-test. e) Reciprocal translocation affecting *TP53* in HMF001913A, a prostate tumor. The 2 predicted derivative chromosomes overlap by approximately 300 bases on both ends, but are trans-phased which rules out the possibility of a templated insertion at either location. Although the *TP53* copy number alternates between 1 and 2, no derivative chromosome contains the full gene and the gene is homozygously disrupted. f) Prevalence of homozygous disruption drivers for top 10 most affected tumor suppressor genes g) Distribution of gene expression in HMF cohort for samples with homozygous deletion, homozygous disruption and wild type for each of *RB1*, *TP53* and *PTEN*. box: 25th-75^th^ percentile; whiskers: data within 1.5 times the IQR. P-values indicate significance from a two-tailed Mann–Whitney U-test.

From 1924 HMF cohort samples with matched RNA and using a curated list of 391 known pathogenic fusion pairs [SUPP TABLE 6], 148/173 inframe fusions (86%) predicted by LINX were also found by Arriba [FIGURE 5B, SUPP TABLE 7]. Of the 25 fusions not identified in RNA, 13 matched the characteristic tumor type of the known fusion pair (9 of which were *TMPRSS2-ERG* fusions in prostate cancer), and are likely to be pathogenic, but with insufficient expression to be detected in the RNA. A further 2 cases predicted by LINX were found by Arriba but only in out of frame transcripts. 13 known pair fusions were predicted by Arriba but not by LINX, 7 of which involve gene pairs less than 1 million bases apart on the same chromosome, and may be caused by readthrough transcripts(He et al. 2018) or circularised RNA(Yu and Kuo 2019) unrelated to structural rearrangements in the DNA.

63 cancer related fusion genes were curated as promiscuous 5’ and 3’ partners. Among these, LINX identified a further 152 candidate inframe fusions, 74 (49%) of which were also detected in RNA. Arriba detected 397 additional promiscuous candidates, but 86% of these were proximate on the same chromosome and are likely read through transcripts with no genomic rearrangement. Altogether, 43 of the 325 (13%) known and promiscuous fusion predictions were chained fusions involving multiple junctions, 26 (60%) of which were validated in the RNA-seq data [FIGURE 5C], highlighting the utility of chaining of derivative chromosomes for DNA fusion calling. TMPRSS2-ERG was the only fusion which LINX found to be recurrently chained in the cohort accounting for 14 of the 43 predicted chained fusions, all in prostate cancers.

Immunoglobulin enhancer rearrangements are a distinct class of pathogenic rearrangements, common in B-cell tumors where errors in VDJ recombination and/or isoform switching in the *IGH*,*IGK* and *IGL* regions may lead to pathogenic rearrangements driving high expression of known oncogenes through regulatory element repositioning(Chong et al. 2018). Although these typically do not make a novel protein fusion product, LINX predicts these pathogenic rearrangements based on the breakend in the *IGH*, *IGK* and *IGL* regions with orientation and position matching locations commonly observed in B-cell tumors (Chong et al. 2018). Amongst 10 lymphoid samples with matching RNA in the cohort, LINX found 6 such rearrangements including 5 cases of *IGH-BCL2* and 1 case of *IGH-MYC*. The 5 identified samples with *IGH-BCL2* rearrangements have significantly higher expression (p=0.008) of *BCL2* than the 5 lymphoid samples with no *BCL2* rearrangement detected [FIGURE 5D].

LINX also identifies disruptive intragenic rearrangements which may cause exonic deletions and duplications. Our knowledge base includes 9 such rearrangements known to be pathogenic and 2 which we have deemed likely pathogenic due to high recurrence in the HMF cohort. Three of these pathogenic exon rearrangements were detected by LINX in at least 5 samples with paired RNA in our cohort: *EGFRvII* (n=6), *EGFRvIII*(n=14), *CTNNB1* Exon 3 deletion (n=6). In all cases with an event detected by LINX in the DNA, we found RNA fragments which supported novel splice junctions in the matched RNA [EXT FIGURE 8E]. Only 1 other sample in the complete cohort (n = 1924) with had more than 1 fragment supporting any of these alternative splice junctions (a gastrointestinal stromal tumor with 3 fragments supporting EGFRvII but with no evidence of rearrangement in EGFR).

In addition to producing novel oncogenic proteins and over expression of well known oncogenes, rearrangements may also lead to tumorigenesis by disrupting the function of tumor suppressor genes. To capture this LINX annotates every breakend that overlaps a gene, determines whether it is disruptive to the gene and reports the number of undisrupted copies. In cases of reciprocal translocation [FIGURE 5E], reciprocal inversion [EXT FIGURE 9A], complex break event [EXT FIGURE 9B] or a tandem duplication which overlaps at least one exon [EXT FIGURE 9C], a gene may be disrupted on all remaining copies even though the copy number is greater than 0 for all exonic bases (Patch et al. 2015). We term this type of genomic rearrangement to be a ‘homozygous disruption’. We find homozygous disruptions to be a common driver in the HMF cohort with 9% of samples containing at least 1 homozygous disruption in a panel of 448 curated cancer related genes [SUPP TABLE 8]. Three well known tumor suppressor genes had homozygous disruptions in more than 1% of the cohort: *TP53* (n=69), *PTEN* (n=56), and *RB1*(n=55) [FIGURE 5F]. We found significantly lower expression for each of these genes (p=2*10^−16^ TP53;p=2*10^−6^ PTEN;p=2*10^−3^ RB1)in the samples where they were homozygously disrupted compared to samples with at least one intact copy [FIGURE 5G]. Homozygous disruptions cannot readily be detected by standard panel or whole exome sequencing, since intronic sequences are typically not included in such panels and they are exonic copy number neutral events.

#### Novel visualisation

LINX produces novel detailed visualisations of the rearrangements in the tumor genome which allow further insights into complex rearrangements. LINX supports either drawing all rearrangements in a cluster or all the rearrangements on a chromosome, creating an integrated Circos output (Krzywinski et al. 2009) showing copy number changes, clustered SVs, the derivative chromosome predictions and impacted genes including protein domain annotation for gene fusions all on the same diagram. The visualisations use a log-based position scaling between events so that small and large scale structures can both be inspected on a single chart. Combined with the circular representation, these features allow unprecedented resolution of complex structures across a broad array of event types including chromoplexy [FIGURE 6A] and complex BFB amplification events [FIGURE 6B]. Supplementary note 1 provides a walk through and explanation of all LINX figures covering the complete SV landscape of the COLO829T melanoma cancer cell line which has been proposed as a somatic reference standard for cancer genome sequencing(Craig et al. 2016; Valle-Inclan et al., n.d.)

**FIGURE 6:**
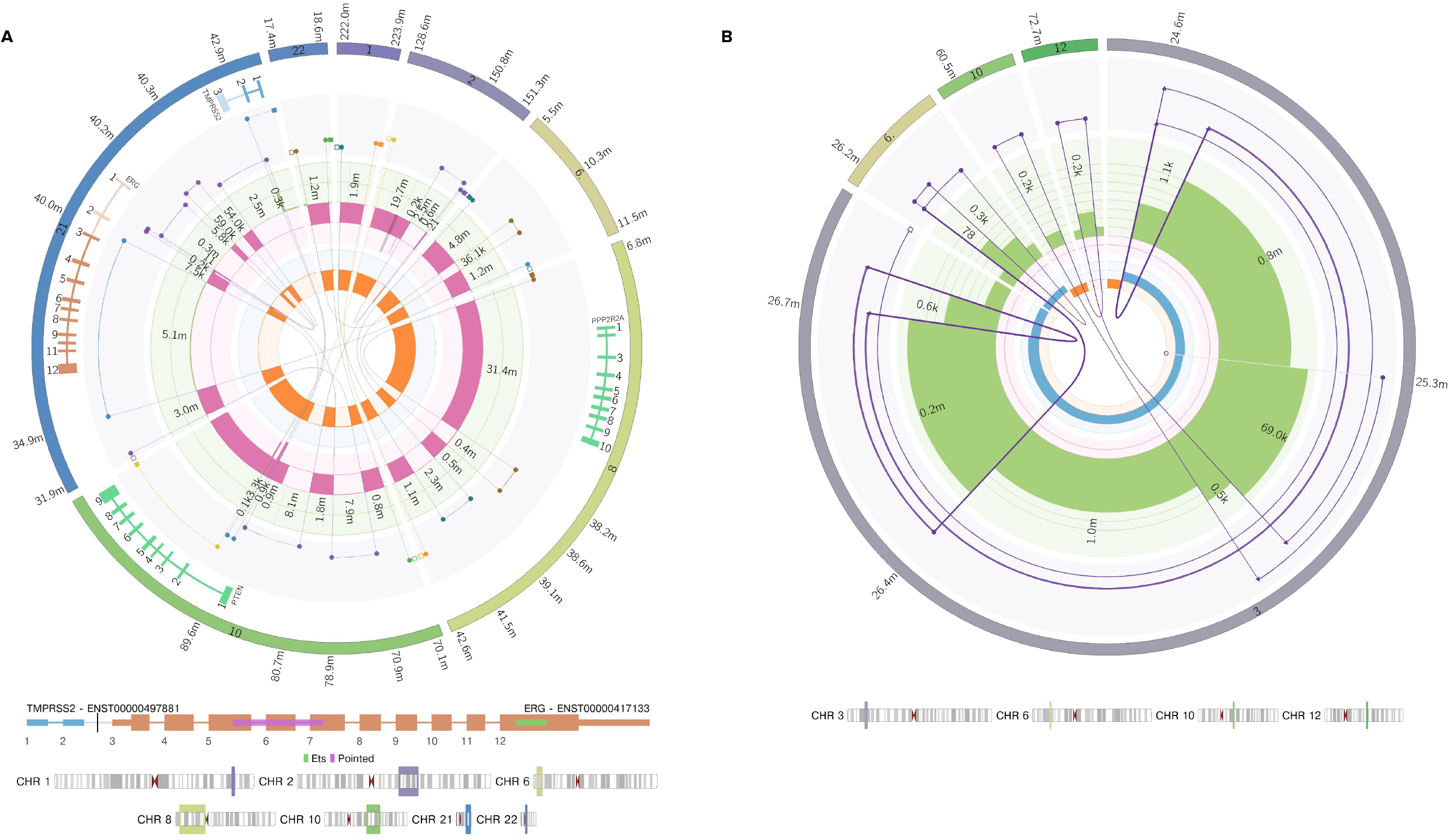
Complex event visualisation a) Chromoplexy-like cluster formed from 19 break junctions across 7 chromosomes in HMF001596B, a prostate tumor. The rearrangement leads 3 distinct putative drivers in a single event, including a chained *TMPRSS2-ERG* fusion with 2 hops, a loss of heterozygosity for *PPP2R2A* which also has a stop gained point mutation (not shown), and an intronic homozygous disruption of *PTEN*. b) Breakage fusion bridge event affecting the P arm of chromosome 3 in the melanoma cell line COLO829T. The predicted derivative chromosome has a copy number of 2 and can be traced outwards starting from the centromere on chromosome 3, traversing 2 simple foldbacks and 2 chained foldbacks and finishing on a single breakend at chr3:25.3M which from the insert sequence can be inferred to be connected a centromeric satellite region (likely chromosome 1 which has a copy number gain of 2 over the centromere from P to Q arm, and which appears to be connected to chromosome 3 in unpublished SKY karyotype figures http://www.pawefish.path.cam.ac.uk/OtherCellLineDescriptions/COLO829.html). One chained foldback at chr3:26.4M includes a genomic shard from chr 6 of approximately 400 bases which has itself been replicated and internally disrupted by the foldback event. The other chained foldback at chr3:25.4M includes 2 consecutive genomic shards inserted from chromosome 10 and 12 of approximately 200 bases each.

## Discussion

We have shown that LINX can help in multiple ways to understand highly rearranged cancer genomes. Linx does however have several limitations. Due to the complexity of dissection of genomic rearrangements through short-read paired-end whole genome sequencing, there are many potential sources of error which can confound correct interpretation of these genomic rearrangements including sample preparation, sequencing errors and biases (such as GC bias), inaccurate fitting of sample purity and ploidy, false positive or false negative structural variant calls, and inaccurate local copy number measurement. High quality deep sequencing tumor-normal is also essential (106x tumor / 38x normal with 500-600bp median fragment length was used in the HMF cohort). Lower coverages are likely to result in less complete reconstructions. Furthermore whilst LINX has been optimised for short read technology the short read length is ultimately the key limitation since it limits the phasing of proximate variants and accurate identification of events in long repetitive regions. Nevertheless, in practice, LINX is able to resolve many structures via various chaining and clustering heuristics, but for more complex events, particularly highly rearranged focal regions, errors are inevitable and the chaining is only partial and representative. Long read sequencing technology (Sedlazeck et al. 2018)may be better suited for resolving such events, although those technologies typically perform less well for small variant detection, which is also key for comprehensive cancer genome characterisation.

The challenges in understanding the complexity of rearrangements in tumor genomes can be daunting. The diversity of overlapping or converging biological mechanisms which may cause similar rearrangement patterns, means that it may be perilous to analyse any one rearrangement as a stand alone analysis. By exhaustively classifying all rearrangements, LINX is a robust foundation for more detailed analysis of specific rearrangement patterns, and can provide a basis for further in depth analysis particularly of structural variant signatures, complex shattering events and high amplification drivers as well as dissection of underlying molecular mechanisms and DNA replication and repair components involved. The full LINX analysis results on the HMF cohort are also available via data request and can be paired with clinical data and other whole genome analyses for further indepth research.

WGS offers the promise of a single comprehensive test for all genomic alterations for both routine diagnostics and future biomarker discovery. LINX takes a step towards that goal by both comprehensively calling clinically relevant fusions from DNA with similar precision and sensitivity to gold standard RNA-seq methods, and by identifying homozygous disruptions, an important class of drivers of tumorigenesis that cannot readily be detected by standard of care methods.

## Methods

LINX v1.12 was used for all analyses in this paper and is described in detail in the supplementary information.

### Patient cohort and WGS pipeline

The patient cohort was derived from the Hartwig Medical Foundation Cohort for which the sample collection and whole genome sequencing and alignment to the GRCH37 reference genome has previously been described (Priestley et al. 2019). We filtered for the highest purity sample from each patient from tumor samples with purity >= 20% and with no QC warnings or fails yielding 4,378 paired tumor-normal whole genome samples in total. GRIDSS (Cameron et al., n.d.) v2.93 and PURPLE (Priestley et al. 2019) v2.48 were used for copy number and structural variant inputs for LINX.

### RNA validation

For 1,860 samples paired whole transcriptome sequence data was also used. The RNA-seq was aligned to the GRCH37 genome using STAR 2.7.3a. Gene expression was calculated using Isofox v1.0, which uses an expectation maximisation algorithm to estimate transcript abundance from genome aligned RNA-seq data, with default parameters. Isofox was also used to count the RNA fragments supporting novel splice junctions predicted in LINX for exon deletions and duplications. Isofox is described in detail at https://github.com/hartwigmedical/hmftools/tree/master/isofox.

Known pathogenic pair and promiscuous gene fusions predictions in the DNA were compared to passing fusion calls in the RNA by Arriba (https://github.com/suhrig/arriba). Fusions were considered to be matched if the gene pair matched between RNA and DNA

### Complex event validation

We compared LINX to ChainFinder v1.0.1 on 2840 samples from the HMF cohort. ChainFinder was run with default parameters. Both LINX and ChainFinder were run using the same GRIDSS/PURPLE input data. As ChainFinder hangs indefinitely on some samples, ChainFinder was run independently on each sample. Only samples for which ChainFinder completed within 24 hours were included in the comparison. ChainFinder clusters of 3 or more variants were considered equivalent to LINX’s COMPLEX classification. For each individual variant we determined whether it was clustered in LINX, in ChainFinder or in both as well as the size of the cluster in each tool.

### LINE insertion validation

We ran LINX on 75 WGS samples from the PCAWG cohort (supplementary table 2) which had previously been run with TraFiC-mem (Rodriguez-Martin et al. 2020). Insertions were considered matched between the tools if the predicted insertion site was within 50 bases.

### ecDNA validation

We ran LINX on 13 previously analysed (deCarvalho et al. 2018) WGS glioblastoma neurosphere cultures samples sequenced to ~10x depth and compared the ecDNA predictions of Linx to those of the AmpliconArchitect tool and FISH. We matched the ecDNA predictions by amplified oncogene per sample.

## Supporting information

Supplementary File 1 - Linx Methods

## Data availability

All raw (BAM), analysed (VCF, SV, purity copy number data) germline and somatic genomic data and LINX results from the HMF cohort were obtained from the Hartwig Medical Foundation (Data request DR-005). Standardized procedures and request forms for access to this data including LINX analysis results can be found at https://www.hartwigmedicalfoundation.nl/en.

## Code availability

LINX is freely available as open source software from the Hartwig Medical Foundation (https://github.com/hartwigmedical/hmftools/tree/master/sv-linx) under a GPLv3 license. Reference data required to run LINX on hg19 or hg38 is available from https://resources.hartwigmedicalfoundation.nl. Supplementary File 1 contains the PURPLE output files required to run LINX for the COLO829T cell line as an example. LINX can be run from raw paired tumor-normal FASTQ files as part of Hartwig’s open source cloud based cancer analysis pipeline (https://github.com/hartwigmedical/platinum). Alternatively a docker image is available from dockerhub as gridss/gridss-purple-linx to run GRIDSS, PURPLE and LINX together from tumor and normal BAMs.

## Acknowledgements

This publication and the underlying study have been made possible partly on the basis of the data that Hartwig Medical Foundation and the Center of Personalised Cancer Treatment (CPCT) have made available to the study. Whole-genome sequencing data were analysed from glioblastoma neurosphere lines, which were established at Henry Ford Healthcare System.

## Extended Figure Legends

**EXTENDED FIGURE 1:**
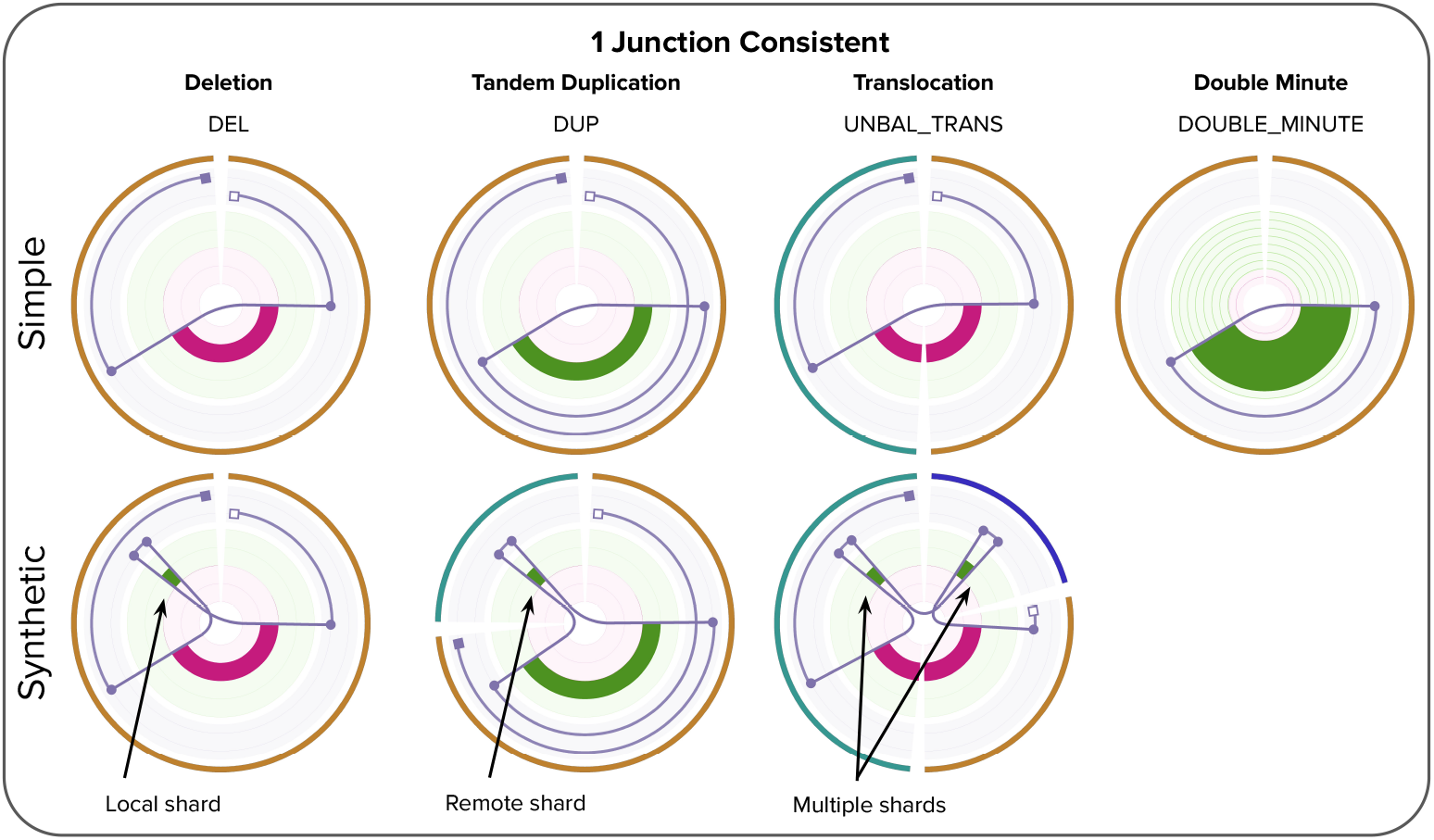
A catalog of consistent 1 junction structures in LINX. The upper row shows the 4 consistent chromosome structures that can be formed from a single junction including a simple deletion, tandem duplication, unbalanced translocation or single junction double minute. The lower 3 images show examples of equivalent multiple break junctions events that can be treated as equivalent synthetic versions of the above 1 junction events with a genomic shard (<1000 base segment of templated DNA) inserted.

**EXTENDED FIGURE 2:**
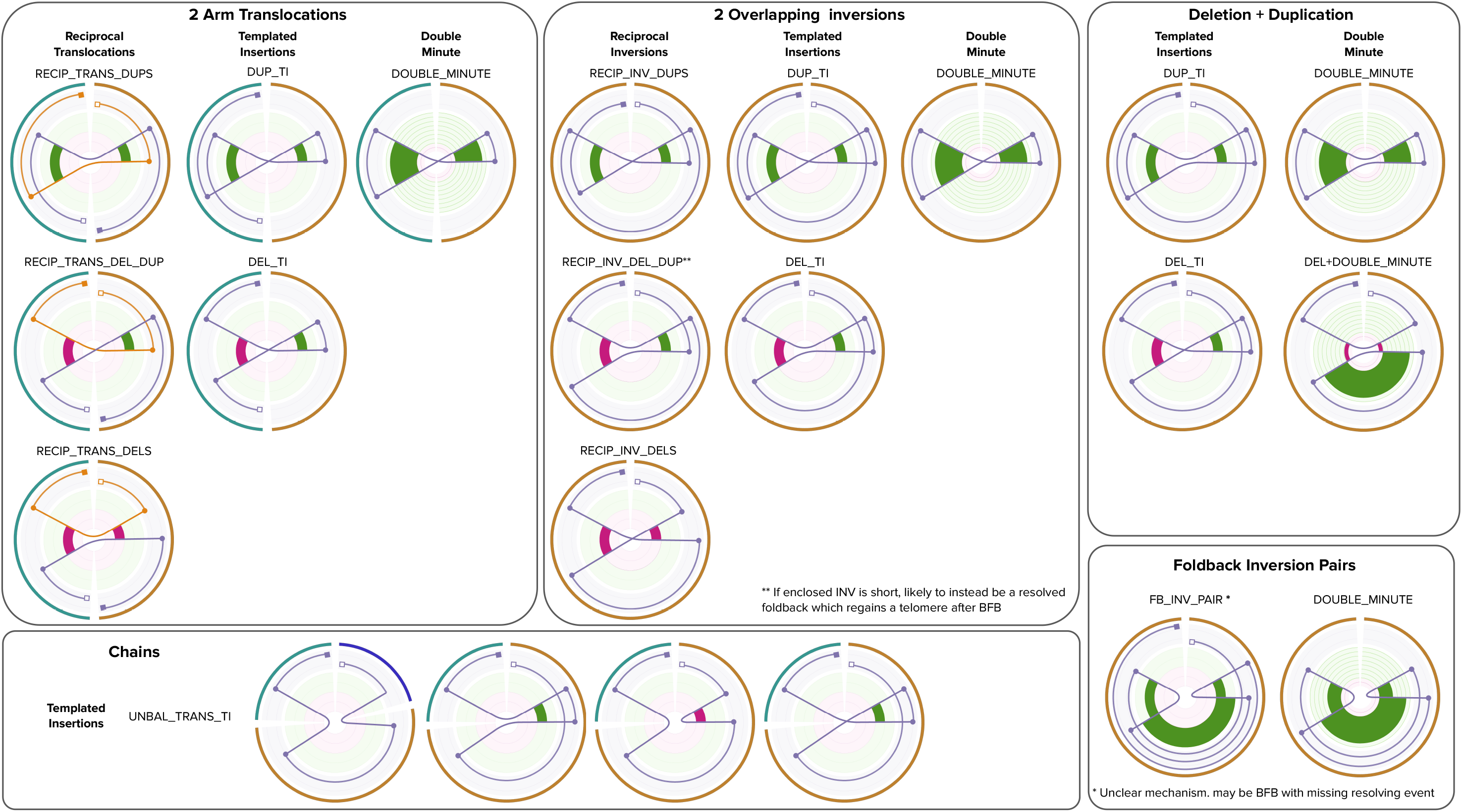
A catalog of consistent 2 junction rearrangements in LINX. The structures are organised by the geometry of the break junctions with the resultant derivative chromosomal structure dependent on both the orientation of the breakends and whether they are phased. Pairs of common arm translocations (top left panel) and overlapping inversions (top middle panel) can each form 3 different topologies of reciprocal events, 2 topologies of templated insertions and 1 double minute topology. Overlapping deletions and duplications may form templated insertion and double minutes, but not reciprocal structures (top right panel). Mixed combinations of translocations and local junctions (bottom left panel) create a chained translocation with templated insertion (classified as 2-break other in LINX), Non overlapping foldback inversions (bottom right panel) may form either a double minute or other linear amplification structure.

**EXTENDED FIGURE 3:**
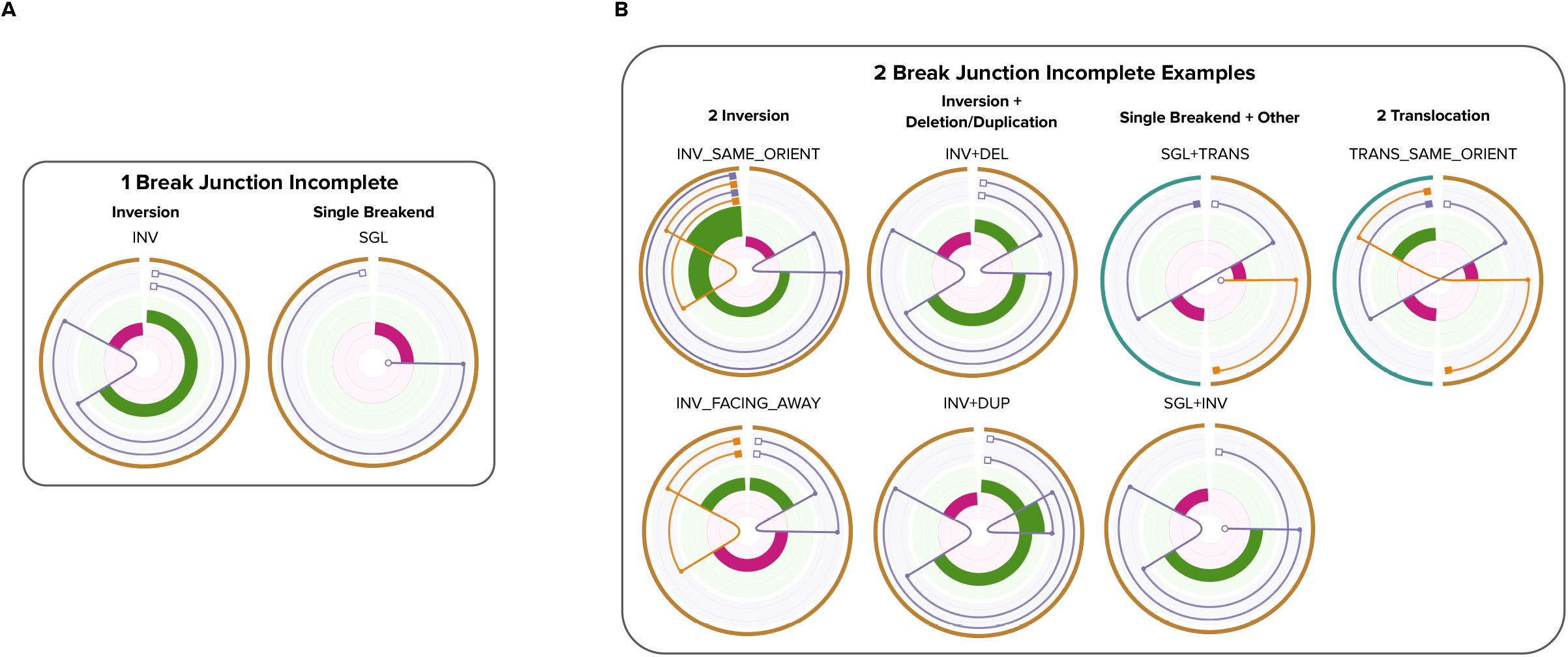
Examples of incomplete 1 and 2 junction rearrangements in LINX. A cluster is considered incomplete in LINX if it does not create a set of 1 or more consistent derivative chromosomes that each link a telomere and centromere. A rearrangement event linking 2 centromeres or 2 telomeres (without an intervening centromere) cannot guarantee equal division at mitosis and hence is likely to be inherently unstable.

**EXTENDED FIGURE 4:**
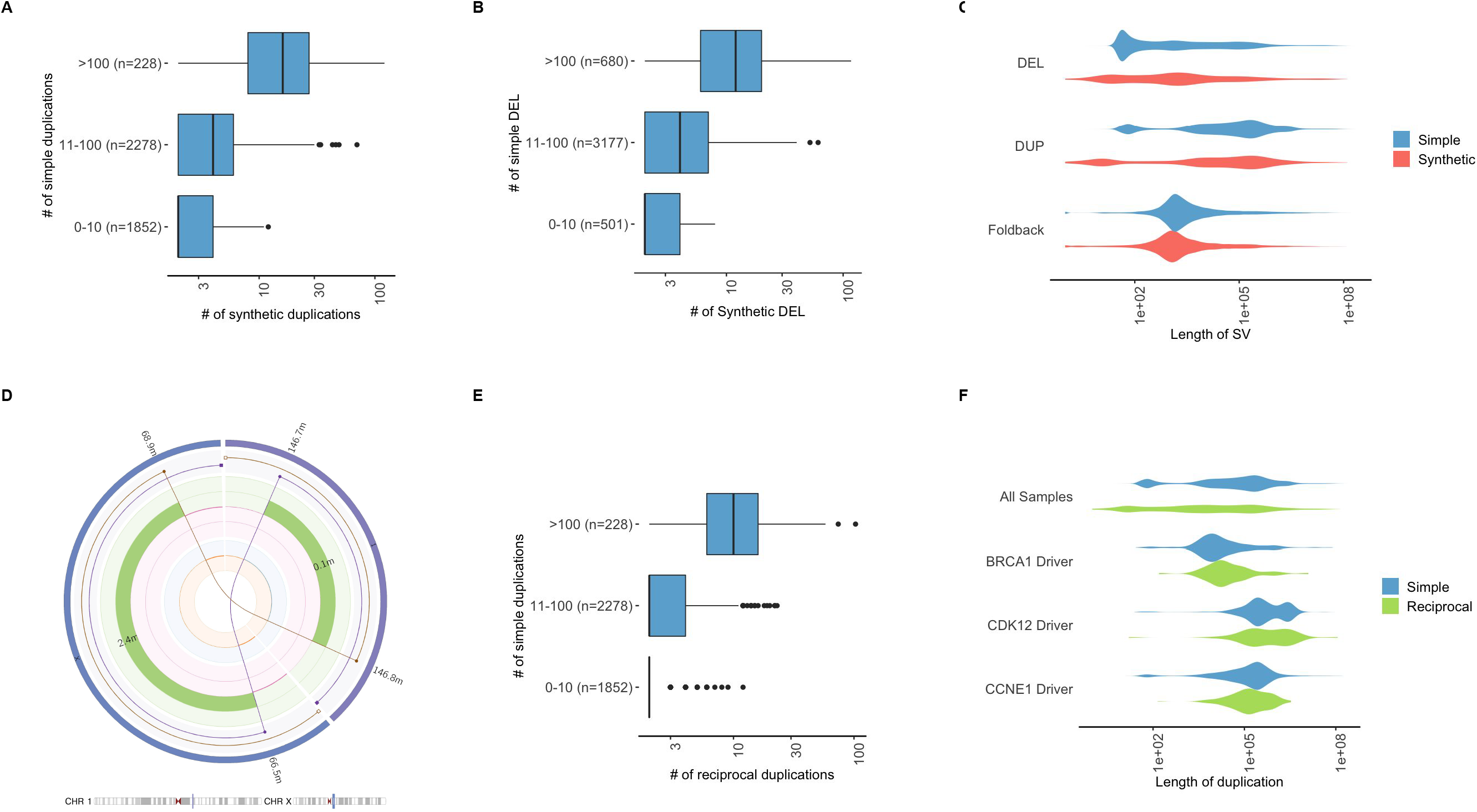
Synthetic events and reciprocal duplicaitons. a) & b) Samples with high number of simple duplications (deletions) also have high numbers of ‘synthetic’ duplications (deletions) c) Length distribution of simple and synthetic deletions, tandem duplications and foldback inversions in HMF cohort showing similarity in length distributions. d) Example of a predicted reciprocal duplication event involving a pair of translocations from chr1 to chrx in HMF003502A, a Breast cancer. The brown and purple lines show 2 derivative chromosomes formed by repairing a pair of overlapping breakends on each original chromosome. The rearrangement causes duplication of the regions between the facing breakends on both chromosomes (visible as increased copy number gain in green on the middle track). e) Samples with high number of simple duplications also have high numbers of reciprocal duplications f) Length distribution of simple and reciprocal duplications in HMF cohort overall and for samples with high confidence BRCA1, CDK12 and CCNE1 drivers showing similarity in length distributions.

**EXTENDED FIGURE 5:**
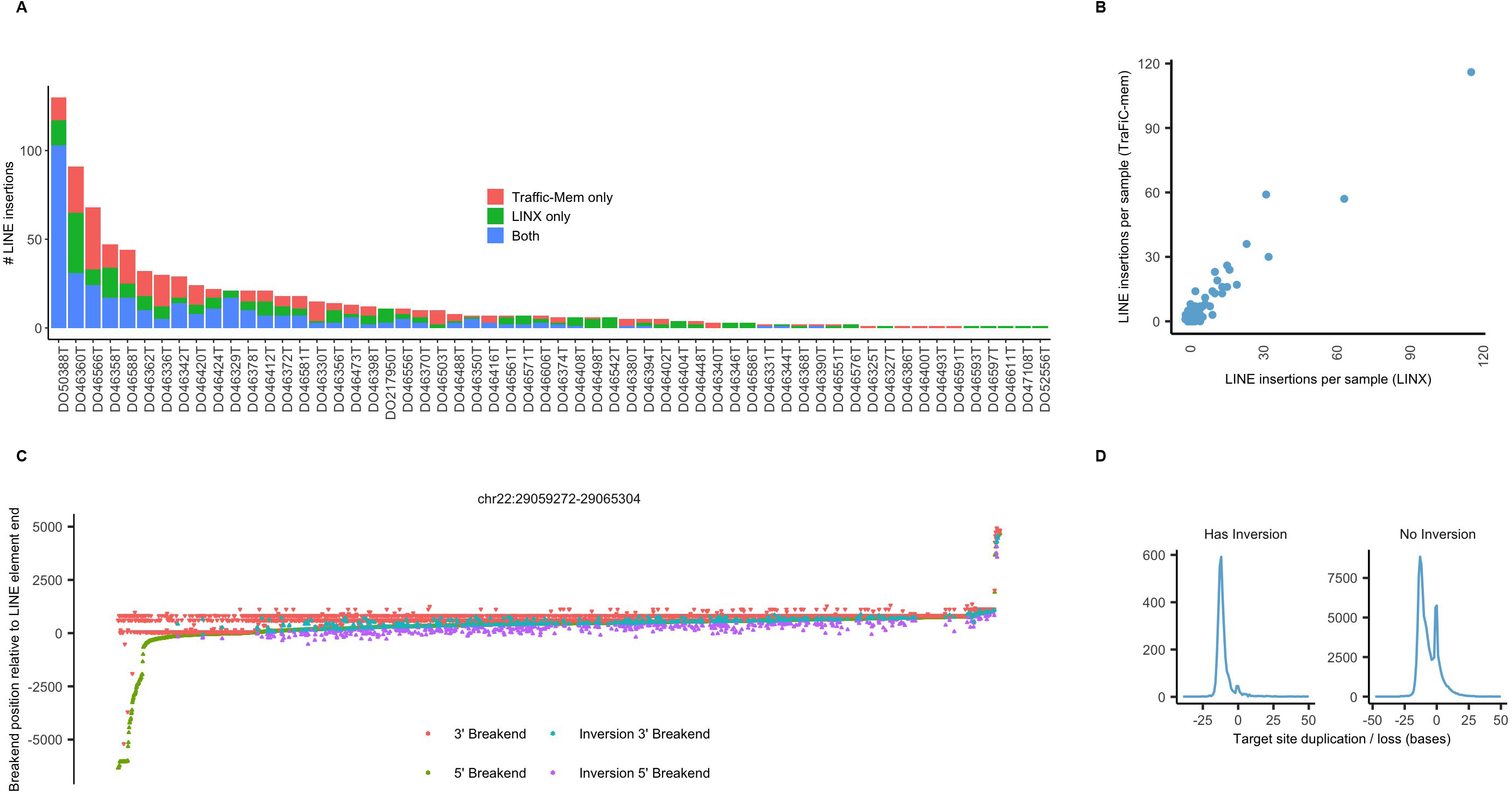
LINE insertions validation and analysis. a) Counts of LINE insertions predicted for 75 samples from the PCAWG cohort showing insertions called exclusively in LINX, exclusively in Traffic-Mem and shared in both tools b) Counts of LINE insertions per sample are highly correlated between LINX and Traffic-Mem c) Breakends positions for all LINE insertions in HMF cohort originating from the frequently somatically activated LINE source element at chr22:29,059,272-29,065,304 relative to the last base of the LINE element in the ref genome. Each column represents one insertion with start and end base of insertion indicated, as well as inversion breakpoints if the insertion contained an inversion. d) Distribution of target site duplication or loss length for insertions with and without an inversion. Negative values indicate duplication and positive values indicate a loss in bases.

**EXTENDED FIGURE 6:**
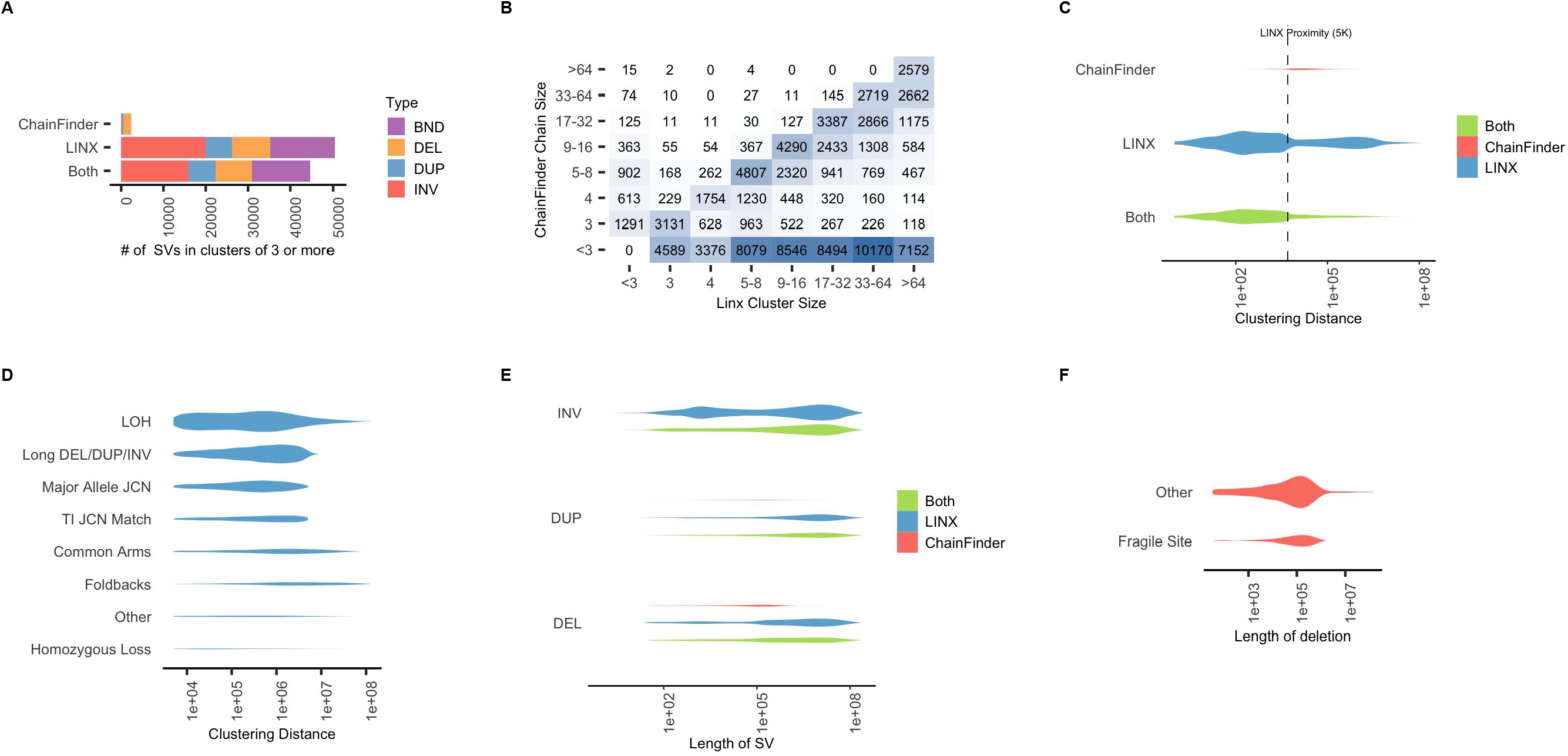
Complex clustering validation. a) Counts of junctions from 4,358 samples clustered into clusters of 3 or more variant by ChainFinder exclusively, LINX exclusively or by both LINX and ChainFinder. More than half of all variants are clustered exclusively by LINX, whereas very few clusters are private to ChainFinder b) Heatmap of counts of variants clustered into complex clusters by either LINX or ChainFinder showing ChainFinder versus LINX cluster size. Most clusters either have a similar size in both LINX and ChainFinder or are not found at all by ChainFinder c) Violin plot showing distribution of distance to nearest clustered breakend for all clustered variants by source. The majority of variants clustered in LINX but not chainfinder, have another breakend within 5kb (LINX proximity clustering distance). Nearly all variants with clustering distance >5kb were clustered exclusively by LINX. Area of violin proportional to count of variants. d) Distribution of distances to nearest clustered breakend for variants clustered exclusively by LINX and for reasons other than proximity. Area of violin proportional to count of variants. e) Length distribution of same chromosome structural variants clustered by LINX only, Chainfinder only and both. Area of violin proportional to count of variants. Note that ChainFinder private variants are predominantly deletion and duplications <1Mb, whilst LINX private variants follow a similar length distribution to shared variants. f) Distribution of distances to nearest clustered breakend for deletions clustered by ChainFinder. Area of violin proportional to count of variants. A significant minority of deletions occur in a handful of fragile sites regions which make up <1% of the reference genome.

**EXTENDED FIGURE 7:**
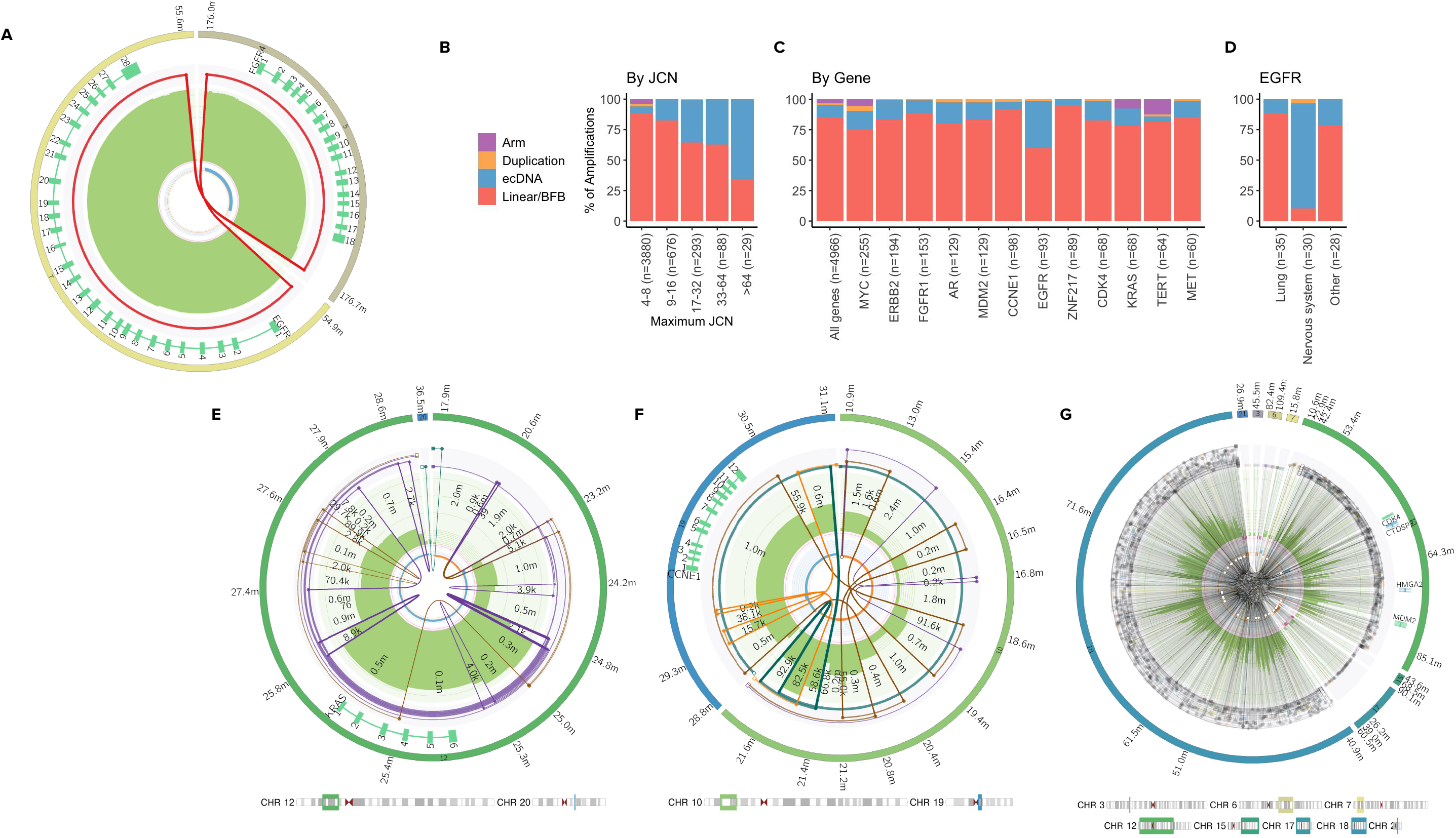
High amplification drivers. a) An ecDNA causing amplification of copy number ~60 in HMF002478A, consisting of 2 translocations which amplifies both *FGFR4* on chromosome 5 and *EGFR* on chromosome 7 b)-d) Proportion of amplification drivers which are predominantly caused by each of linear amplifications or breakage fusion bridge,ecDNA, simple duplication or arm level amplifications by maximum JCN (b), driver gene (c) and by tumor type for EGFR amplifications only (d). e) typical breakage fusion bridge event with 6 foldback inversions leading to high amplification of KRAS in HMF003560A, an ovary tumor. The brown and purple lines show 2 partial and incomplete chains of the foldback event. The green line shows a separate derivative chromosome formed from the same event, clustered due to the breakends bounding the same region of loss of heterozygosity f) Complex amplification event affecting CCNE1 in HMF001857A, a non-small cell lung carcinoma. The cluster is not fully resolved by LINX and is instead chained into 4 partial chains (shown in orange, brown, green and purple). The event does not have any foldback inversions, but also lacks regions and breakpoints of uniform high junction copy number expected in ecDNA. g) A highly complex rearrangement with over 800 junctions in HMF003994A, a malignant peripheral nerve sheath tumor, leading to coamplification of MDM2 and CDK4.

**EXTENDED FIGURE 8:**
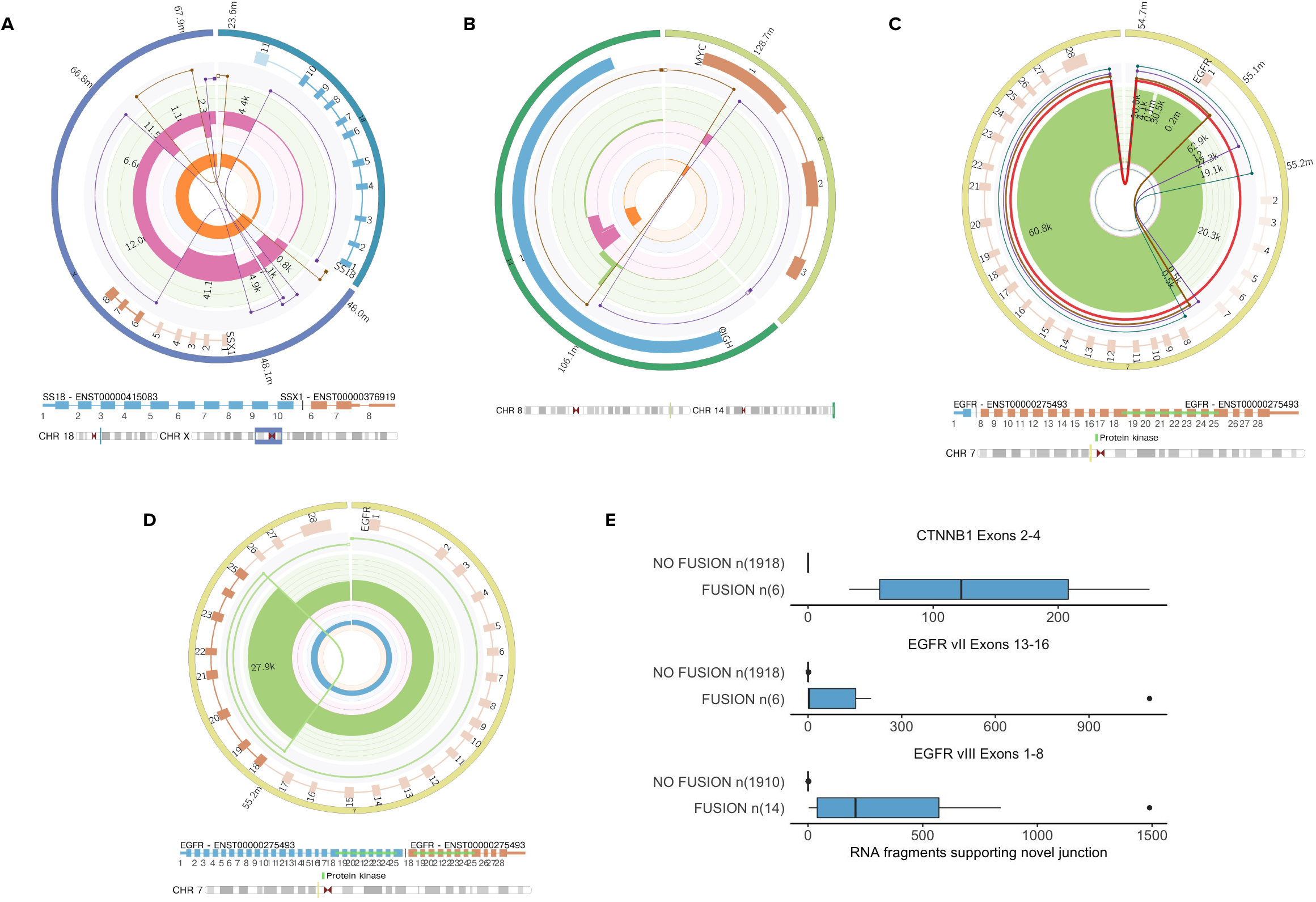
Diversity of fusion types. a) A *SS18-SSX1* chained fusion caused by a complex cluster of 6 variants in HMF003579A, a Sarcoma. The predicted derivative chromosome links exons 1-10 in *SS18* to exons 6-8 in *SSX1* via a chain of 2 structural variants. b) A reciprocal translocation in HMF003019A, a Lymphoid tumor, causing a pathogenic Immunoglobulin rearrangement event by moving the IGH Eμ enhancer adjacent to *MYC*. c) A double minute in *EGFR* in HMF000649A, a Glioblastoma, formed by a single duplication variant with 3 subsequent trans-phased (and hence independent) subclonal internal deletions of exon 2-7. d) An *EGFR* kinase domain duplication in HMF004524A, a Lung tumor. Exons 18-25 are duplicated. e) Counts of RNA fragments with splicing supporting exon deletions for RNA samples with and without the LINX fusion prediction for 3 highly recurrent exon deletions

**EXTENDED FIGURE 9:**
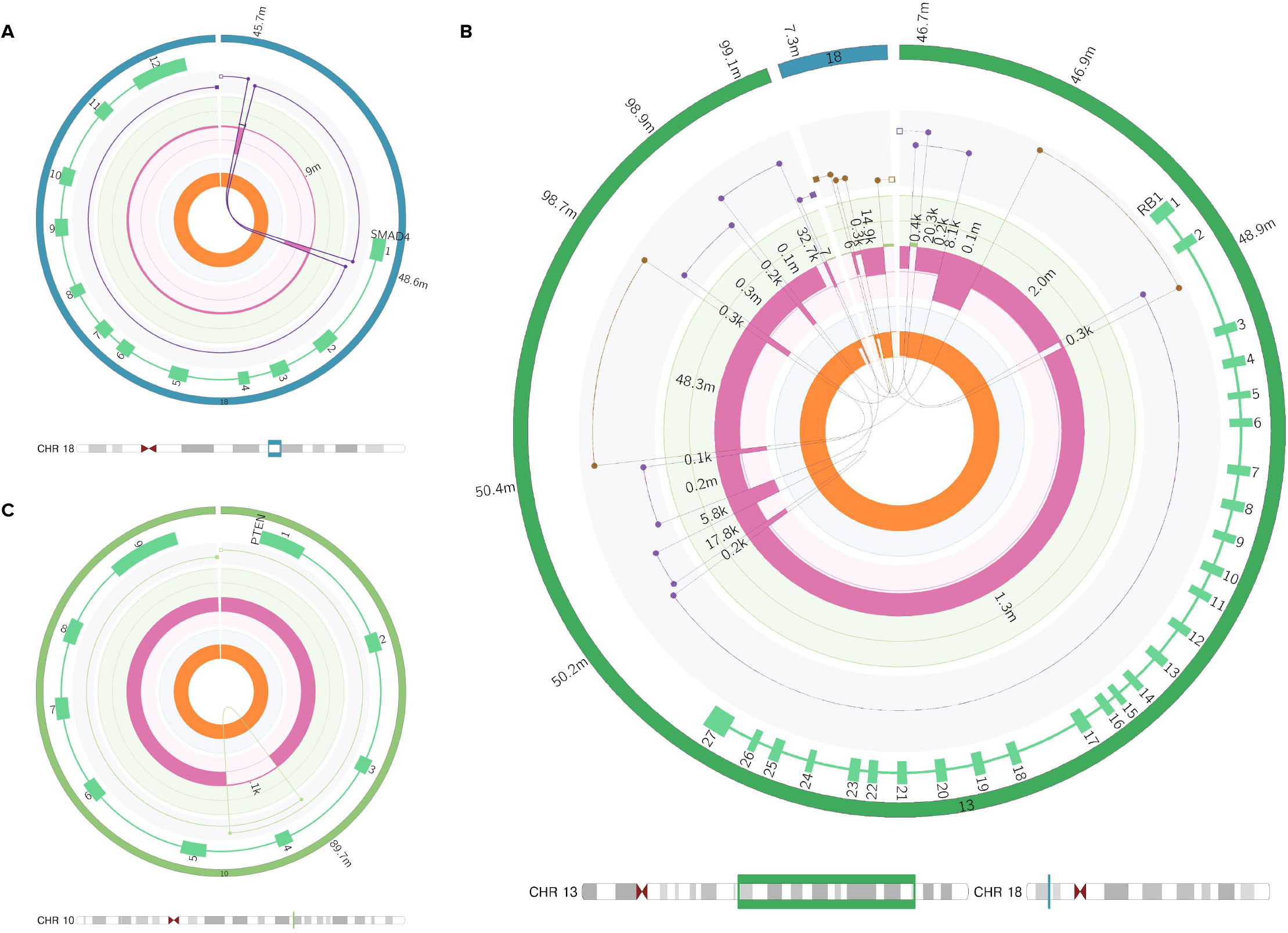
Diversity of homozygous disruptions. a) A reciprocal inversion in HMF002869A, a colorectal tumor, which homozygous disrupts *SMAD4* by reversing the direction of exon 1 on the derivative chromosome. b) A chromoplexy like event in HMF002065A, a prostate cancer which homozygously disrupts *RB1*. Exons 1 and 2 are predicted to be inserted into chromosome 18 on a separate derivative chromosome (shown in brown) from exons 3-27 which are part of a rearranged chromosome 13 (shown in purple) c) A homozygous disruption in *PTEN* in HMF001353A, a melanoma, caused by a simple tandem duplication which duplicates exon 4. Since the other parental chromosome is lost, the duplication is predicted to disrupt the last remaining copy of *PTEN*

## Supplementary Data

**Supplementary table 1**: Panel of driver genes used in study

**Supplementary table 2**: Hartwig DNA and RNA samples used in study

**Supplementary table 3**: Counts of structural variants by sample and classification

**Supplementary table 4**: LINX and Traffic-Mem LINE insertion predictions for 75 PCAWG samples

**Supplementary table 5**: ecDNA validation on 13 glioblastoma samples

**Supplementary table 6**: Curated fusion knowledge base used to search for pathogenic rearrangements

**Supplementary table 7:** Inframe pathogenic rearrangements detected in 1924 samples by DNA and / or RNA fusion calling

**Supplementary table 8:** Pathogenic homozygous disruptions detected in tumor suppressor genes in 4,358 samples

**Supplementary file 1:** LINX algorithm detailed description

**Supplementary file 2:** COLO829T input files required to run LINX

**Supplementary note 1:** COLO829T walk through of LINX results and visualisations

## Notes

### Competing Interest Statement

The authors have declared no competing interest.

