## Supplementary File 1 - Linx Methods for "Unscrambling cancer genomes via integrated analysis of structural variation and copy number"

### Key concepts in LINX

LINX is an annotation, interpretation and visualisation tool for somatic structural variants using genome-wide GRIDSS breakpoint and PURPLE copy number data as input. The primary function of LINX is to group together individual SV calls into rearrangement 'events' and properly classify and annotate each event to help understand both its mechanism and genomic impact.

#### LINX terminology and conventions for linking proximate breakends

##### Assembled, transitive and inferred links

In LINX, 'links' are chromosomal segments flanked by cis phased junctions which are predicted to form part of a single derivative chromosome. Assembled links are those that were called in a single assembly by GRIDSS and are very high confidence somatically phased. Transitive links are not fully assembled by GRIDSS but there is discordant read pair evidence supporting the link as a whole. All other links are inferred, based on proximity, topology and copy number characteristics using the chaining logic described below.

##### Templated insertions

We have adopted the term 'templated insertion' as has been used previously (Li et al. 2020) to describe any piece of DNA which is a templated sequence from a section of the reference genome flanked by breakends on either side inserted elsewhere (either locally or on a remote chromosome) into a chain to form part of a derivative chromosome. The inserted DNA may be either cut (causing disruption) or copied from the template.

##### 'Shards' and 'synthetic' events

A special and very common case of templated insertions we observe are very small templated genomic fragments of up to several hundred bases in length, which are frequently inserted into breakpoints without disruption at the source site for the inserted sequence. These have been observed previously (Bignell et al. 2007) and termed as genomic 'shards'. In LINX we model shards explicitly as short templated insertion lengths of less than 1k bases. These inserted sequences can make simple events such as deletions and tandem deletions appear to have complex topologies. For example, if we have a simple short deletion with a shard inserted, and the templated sequence of the shard is from another chromosome the deletion now presents notionally as a chained pair of translocations. Where more than 1 shard is inserted, the complexity can grow even further. LINX simplifies events that could be explained as a 1 or 2-break clusters with shards and marks the cluster as 'synthetic'.

Extended Figure 1 shows a number of examples of synthetic events with the shards marked.

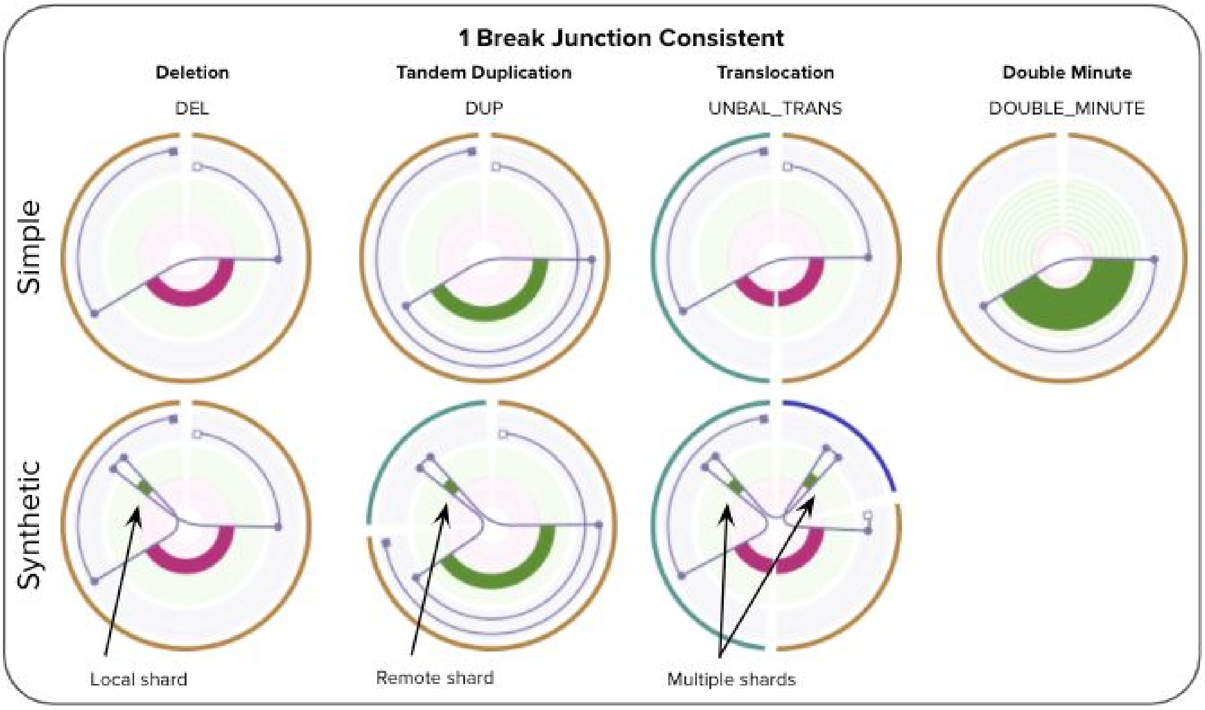


#### Deletion bridges, anchor distance & overlapping deletion bridges

We use the term ‘deletion bridge’ as defined previously (Baca et al. 2013) to refer to sections of DNA loss between 2 breakpoints on the same paternal chromosome that are fused to other segments of the genome.  GRIDSS provides an anchor support distance for each structural variant breakend which is the number of bases mapped to the reference genome at that breakend as part of the assembly contig, which is typically in a range from 29 bases (the minimum anchor distance for GRIDSS to be able to call) up to approximately 800 bases for short read sequencing. Any other breakend that falls within this anchor distance cannot be ‘cis’ phased with the variant as the contig was able to be mapped past the breakend and the 2 breakends are deemed to be ‘trans’ phased. Trans breakends within this distance range are common in cancer. One possibility is that the breakends could occur on the other paternal chromosome, but this highly unlikely as there is no reason to expect 2 different paternal chromosomes to both be damaged within a few hundred base region. Much more likely is that when the double stranded break occurred, that there was significant overlap between the break locations on the 2 strands and the shorter strand of each overlapping break end has been repaired prior to fusing with other regions of the genome . This is highly analogous to a deletion bridge except with small sections of replication of DNA instead of loss. LINX uses the term ‘overlapping deletion bridge’ to describe this breakend  topology.

#### Copy number conventions

LINX determines the number of absolute copies of each rearrangement junction in a sample, and terms this as the “junction copy number” (JCN). PURPLE SV output provides both a raw estimate of the JCN (estimated from the purity adjusted VAF of the junction) as well as the change in copy number observed at each breakend.  LINX uses both the raw estimate and the copy number change to predict both a JCN point estimate and uncertainty for each rearrangement junction.

#### Overview of event classification system in LINX

LINX attempts to classify all variants into a set of consistent events, i.e. events that transform the genome from one stable configuration into another.  LINX classifies all events with one or two junctions and groups events with 3 or more junctions as ‘COMPLEX’.

A key assumption in LINX is that each derivative chromosome arm in a stable configuration must connect a telomere to a centromere (since centromere to centromere joins will cause unstable breakage fusion bridge and telomere to telomere joins will have no centromere and will be ultimately lost during stochastic mitosis processes). A special case is allowed in highly restricted circumstances for double minute chromosomes which are circular and have no telomere or centromere but are highly positively selected for. This assumption means that variants such as a lone head to head or tail to tail inversion are considered incomplete, and in these cases we intensively search for other variants which may have occurred concurrently and could restore a stable configuration. Because of limitations of both input data accuracy and completeness and our clustering and chaining algorithm, many COMPLEX clusters will not be fully resolved to a stable configuration although it is assumed that such a resolution exists. Furthermore, we have a number of residual 1 and 2 clusters (eg. a lone inversion) which are inconsistent cannot be accurately clustered and hence we classify them as INCOMPLETE.

Ultimately we classify each cluster into 1 of 7 major event categories:

| Event Category | Description |
|---|---|
| SIMPLE | Single junction cluster which forms a local deletion, tandem duplication or unbalanced translocation |
| RECIPROCAL | Reciprocal inversion or translocation events forming from 2 concurrent breaks interacting with each other |
| TEMPLATED INSERTION | DEL or DUP or unbalanced translocation (‘chain’) with templated insertion |
| INSERTION | SV that are formed by the insertion of a templated piece of DNA normally via either a mobile element insertion or virus. |
| DOUBLE_MINUTE | Any 1 or 2 variant cluster where all variants form part of an ecDNA ring |
| COMPLEX | Clusters with 3 or more variants that cannot be resolved into one of the above categories |
| INCOMPLETE | 1 or 2 breakpoint clusters which are inconsistent, but cannot be clustered further OR clusters which are inferred from copy number changes only |

A brief overview of  each of the non SIMPLE categories is given below:

#### Reciprocal events

Linx models reciprocal 2-break junction clusters as events that could be caused by the interaction of 2 simple local concurrent breaks which would normally form deletes and tandem duplications. Depending on whether the breaks are on the same or different chromosomes this forms reciprocal inversions or reciprocal translocations respectively. Note that in the translocation case, if one side of the reciprocal event is subsequently lost either before or after repair, then we will instead observe an unbalanced translocation .

The possible geometries for reciprocal events supported by LINX are explained in the table below and drawn in extended figure 2:

| Interacting Break Types | Same chromosome (Inversion) | Translocation |
|---|---|---|
| Concurrent double stranded breaks | **RECIP_INV** - 2 facing inversions with outer breakends overlapping | **RECIP_TRANS** - 2 translocations forming deletion bridges on both arms. |
| Concurrent tandem duplications | **RECIP_INV_DUPS** - 2 facing inversion with inner breakends overlapping | **RECIP_TRANS_DUPS** - 2 translocations with facing breakends on both arms |
| Tandem Duplication + Double Stranded Break | **RECIP_INV_DEL_DUP -** inversion enclosing inversion with opposite orientation | **RECIP_TRANS_DEL_DUP -** 2 translocations forming a deletion bridge on one arm and facing breakends on other arm |

A facing pair of foldback inversions (FB_INV_PAIR) is also classified as a reciprocal, although the mechanism for forming this structure is unclear. It is possible that many of these events are formed from a breakage fusion bridge event but have not been properly clustered with a resolving break junction which may be distant in a breakage fusion bridge scenario.

#### Templated insertions

For the 4 reciprocal event cases above involving duplication (ie. RECIP_INV_DUPS, RECIP_INV_DEL_DUP, RECIP_TRANS_DUPS & RECIP_TRANS_DEL_DUP), the same junctions can be alternately chained to form a single derivative chromosome with a templated insertion (see extended figure 2). LINX gives precedence to the reciprocal interpretation, but if any of the duplicated segments bound a telomeric or centromeric loss of heterozygosity, the reciprocal interpretation is implausible

A deletion and duplication can together also form either a duplication or deletion with templated insertion structure (extended figure 2) identical to the 2 inversion case but with the inserted segment in the opposite orientation. Unlike inversions, simple deletions and tandem duplications are consistent standalone events and are common genomic events so some of these structures may be clustered incorrectly where separate DEL and DUP events are highly proximate or overlapping by chance.

#### Insertions

An insertion event is modelled by LINX as a pair of structural variants which inserts a section of templated sequence from either another part of the genome WITHOUT disruption to the

DNA at the source location OR from an external sequence such as an insertion from a viral genome.

The most common class of insertion in tumor genomes by far are mobile element insertions, which are not typically active in the germline, but can be highly deregulated in many different types of cancer. Mobile elements insertions frequently insert short sequences of their own DNA sequence and templated segments from adjacent to the source LINE element, with sometimes many segments from the same source location being inserted at multiple locations around the genome (Rodriguez-Martin et al. 2020). Mobile LINE elements can also cause SINE and pseudogene insertions. LINE insertion source breakends can be often difficult to map correctly on both ends, since they typically involve a repetitive LINE element at the start of the insertion element and a poly-A section at the end of the inserted section. LINX uses a combination of previously known active LINE element source information and identification of both the local breakpoint structure and POLY-A sequences to classify both fully and partially mapped breakpoints as LINE insertions.

##### Double minute

Any 1 or 2 variant cluster which is predicted to form a closed loop by LINX without a centromere is resolved as a ‘double minute’.   All variants must form part of the ecDNA to be classified as event type double minute, although ecDNA may also occur as part of a complex cluster.    An exception is made for a simple DUP double minute clustered with an enclosing DEL, which is classified as double minute despite the DEL not being a part of the ecDNA structure.   Complex clusters may also contain double minutes.

##### Complex events

COMPLEX events are defined in LINX as clusters with 3 or more variants that cannot be resolved into either a simple or synthetic type of insertion, DEL, DUP or 2-break event.

COMPLEX events may be formed by any combination of non-mutually exclusive processes including multiple concurrent breaks, replication prior to repair, breakage fusion bridge processes. Local topology annotations in LINX are intended to shed light on these complex processes.

##### Incomplete events

There are a number of possible configurations which are not ‘COMPLEX’ by the above definition since they are formed from 1 or 2 SVs, but lead to inconsistent genomes or involve single breakends. For these clusters there is assumed to be missing SVs, potential false positive artefacts or under clustering and they are marked as INCOMPLETE.

INCOMPLETE includes but is not limited to the following configurations:

- Lone inversion
- Lone single breakend
- Lone inferred breakend
- Any 2-break junction cluster with a single or inferred breakend that cannot be resolved as LINE or inferred as a synthetic.
- Any 2-break junction cluster which cannot be chained OR resolved as either a LINE, synthetic,templa or reciprocal event

Clusters of 2 inferred breakends are also classified in this category. Many of these are likely artefacts due to residual large scale GC biases affecting coverage unevenness in the sequencing data and false positive CNA calls.

### LINX Algorithm

There are 4 key steps in the LINX algorithm:

- Annotation of genomic properties and features
- Clustering of SVs into events
- Chaining of derivative chromosomes
- Gene impact and fusion prediction

The following schematic outlines the overall workflow in the LINX algorithm. Each step is described in detail below.

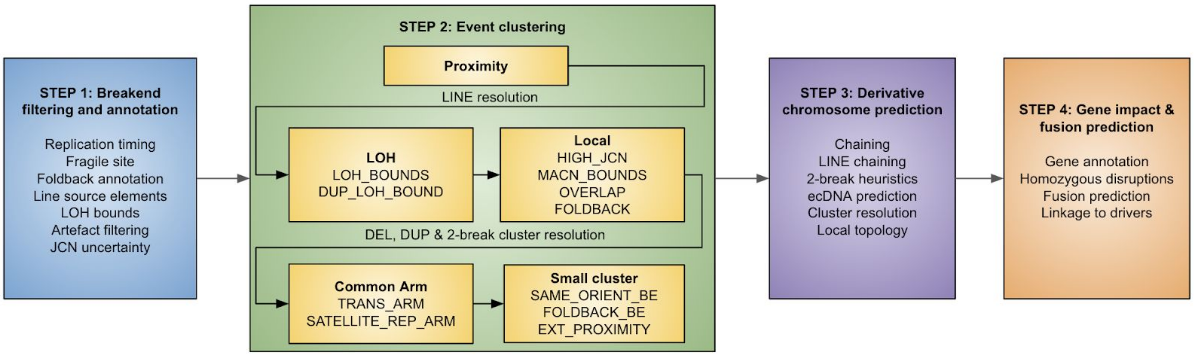


#### 1. Annotation of genomic properties and features

To help resolve and characterise events, LINX first annotates a number of genomic properties:

##### Externally sourced genomic annotations

Each breakend is first annotated with the following information from external sources
- Whether it is in a known fragile site (Priestley et al. 2019)
- Whether it is in a known LINE source element (Rodriguez-Martin et al. 2020)
- The replication timing of the breakend (ENCODE Project Consortium 2012)

##### Identification of foldback inversions

Foldback inversions are important structural features in the genome since they are a hallmark of the breakage fusion bridge process. LINX use foldbacks in a number of ways in both the clustering and chaining algorithms, and they can be objectively identified independently of the clustering and chaining so it is useful to identify them upfront. We perform a genome wide search for both simple foldback inversions and chained foldback inversions which are disrupted by the insertion of a shard. A pair of breakends is marked as forming a foldback if they meet the following criteria:

- the breakend orientations are the same and are consecutive (ignoring any fully assembled interim breakends) and after allowing for overlapping deletion bridges on both ends, specifically both:
    - The outer breakend may be overlapped by a variant within it's anchor distance
    - The inner breakend may not have a facing breakend within it's anchor distance
- the breakends belong to a single inversion or are linked by an assembled or short chain (<= 5K bases)
- A single breakend where the other end of the structural variant is assembled to itself via a chain
- Neither breakend forming the foldback is linked via assembly to another breakend.

##### Identification of suspected LINE source elements

LINE source elements are also important genomic features and are modelled in LINX as regions of ~5000 bases which LINX suspects are the source for templated LINE insertions. LINE driven mobile element insertions are common in cancer genomes and typically present as a pair of balanced SVs representing a templated sequence from around the source element with a poly-A tail inserted into random locations in the genome although favoring a A|TTTTT motif (Rodriguez-Martin et al. 2020)for the insertion site with no net copy number change at either source or insertion site. However, due to both the repetitive nature of the LINE source elements and the difficulty of accurately sequencing across the poly-A tail, one or both of the SVs that make up the insertion may be mapped as a single breakend (failing to uniquely map on the other side) OR be missed altogether. Since the lone remaining breakend can be mistaken as an unbalanced translocation, it is important to correctly identify it as a LINE element. This picture can be complicated even further by the fact that many overlapping fragments from a single source location may be copied to many different locations in the same genome, each potentially with one or both sides incompletely mapped.

Although we already annotate 124 well 'known' mobile LINE source elements which have been previously discovered(Rodriguez-Martin et al. 2020), there are many more potential mobile LINE source elements which may be less common in the population or more rarely activated. We look exhaustively for likely LINE source elements in each individual tumor genome, by looking for both the characteristic poly-A tail of mobile element insertions and the local break topology structure at the insertion site. We define a poly-A tail as either at least 16 of the last 20 bases of the sequence are A or there is a repeat of 10 or more consecutive A or within the last 20 bases of the insert sequence. The orientation of the breakend relative to the insertion can help distinguish between the source and insertion site for a mobile element. At the mobile element source site, the poly-A tail positive oriented breakends will have the poly-A at the start of the insert sequence, or poly-T at the end of the insert sequence for negative oriented breakends (if sourced from the reverse strand). Conversely at the insertion site, negative oriented breakends will have poly-A tails at the end of the insert sequence and positive oriented breakends have poly-T at the start of the insert sequence (if inserted on the reverse strand)

Each breakend will be classified as being a 'Suspected' LINE source element if any of the following conditions are met:

- there are 2+ breakends within 5kb with poly-A/poly-T tails with expected orientations for a source site.
- there are 2+ translocations or local junctions > 1M bases which are not connected at their remote end to a known LINE site within 5kb with at least one not forming a

deletion bridge of < 30 bases AND at least one breakend within 5kb having a poly-A tail with expected orientation for a source site
- we find at least 1 translocation or local junction >1M bases with it's remote breakend proximity clustered with ONLY 1 single breakend and forming a deletion bridge of < 30 bases AND EITHER the junction has a poly-A / poly-T tail with the expected orientation of the source site OR the remote single breakend has a poly-A/poly-T tail with expected orientation for an insertion site.

The suspected LINE source element is also checked that it is not a potential pseudogene insertion by checking that there is no deletion within 5kb of the suspected source element that matches an exon boundary at both ends. Both known and suspected source elements have special logic applied in the clustering phase.

##### Identification of LOH boundaries

LINX also identify each pair of breakends flanking regions of Loss of Heterozygosity (LOH), restricted to cases where there is no subset of the region with homozygous loss that is not caused by anything other than a simple deletion. This is a useful annotation as since an entire paternal allele is lost for the whole distance between these 2 breakends (and since there is no homozygous loss we know it is the same allele lost at both ends, not two overlapping losses) then the structural variants are very likely to have occurred at the same time.

Note that an uninterrupted deletion or tandem duplication cannot theoretically form an LOH boundary with another variant and hence these are excluded from LOH boundaries.

##### Long DEL and DUP length calculation

Shorter deletions and tandem duplications are found frequently as standalone events in the tumor genome, but longer standalone deletions and duplications are relatively rare and when they do occur are often associated with more complex events. The following method is used to determine a characteristic length threshold for each sample for what is considered a 'long' DEL or DUP.

- find all DUPs and DELs on arms with no inversions (inversions are used as a proxy for the presence of complex events)
- Set LONG_DUP_LENGTH to the length of the longest DUP excluding the 5 longest DUP lengths (normalised for the proportion of arms without inversions). Min =100k, Max = 5M
- Set LONG_DEL_LENGTH to the length of the longest DEL excluding the 5 longest DEL lengths (normalised for the proportion of arms without inversions). Min =100k, Max = 5M

The threshold is subsequently used in clustering rules by LINX.

##### Estimation of JCN and JCN uncertainty per SV

We know that by definition JCN must equal the copy number change at start breakend and the copy number change at end breakend for each and every structural variant. However, frequently the JCN and copy number change at start and end may not match up for one of several reasons including measurement error, false positive SV artefacts and false negative SV calls adjacent to one or both of the breakends for the structural variant.

To allow for accurate chaining we would like to have a single consolidated JCN estimation and an idea about the uncertainty in the JCN for each SV. Since the distribution of the errors in our measurements are unknown and fat tailed due to potentially missing and false positive data, we create a simple model for a reasonable estimation of the likely JCN range.

We use 3 steps in the process:

##### 1. Estimate an uncertainty for copy number change at each breakend

For this use the principle that the uncertainty in copy number change is driver primarily by the uncertainty in the copy number of the least confident adjacent copy number region which in turn is driven primarily by the number or read depth windows used to estimate the length of the adjacent regions.

Hence we use the following formula to calculate a copy number uncertainty

> CNChangeUncertainty = MAX(maxAdjacentCopyNumber * BaseRelativeUncertainy [0.1],BaseAbsoluteUncertainty [0.15])+ MAX(AdditionalAbsoluteUncertainty [0.4],AdditionalRelativeUncertainty [0.15]*maxAdjacentCopyNumber)/SQRT(max(minAdjacentDepthWindowCount,0.1))

If the minAdjacentDepthWindowCount = 0, then this means the segment is inferred by the raw JCN in PURPLE already and no copy number estimate is calculated.

For the special case of foldback inversions, if the flanking depth window counts are both higher than the internal depth window count, a single copy number change observation of half the combined copy number change is made with the confidences determined from the flanking windows

##### 2. Estimate an uncertainty for raw JCN

The raw JCN of the SV is already estimated in PURPLE by multiplying the purity adjusted VAF of the SV by the copyNumber at each breakend. The VAF estimate depends ultimately on the measured readcount of supporting tumor fragments which is a binomial distribution.

To estimate the uncertainty in the VAF, we estimate the 0.5% and 99.5% confidence intervals of the true read count from the observed read count and then calculate the JCN uncertainty as half the relative range of the confidence interval. We also add half the minimum of the 2 breakend copy number uncertainties to reflect the copyNumber impact on uncertainty. This gives the formula:

> JCN Uncertainty = JCN * (ReadCountUpperCI-ReadCountLowerCI) / 2 / ObseverdReadCount + 0.5 * min(CNChangeUncertaintyStart,CNChangeUncertaintyEnd)

##### *3. Average the 3 JCN predictions and estimate a consolidated uncertainty*

Weight the observations by the inverse square of their estimated uncertainties:

> consolidatedJCN = SUM[Observation(i)*(1/Uncertainty(i)^2)] / Sum[1/Uncertainty(i)^2]

The combined uncertainty is estimated as the square root of the weighted sum of squares of the difference between the final JCN estimate and each individual estimate, but capped at a minimum of half the input uncertainty. Ie.

> consolidatedUncertainty = SQRT( countObservations / (countObervations-1) *
> SUM[1/Uncertainty(i)^2*(MAX(Observation(i)-consolidatedJCN,Uncertainty(i)/2))^2] / Sum[1/Uncertainty(i)^2] )

##### Artefact filtering

Prior to clustering and chaining, LINX applies additional artefact filtering, since despite applying stringent filters in both PURPLE and GRIDSS upstream, we may still find a number of false positive artefacts in our data. False positive artefacts are typically either SVs with little or no copy number support (ie copy number change at both breakends < 0.5) or inferred SV breakends from the copy number analysis. Unfortunately these can be difficult to distinguish from bonafide subclonal variants with ploidies of 0.5, and from genuine clonal variants where we have missed an offsetting SV call which netted out the copy number change.

To remove residual artefacts, but preserve genuine subclonal variants, we limit filtering to 3 very specific situations which strongly appear to be artifactual in our data:

- **Short foldbacks Inversions (<100 bases) unsupported by copy number change** - Typically foldback inversions range from several hundred to several thousand bases. However, we also frequently find many very short foldbacks inversions in highly damaged samples. Across the cohort as a whole we find these to overwhelmingly have low junction copy number and with little copy number support. Hence we mark foldback inversions <100 bases in length as artefacts if both start and end copy number change < 0.5 or VAF < 0.05 at both ends.
- **Isolated translocations and single breakends unsupported by copy number change at both breakends -** These also common artefacts and similar to foldback inversions, we find an elevated rate of low junction copy number variant calls unsupported by copy number change indicating likely artefacts. We filter if both breakends of a translocation are > 5000 bases from another variant AND both breakends have copy number change <0.5 AND the insert sequence neither has a poly-A tail nor insertSequenceRepeatClass = 'LINE/L1' (which may indicate a Line insertion).
- **Neighbouring inferred breakends with opposite orientation and matching or overlapping JCN uncertainty** - Residual GC bias and other forms of read depth noise can cause many inferred segments to be called. These are unhelpful in chaining and are resolved in pairs of offsetting variants as artefacts

All of the filtered variants will be marked as resolved type = ARTIFACT and will be restricted from any subsequent clustering.

#### 2. Clustering of structural variants into events

LINX uses a clustering routine to classify events. All SVs within a sample are grouped into clusters in 7 steps:

- Proximity clustering
- Resolution of LINE clusters
- LOH clustering
- Local clustering
- Resolution of simple events
- Common arm clustering
- Incomplete and small complex cluster merging

#### Proximity clustering (PROXIMITY)

Any SV with a breakend within the specified proximity_distance (default = 5Kb) of another SV's breakend causes the SVs to be clustered. An exception is made for overlapping DELs, which are split out into separate clusters, on the assumption that overlapping DELS must by definition be trans-phased.

#### Resolution of LINE clusters

LINX performs early resolution of mobile element insertions as insertions typically cause translocations that may inadvertently be clustered with other variants, particularly in tumors with highly deregulated LINE activation.

LINX resolves a cluster as type LINE if any of the following conditions are met:

- It contains a suspected LINE element AND (the cluster has <=10 variants OR at least 50% of the non single or inferred junctions in the cluster have a known or suspected breakend)
- Every non single or inferred breakend variant in the cluster touches a KNOWN LINE element AND at LEAST one of the variants is a translocation

Due to the low mappability of the LINE elements, frequently we will also identify LINE insertions only as single breakends as the insertion site. We therefore also resolve a cluster as LINE insertion if it contains either :

- 2 single breakends only, forming a deletion bridge (less than +/-30 bases) with [one side having a poly-A/poly-T tail with the expected orientation of an insertion site OR one of them has insertSequenceRepeatClass = 'LINE/L1']
- 1 single breakend and 1 inferred breakend only and the single breakend has insertSequenceRepeatClass = 'LINE/L1' or a poly-A/poly-T tail with the expected orientation of an insertion site
- 1 single breakend with insertSequenceRepeatClass = 'LINE/L1' or a poly-A/poly-T tail with the expected orientation of an insertion site and copy number change < 0.5 / 15%

LINE clusters are excluded from all subsequent clustering rules.

#### LOH Clustering

##### Loss of heterozygosity bounds (LOH_BOUNDS)

The 2 breakends forming the bounds of any LOH region not disrupted by a homozygous deletion are clustered together, reflecting the fact that both ends of the lost paternal chromosome must have been lost at the same time.

For LOH regions which are disrupted by 1 or more homozygous deletions, both paternal chromosomes are presumed to have been deleted in separate events and one deletion may enclose the other deletion OR they may overlap each other. In the enclosing case, for homozygous deletions which are in a region where the surrounding LOH bounds which form a simple DEL or are already linked or extend to at least the full arm, then we cluster the 2 homozygous deletion bounds. Conversely if all the homozygous deletions inside a LOH region are simple DEL or are already linked then we can cluster the 2 LOH region bounds. Finally, in the overlapping case, if all the overlapping deletion bounds are from simple DELs or are already linked except for 2 breakends, then link the remaining 2 breakends.

Note that variants with breakends that bound an LOH are not permitted to link with breakends in the bounded LOH region via any subsequent rule, since by definition they are expected to be on the other paternal chromosome.

###### Chaining bounds for DUP variants causing LOH (DUP_LOH_BOUNDS)

No breakend in a cluster can chain across an LOH which has been caused by a breakend in the same cluster. Hence if the other breakend of a DUP type variant bounding an LOH can only chain to only one available (not assembled, not LINE) breakend prior to the LOH, then we cluster the DUP and the other breakend.

##### Local Clustering

###### High JCN (HIGH_JCN)

Merge any pair of breakends which both have JCN> max(5, 2.3x the adjacent major allele copy number) that face each other that are also joined by a continuous region of majorAlleleCN > 5

###### Major allele copy number bounds (MACN_BOUNDS)

The major allele copy number of a segment is the maximum copy number any derivative chromosome which includes that segment can have. Hence a breakend cannot chain completely across a region with major allele copy number < JCN of the breakend, or partially across the region with a chain of JCN more than the major allele.

Therefore any breakend is clustered with the next 1 or more facing breakends (excluding LINE & assembled & simple non overlapping DEL/DUP) IF the major allele copy number in the segment immediately after the facing breakend is lower than the breakend JCN, after discounting facing breakends in the original cluster. In the case of there being more than 1 facing breakend, the highest JCN breakend is clustered and the process repeated. This clustering is limited to a proximity of 5 million bases and bounded by the centromere, since although more distal events on the same chromosome may be definitely on the same derivative chromosome, this does necessarily imply they occurred concurrently.

###### Local Overlap (OVERLAP)

Merge any clusters or SVs where each cluster has either an inversion or a DEL exceeding the LONG_DEL_LENGTH, or a DUP exceeding the LONG_DUP_LENGTH and they overlap or enclose each other AND the 2 variants have at least one pair of breakends not more than 5M bases apart that either face each other or form a deletion bridge.

##### Foldbacks on same arm (FOLDBACK)

Merge any 2 clusters with a foldback on the same chromosomal arm.

#### Resolve DEL, DUP and consistent 2-break junction events

At this step, DEL, DUP and reciprocal clusters are resolved to prevent them from over clustering with later clustering rules. Specifically, the following types of events are resolved:

- **Deletions** - Simple deletions less than the LONG_DEL_LENGTH
- **Tandem duplications** - Simple duplications less than the LONG DUP_LENGTH
- **Reciprocal events and 2-break junction templated insertions -** Pairs of overlapping inversions or translocations where the deletion bridge and/or overlap at both breakends are less than the LONG_DEL_LENGTH / LONG_DUP_LENGTH and neither deletion bridge or overlapping region contains a non simple DEL/DUP breakend.

We also allow for synthetic variants of these events to be resolved, where the variants create a DEL, DUP or reciprocal event with the same geometry but with one or more shards inserted. All of these simple and synthetic clusters types are excluded from all subsequent clustering rules.

#### Common arm clustering rules

##### Translocations with common arms (TRANS_ARM)

Merge any 2 unresolved clusters if they touch the same 2 chromosomal arms. SVs which link the 2 arms but are in a short templated insertion (< 1kb) are ignored.

##### Single breakends on same arm with matching satellite repeat type clustering (SATELLITE_SGL_ARM)

Where complex events touch satellite repeats we frequently find many single breakends on the same chromosome with links to the same type of repeat. In particular this can occur when shattering events include complex focal scarring in centromeric regions leading to many unresolved single breakends

We therefore merge any cluster with less than or equal to 1 non single breakend and non inferredbreakend with any other cluster which contains a single breakend on the same chromosome arm with matching repeat class or type for the following cases:

- RepeatClass = 'Satellite/centr’ (centromeric)
- RepeatType = '(CATTC)n’ (satellite repeat type)
- RepeatType = ‘(GAATG)n’ (satellite repeat type)
- RepatType = ‘HSATII’ (pericentromeric)

To protect against false positives and joining complex clusters that both touch repeats, but otherwise don’t appear to overlap, we avoid clustering 2 clusters which already have multiple non single breakends.

We don’t cluster other common sequences such as telomeric sequences, Sine/Alu or LINE/L1 as these tend to be associated with genome wide insertion patterns rather than specific clusters which touch a repetitive region.

#### Incomplete and small complex cluster merging

These rules are implemented to merge small unresolve with 3 or less variants to other unresolved clusters with an arbitrary cluster size where the location and orientation of proximate or overlapping breakends between the 2 clusters indicates that they may be linked.

##### Breakends straddled by consecutive same orientation breakends (SAME_ORIENT_BE)

Merge any non-resolved breakend to a cluster which straddles it immediately on both sides with 2 breakends facing the same direction, and where the facing breakends have matching JCN.

##### Breakends straddled by foldbacks (FOLDBACK_BE)

Merge any non resolved breakend into a cluster which has 2 different foldbacks straddling it immediately on both sides and at least one of the foldbacks faces the breakend.

##### Extended chainable proximity for complex and incomplete events (EXT_PROXIMITY)

Merge any neighbouring non resolved clusters that are within 5M bases and which have facing flanking breakends on each cluster which could form a templated insertion with matching JCN. In the case of a foldback the JCN of the facing breakend is also permitted to match 2x the JCN.

#### 3. Chaining of derivative chromosomes

A chaining algorithm is used to predict the local structure of the derivative chromosome within each cluster. The chaining algorithm examines each cluster independently and considers all possible paths that could be made to connect facing breakends into a set of continuous derivative chromosomes.

##### Overview of chaining model

Chains in LINX are a walkable set of linked breakends with a common JCN. Initially each cluster begins with 1 chain for every SV in the cluster. LINX iteratively makes 'links' between chain ends, resolving 2 of the chains into 1 combined chain and determining a combined JCN and JCN uncertainty for the new combined chain from the 2 constituent chains. The order of linking of chains is prioritised such that the most likely linkages are made first. Structural variant calls from GRIDSS that are shown to be linked empirically by GRIDSS are linked first followed by a set of heuristics to prioritise the remaining uncertain links. This process continues to extend the length of and reduce the number of chains in each cluster until no further links can be made.

###### Constraints on linking breakends

In general each pair of facing breakends is considered by LINX as a candidate chain. However, 3 key constraints are applied in limiting candidate pairs:

- **'Trans' phased breakends are not permitted to chain -** A breakend may not be chained to another breakend within it's anchoring supporting distance unless it is linked by assembly.

- **Closed loops without centromeres are only allowed for ecDNA** - With the exception of ecDNA (see below), derivative chromosomes are assumed to connect to either a centromere or telomere at each end. Hence, chains which do not cross a centromere are not allowed to form closed loops (ie. the 2 breakends flanking a chain are not allowed to join to each other) except when specifically identified as ecDNA.

- **A chain should not pass through a region with lower available allele copy number than the JCN of the chain** - Chains are generally assumed to take place on a single paternal chromosome except in the rare case that 2 distinct overlapping paternal alleles of the same chromosome are involved . The copy number for the affected allele is calculated for each segment adjacent to each cluster by first determining the copy number of the unaffected allele across each contiguous set of breakends in the cluster and subtracting that from the total copy number. The total JCN of linked chains crossing a segment should not exceed the calculated allele copy number for that segment. In some noisy copy number regions or where both paternal alleles are involved we may not be able to determine an undisrupted allele and the allele copy number is not calculated. Additionally, if all possible links have been exhausted using this rule, the rule is relaxed such that chaining can continue under the assumption that allele specific copy number may not have been estimated accurately.

##### Uniform JCN clusters

Uncertainty in JCN measurement is one of the key confounding factors in predicting the derivative structure. Many clusters however do have the same JCN for all variants and these uniform JCN clusters are significantly simpler to chain into a derivative chromosome as each pair of breakends can only be linked with a single JCN.

Hence, each cluster is tested to see whether all its SVs could be explained by a single JCN value, using each SV's JCN estimate and range. If all SVs have a JCN range which covers the same value, even if not an integer, then the cluster is considered to be of uniform JCN and no replication will occur in the chaining routine. Furthermore, if all SVs have a JCN min less than 1 and a max > 0.5, then the cluster is also considered uniform.

##### Variable JCN clusters

For all clusters that cannot be resolved by a single uniform JCN, further considerations apply to explain the amplification of parts of the cluster. Biologically, each SV initially occurs by joining 2 breakends with a JCN of 1. However, a single chain in a cluster may contain SVs with different ploidies as a result of a replication from either a foldback inversion in a breakage fusion bridge event, or a later tandem duplication of part of the derivative chromosome. In these scenarios, the duplicated variants will appear multiple times in a single chain either repeated in the same direction in the case of a duplication or inverted in the case of a foldback.

Duplication events are permitted to 'replicate' a chain in 2 different ways:

1. **Foldbacks:** Foldbacks with half the JCN of another chain are permitted to link both their breakends to the same breakend of that chain, making a new chain of half the JCN with the other unconnected breakend of the non foldback chain at both ends of the new chain. In this foldback replication case the new chain may also be treated as a 'chained foldback' and extended further in the same manner if possible.

2. **Complex Duplications:** Conversely, complex duplications with half the JCN of another chain are permitted to join both of their breakends simultaneously to either ends of the other chain, effectively duplicating the entire chain, but keeping the same start and end breakends with half the JCN.

Partial replication of a chain is also possible by later variants which affect chains that have also been duplicated. In this case the higher JCN chain is split into 2 separate chains, one which is linked to and given the JCN of the lower JCN chain, and the other which is given the residual JCN.

#### Implementation of chaining algorithm

Linx first resolves all assembled and transitive SVs in the cluster into chains. LINX then keeps a cache of a set of chains consisting initially of all 'single variant' chains (ie. lone SVs) and assembled chains. Each chain has 2 breakends, a JCN and a JCN uncertainty. Each potentially linkable pairs of facing breakends (subject to the available allele copy number and no closed loops rules described above) is also cached as potentially linked chains.

The following steps are then applied iteratively to join chains together until no more chain links can be made:

1. Apply priority rules below to choose the most likely linked pair of chains from the cache
2. Merge chains:
    a. Create new combined chain and calculate the JCN & JCN uncertainty
    b. For replication events, replicate and halve the JCN of the chain, inverting if it is a foldback type event.
    c. In the case of partially split chains, the higher JCN original chain is kept with its residual JCN
    d. Remove merged chains
3. Update cache of linked breakend pairs that include a breakend on the merged chains

##### Prioritisation of chain links

Where more than 1 possible pair of linkable chain exists in the cache, links are ranked and chosen by the following criteria in descending order of importance:

1. Links with available allele copy number*
2. Links containing highest JCN foldback or complex duplication chain
    a. Links where it splits another chain with 2x its JCN
    b. Links where it matches the JCN of another chain
    c. Links where it splits another foldback with greater than 2x JCN
    d. Links where it is itself split by another foldback or complex duplication with half the JCN
3. Breakends with a single link possibility
4. Links with highest matching JCN status (MATCHED > OVERLAPPING_JCN_RANGE > NO_OVERLAP)
5. Adjacent links
6. Higher JCN links (allowing for 0.5 abs and 15% threshold)
7. Shortest link

* For the purpose of the available allele copy number rule, a junction with an offsetting inferred breakend at the exact base is assumed to create a link if and only if the JCN of both the junction and the offsetting inferred breakend is greater than adjacent major allele copy number (allowing for 0.5 abs and 15% threshold)

For uniform JCN clusters, only rules 1, 3,5 & 7 are considered in the prioritisation of links.

A cluster can be chained into more than 1 chain, each one representing the neo-chromosomes resulting from the rearrangement event. Chaining is often imperfect and incomplete due to the inclusion of single breakends and uncertainty about JCN and subsequent breakend replication.

##### Special considerations for LINE clusters

LINE clusters typically involve one or more insertions from a single source location to multiple target sites in the genome, with occasional inversion or rearrangement of the inserted sequence. Due to the highly localised origin and frequent overlap of these inserted elements, the above chaining rules are not appropriate for LINE clusters. Instead, assembled and transitive links are chained first and then pairs of SVs or assembled/transitive chains that make consistent insertions to a single remote site are chained together. Single breakends at an insertion site that are paired with a breakpoint to a known or suspected LINE source element and have insert sequence alignment matching the same source LINE element with the correct orientation are also chained at the source site. SVs that cannot be paired off to form consistent insertions are not chained.

##### Special considerations for extrachromosomal DNA (ecDNA)

ecDNA ( or double minutes) and the SVs contained within them are subject to special chaining rules in LINX. They are therefore identified prior to chaining. The key principle used to identify ecDNA is to look for high JCN junctions adjacent to low copy number regions which can be chained into a closed or predominantly closed loop.

LINX use the following algorithm to identify ecDNA

- Identify candidate ecDNA clusters: the cluster contains at least 1 non deletion junction with one breakend with a JCN > 5 AND at least 2.3x the adjacent major allele copy number. For clusters with maxJCN < 8, or if the only candidate DM variant is a single duplication, then the JCN of all candidate breakends must be at least 2.3x the adjacent major allele copy number
- Find all other potential ecDNA junctions in candidate clusters: any variant with JCN > max(5,25% of max ecDNA candidate JCN) and >2x the adjacent major allele copy number is classed as a candidate ecDNA junction.
- Attempt to chain candidate ecDNA junctions into closed segments: LINX tries to chain the identified candidate ecDNA junctions both with uniform and variable junction copy number and chooses the chain with the most closed segments
- Determine whether chained ecDNA junction meets ecDNA criteria: All of the following criteria must be met either for a closed loop or for the all the DM candidate variants as a whole if a closed loop cannot be formed:
    - At least one pair of breakends must be chained to form a closed segment (ignoring non overlapping deletion junctions)
    - Either a complete closed chain is formed or the number of closed breakends must be at least double the number of open breakends. If the max ecDNA candidate JCN < 8, then the entire chain MUST be closed.
    - The total length of closed segments must be > 1500 bases. If the cluster includes only closed segments enclosed by adjacent single or inferred breakends on both sides, then at least one closed segment must have Purple depthWindowCount > 5
    - The sum of the JCN from foldbacks in the cluster + the sum of JCN of junctions from regions internal to the ecDNA segment bounds to regions external (excluding assembled templated insertions) + the maximum JCN of any single or inferred breakend (excluding proximate pairs of clustered breakends with opposite orientation) on closed segments + 4 < reference JCN of DM
        - This rule is intended to ensure that the JCN of the DM could not have been achieved via amplification in a linear chromosome
        - If no foldbacks are DM candidate variants, then the reference JCN is set to the maximum JCN of any candidate DM SV in the cluster. Otherwise it is set to the lowest JCN of the foldback DM candidate variants. This is to allow for the uncertainty of whether foldbacks

are part of the presumptive ecDNA or may have been part of a BFB event that caused the amplification in a linear chromosome

If LINX determines that an ecDNA event has occurred using the above criteria, it will retain and annotate the ecDNA chaining. Other junctions in the cluster with both breakends fully contained within a closed ecDNA segment are then only allowed to link to other variants within the ecDNA OR to the ecDNA forming variants. These links are likely lower JCN disruptions which occurred after the ecDNA was first replicated are present on a subset of the ecDNA.

##### Special considerations for clusters with 2 inversions or 2 translocations

Chains consisting of 2 inversions or 2 translocations that do not meet the criteria for ecDNA are common. Where the breakends overlap they may have multiple plausible paths, in each case either forming a reciprocal inversion / translocations or a deletion / duplication with templated insertion. In cases where breakends cannot be phased, LINX can not uniquely distinguish between these 2 scenarios. LINX implements the following heuristics to attempt predict the event type:

For a pair of translocations:

- If the breakends face away from each other or are cis-phased on both chromosomes, then resolve as RECIP_TRANS (no chaining)
- If the breakends face towards each other and are not cis-phased on both chromosomes then:
  - If double minute rules are satisfied then resolve as DOUBLE_MINUTE
  - If the segment on either chromosome is bounded by LOH resolve then chain the other segments and resolve as a DUP_TI chain.
  - Else resolve as RECIP_TRANS_DUP (no chaining)
- If the breakends face towards each other and are not cis-phased on one chromosome and the breakends face away from each other or are cis-phased on the ohter chromosome then:
  - If the segment on the chromosome with facing breakends is bounded by LOH resolve then chain the other segments and resolve as a DEL_TI chain.
  - Else resolve as RECIP_TRANS_DEL (no chaining)

Similarly, for clusters with a pair of inversions with opposite orientations:

- If the 2 inversions face away from each other and overlap on their outer breakends only, then resolve and chain as a RECIP_INV
- If the 2 inversion face towards each other and overlap on their inner breakends only, then:
  - If double minute rules are satisfied resolve as DOUBLE_MINUTE
  - If either segment is bounded by LOH, chain the other segment and resolve as DUP_TI chain
  - Else chain the outer breakends and resolve as RECIP_INV_DUP
- If one inversion fully encloses the other then:

  - If both inversions have the same JCN and the shorter possible templated insertion is bounded by LOH then chain the shorter templated insertion and resolve as a DEL_TI
  - Else if the inner inversion length < 100K, chain the shorter templated insertion and resolve as RESOLVED_FOLDBACK
  - Otherwise chain the longer of the 2 templated insertions and resolve as a RECIP_INV_DEL_DUP
- If there are 2 foldback inversions facing each other without overlap then
  - If double minute rules are satisfied then resolve as DOUBLE_MINUTE
  - Else resolve as FB_INV_PAIR (This may be formed from a breakage fusion bridge event)

#### Chain annotations

The following data is captured for each templated insertion in a chain:
- Whether the link is assembled
- Distance to the next link and whether it traverses any other breakends or links
- Genic overlap and any exact exon boundary exon matches (ie. a pseudogene)

##### Annotation of local topology

Consecutive breakends with no more than 5kb between them or which are part of the same foldback inversion are grouped together into a local topology group and given an id. The number of TIs formed by chained segments wholly within the local topology group are counted and a topology type is given to the remaining variants based on the breakend orientations. The topology types are categorised as one of the following (after excluding all TIs)
- TI_ONLY - All breakends in the group form templated insertions
- ISOLATED_BE - One breakend only
- DSB - A par of breakends forming a deletion bridge
- FOLDBACK - One foldback only
- FOLDBACK_DSB - A foldback with the outer breakend forming a deletion bridge
- SIMPLE_DUP - A single DUP of <5k bases
- COMPLEX_LINE - Any other cluster that is resolved as LINE
- COMPLEX_FOLDBACK - Any other cluster that includes a foldback
- COMPLEX_OTHER - Any other cluster

#### 4. Gene impact & fusion prediction

##### Annotation of breakends with potential gene impact

For each breakend we search for genes that could be potentially disrupted or fused by the structural variant. To do this we find and annotate the breakend for any transcript that either:

- Has an exon or intron overlapping the breakend
- Has its 5' end downstream of and facing the breakend and less than 100k bases where no other splice acceptor exists closer to the breakend.

Each breakend is additionally annotated for the transcript with the following information:

- **disruptive**: a breakend is disruptive for a particular transcript if the SV is an inversion, translocation or single breakend or a deletion/duplication that overlaps at least part of an exon in the transcript AND the variant is NOT part of a chain which does not disrupt the exon ordering in the transcript. A breakend which is resolved as type 'LINE insertion' is never marked as disruptive.
- **transcript coding context**: UPSTREAM, 5_UTR, CODING, 3_UTR, DOWNSTREAM OR NON_CODING
- **gene orientation**: relative orientation of gene compared to breakend (UPSTREAM or DOWNSTREAM)
- **exonic** (TRUE/FALSE)
- **exact base phase**: The exact base phasing of the current location
- **Next splice site information**: The distance to, phasing of and exon rank of the 1st base of the next facing splice acceptor or donor (note: phasing will be different from exactBasePhase if the breakend is exonic in the transcript or the coding context is upstream. Null if there are no subsequent splice sites in the gene.
- **exon total count**: The total number of exons in the transcript (for reference)
- **transcript biotype**: The ensembl biotype of the transcript

##### Known pathogenic fusions and promiscuous partners

Configuration of pathogenic fusions impacts LINX in 2 ways:

1. Some criteria for fusion calling are relaxed for known fusions due to the high prior likelihood of pathogenic fusions
2. As well as attempting to predict all fusion events, LINX uses the configured list of fusions to determine a subset which are 'reported' as likely pathogenic.

To produce the list of known fusions provided with LINX, a broad literature search was performed to find a comprehensive list of well-known fusions that are highly likely to be pathogenic. The criteria used for inclusion of a particular fusion in the curated list was either multiple independent reports of the fusion, or single case reports with either convincing demonstration of the pathogenicity in a model system or clear response to a therapy targeted to the specific fusion.

The curated fusions were classified into 3 categories:

- **Known fusions** (n= 396) – these are transcript fusions which fuse either the coding regions of 2 genes to form a novel protein or the 5' UTR regions of 2 genes which may lead to increased expression of the 3' partner. A well-known example is TMPRSS2_ERG

- **Known IG enhancer rearrangements** (n= 17) – these are structural rearrangements in B-Cell lymphomas and leukemias that relocate enhancers from one of the @IG regions (IGH,IGK,IGL) to increase expression of a 3' partner. A well-known example is @IGH-MYC
- **Known exon deletions & duplications** (n=11) – these are deletions or duplications of exons in specific exon ranges of a handful of genes which are known or highly likely to be pathogenic.  Common examples are EGFR vII and vIII

A set of 'promiscuous' fusion partners was also determined from this list so that potential novel fusions with fusion partners that have been identified in multiple fusions previously can also be reported as potentially pathogenic. Any gene which was identified in 3 or more known fusions was marked as a promiscuous 5' partner and likewise if it was identified in 3 or more known fusions in our curated list was marked as a promiscuous 3' partner. MYC and CRLF4 were also marked as 3' promiscuous since they feature in known fusions with both IG enhancer and known fusions. FGFR1 is also added as a 5' promiscuous partner.

For 12 promiscuous fusion partners {FGFR1, FGFR2, FGFR3, TMPRSS2, SLC45A3, HMGA2,BRAF, RET, ALK, ROS1, ETV1, ETV4} a specific exon range has been identified as highly promiscuous and is identified in the knowledge base. Fusion reporting criteria are relaxed in these ranges.     10 promiscuous 3' genes {BRAF ,RET ,ROS1 ,ALK ,MET ,NRG1 , NRTK1, NTRK2 & NTRK3) are marked as 'high impact' and also have special treatment in the fusion reporting logic.

A section of @IGH gene stretching from the diversity region to the end of the constant region was also marked as a 'promiscuous IG partner' as it features in many IG fusions.

#### Fusion Prediction

##### Identify fusion candidates

Fusions are predicted in LINX by looking for consecutive and novel splice donor-acceptor pairings that are joined together in derivative chromosomes by either a single structural variant or a continuous chain of structural variants.

For each single SV and for every facing pair of SVs in the same chain identify all viable splice acceptor and splice donor fusion combinations which satisfy the following conditions:

- 3' gene partner must have coding bases
- 5' gene partner transcript must have one of the following ensembl biotypes: 'protein_coding','retained_intron','processed_transcript','nonsense_mediated_decay', 'lincRNA'
- The upstream breakend must fall within the 5' partner transcript and be disruptive to the transcript.
    - The downstream breakend must fall either within the 5' gene or within 100kb upstream.
    - The combined length of all segments in the chain must be less than 150kb.
- The SV or chain must join appropriate contexts of the 5' and 3' genes (see table below) and for coding regions must be inframe after allowing for any skipped exons. For exonic to exonic fusions exact base phasing is also checked as splice acceptor to splice donor phasing. The following table shows allowed contexts:

| | 3’ Fusion Partner Context | | | | | |
|---|---|---|---|---|---|---|
| 5’ Partner Context | Upstream | 5’UTR Intronic | 5’UTR Exonic | Coding Intronic | Coding Exonic | 3’ UTR or Non-Coding |
| Upstream | X (Enhancer) | X (promoter loss) | | | | X (No Downstream Impact) |
| 5’UTR or non-coding Intronic | YES (1) | YES | YES | alt start codon (4) | | |
| 5’UTR or non coding Exonic | exon skipped (3) | | YES | exon skipped & alt start codon (3,4) | alt start codon (4) | |
| Coding Intronic | YES (1) | SEE NOTE (5) | | YES | YES (2) | |
| Coding Exonic | exon skipped (3) | | | | YES | |
| 3’UTR | X (post stop codon in upstream gene) | | | | | |

(1) for breakends in the upstream region of the 3’ partner, the 1st exon is not considered as it does not have a splice acceptor and so the 2nd exon is assumed to be the 3’ fusion partner. 5’ partner coding to 3’ partner upstream is also possible if the 3’ partner coding region starts in the 1st exon.
(2) If fusing intron to exon, the fusion occurs with the next downstream exon, so check against the frame of the end of the exon instead of the exact base.
(3) Exonic to Intronic can occur if alternative splicing causes exon with exonic breakend to be skipped
(4) 5’ partner 5’UTR or non-coding to coding region of 3’ partner can technically make a fusion, but would need to find 1st alternative start codon also in-frame. These are called as out of frame and only reported for known fusions
(5) Coding Intronic to non-coding allowed only when transcript starts on 1st base of the next downstream exon - in this case we fuse to the first base of the gene which is allowed.

##### Special rules for single breakends which align to a site which forms a pathogenic fusion

The post processing steps of GRIDSS annotate the best alignments for the insert sequence of single breakends which cannot be uniquely mapped. If any single breakend has an alignment which would create fusion matching a known fusion in our knowledge base then that fusion is called as if the single breakend was a translocation to that alignment.

##### Special rules for IG rearrangements

In the special case of IG enhancer rearrangements, the rearrangement normally occurs either between the ‘D’ and ‘J’ region (due to RAG mediation D-J recombination failure – common in IGH-BCL2 fusions) or in the switch region just upstream of the constant regions (due to failure of isoform switching mechanisms – common in IGH-MYC rearrangements). In the former case, the Eμ enhancer is the likely driver of elevated expression whereas in the latter the driver is likely the alpha 1,2 & 3 regulatory region enhancer. To predict a relevant rearrangement, LINX requires that the breakend in IG is oriented downstream towards the enhancer regions and is connected to the 5’ UTR of the 3’ gene partner.

##### Special gene specific cases

LINX also has special rules to support unusual biology for a handful of known pathogenic fusions:

- For CIC_DUX4 and IGH_DUX4, the DUX4 end may map to a number of different chromosomal regions, including the telomeric ends of both chromosomes 10q and 4q and the hg19 alt contig GL000228.1.
- For RP11-356O9.1_ETV1 fusion (pathogenic in Prostate cancer) the breakend on the 5' side breakend is permitted to be up to 20kb downstream of RP11-356O9.1.
- In the case of IGH-BCL2, LINX also looks for fusions in the 3'UTR region and up to 40k bases downstream of BCL2 facing in the upstream orientation towards BCL2 (common in Folicular Lymphomas).

##### Prioritise genes and transcripts

Each candidate chained splice acceptor and splice donor fusion pair may have multiple gene fusion transcripts on both the 5' gene and 3' gene that meet the above criteria. Occasionally genes may also share a splice acceptor or splice donor in which case the transcripts of that groups of genes are considered together. LINX prioritizes the potential transcript candidates and choose a single pair of 5' and 3' transcripts via the following criteria in order of priority:

- Fusion is a KNOWN_PAIR or known EXON_DEL_DUP
- A phased fusion is possible
- Chain is not terminated early
- 3' partner biotype is 'protein_coding'
- 3' partner region is INTRONIC or EXONIC
- No exons are skipped
- Best 3' partner transcript, ranked by canonical and then longest (non NMD) protein coding
- Best 5' partner transcript ranked by canonical, then longest protein coding, then longest

If multiple chains link the same 2 genes then they are prioritised again according to the above logic.

##### Reportable fusions

In addition to predicting fusions, LINX also tries to identify likely viable pathogenic fusions and marks as reportable. To maximise precision whilst ensuring high impact fusions are always likely to be reported, the criteria vary by fusion type with more relaxed criteria for known pathogenic pairs due to high prior likelihood. High impact promiscuous fusion partners which may be clinically relevant (including NTRK1-3, BRAF, RET, ROS1, ALK) also have more relaxed criteria

The criteria are summarised in the below table.

| Criteria | KNOWN PAIR | IG KNOWN PAIR | EXON DEL DUP | HIGH IMPACT PROMISCUOUS | PROMISCUOUS OTHER |
|---|---|---|---|---|---|
| Knowledge-base match | GENE PAIR | GENE PAIR | Breakends within EXON RANGE | INTERGENIC ONLY | INTERGENIC ONLY |

| | | | | | |
|---|---|---|---|---|---|
| ‘Nonsense Mediated Decay’ biotype allowed for 3’ partner | FALSE | FALSE | FALSE | FALSE | FALSE |
| Maximum chain links | 4 | 4 | 4 | 4 | 4 |
| Maximum upstream distance for 3’ partner | 100kb | 10kb | NA | 100kb | 10kb |
| Phasing | INFRAME or SKIPPED EXONS | NA | INFRAME or SKIPPED EXONS**,*** | INFRAME or SKIPPED EXONS | INFRAME**** |
| Allow early chain termination or disruption by intermediate splice acceptor or donor | TRUE | TRUE | TRUE | FALSE | FALSE |

* 5’UTR to coding regions are also allowed for KNOWN_PAIR fusions (>1Mb length)
** The breakend and the fused exon must both be in the specified ranges on both 5’ and 3’ side
*** Out of frame also reported for exonic to exonic EXON_DEL_DUP only (under assumption of possible phased indel)
**** Skipped exons are allowed if a known exon range is configured, but only if the breakend and fused exon must be within the specified range on the promiscuous side, and any skipping is allowed on the non-promiscuous gene.

Additionally, LINX checks that the protein domains retained in the 4’ partner may form a viable protein. Specifically The following domains must be preserved intact in the 3’ partner if they exist: Ets domain; Protein kinase domain; Epidermal growth factor-like domain; Ankyrin repeat-containing domain, Basic-leucine zipper domain,High mobility group box domain. The Raf-like Ras-binding domain must be disrupted if it exists (mainly affects BRAF).

Finally linx sets a likelihood for each reported fusion. KNOWN_PAIR, IG_KNOWN_PAIR and EXON_DEL_DUP are set to HIGH likelihood. PROMISCUOUS fusions are set to HIGH likelihood only if the fused exon matches the known exon range, or else LOW otherwise.

#### Amplification, deletion and disruption drivers

##### Homozygous disruption drivers

LINX can optionally take as input a catalog of point mutation, amplification and homozygous deletion drivers which is created by PURPLE based on the raw somatic variant data and determined copy number profile. LINX leverages it’s chaining logic to extend the driver catalog by searching for 2 additional types of biallelic disruptions which disrupt all copies of the gene but do not cause a homozygous deletion in an exonic segment (which is PURPLE’s criteria for homozygous deletion). Specifically LINX searches for 2 additional types of homozygous disruptions:

- **Disruptive breakends -** Any pair of disruptive breakends that form a deletion bridge or are oriented away from each other and both cause the copy number to drop to <0.5 after allowing for the JCN of any overlapping deletion bridges.
- **Disruptive duplications -** Any duplication which has both breakends disruptive in the transcript and a JCN >= flanking copy number at both ends.

##### Linkage of drivers to contributing structural variant clusters

We link each driver in the catalog that is affected by genomic rearrangements (ie. high level amplifications, homozygous deletions and biallelic point mutations in TSG with LOH and the homozygous disruptions found by LINX) to each structural variant cluster which contributed to the driver. 1 or more structural variant clusters may contribute to each event and/or the driver may be caused by a whole chromosome or whole arm event which cannot be mapped to a specific variant but which has caused significant copy number gain or loss

Amplifications drivers are linked to either one or more clusters which explain the gain in copy number over the gene (marked as type 'GAIN'), a gain in copy number in the centromere (marked as type 'GAIN_ARM') or across the whole chromosome (marked as type 'GAIN_CHR'). More than 1 of these factors can be recorded as a cause of amplification if its copy number gain is at least 33% of the largest contributing factor. To determine whether a cluster contributes to gene amplification, the copy number change of all its breakends surrounding the gene are summed into a net cluster copy number gain, and the copy number loss of any opposing clusters are subtracted. If a net gain remains, then the cluster is considered as contributing to the gene amplification.

Homozygous deletion drivers are linked to clusters that are either directly bound by the homozygous deleted region (svDriverType = DEL) or a cluster that bounds the LOH (svDriverType = LOH). Biallelic point mutations drivers in TSG with LOH are linked only to the cluster that bounds the LOH (svDriverType = LOH). If the LOH bounds extend to the whole arm or whole chromosome, then a record is created with linked clusterId is set to NULL and the svDrivertype is marked as LOH_ARM or LOH_CHR respectively.

The supported linkages between drivers and SVs are summarised in the table below

| Driver Type | Events per driver | svDriverTypes |
|---|---|---|
| Amplification | 1+ | GAIN<br>GAIN_ARM<br>GAIN_CHR |
| Homozygous Deletion | 2 | DEL<br>LOH<br>LOH_ARM<br>LOH_CHR |
| Biallelic point mutation in TSG | 1 | LOH<br>LOH_ARM<br>LOH_CHR |

### Visualisation

LINX provides functionality to present detailed visualisation of genomic rearrangements including genic impact in CIRCOS (Krzywinski et al. 2009) format. LINX writes a set of 'VIS' files in a specific format which form the base data to generate the visualisation. The visualisation tool only depends on these files and so in principle any tool could provide SV & CN data in this format.

There are 3 main components in the output of the LINX visualisation each described below.

#### CIRCOS view

The CIRCOS view shows either the complete set of rearrangements in a cluster or set of clusters if 1 or more cluster ids are specified ('cluster mode') or all clusters that touch a chromosome if a chromosome is specified ('chromosome mode').

The CIRCOS view has 6 tracks showing from innermost to outermost:

1. Junctions
2. Minor allele copy number profile
3. Copy number profile
4. Derivative chromosomes
5. Impacted genes (if any)
6. Affected chromosomes

The scaling of both distances and copy numbers in the figure has been modified to make the figure readable. Specifically, the distances between each feature in the chart (either breakend or gene exon start or end) are modified to a log based scale, so that the entire genomic rearrangement spanning millions of bases and multiple chromosomes can be viewed, but that local topology of regions with high densities of breakpoints can be introspected. JCN is set with a linear scale but is scaled down if the maximum cluster JCN exceeds 6 or if the total density of events exceeds a certain maximum to ensure that even the most complex clusters can be introspected. Additionally, if the total number of breakends displayed exceeds XX then the junctions become increasingly transparent such that other features on the plot don't become obscured.

Another key feature of the CIRCOS plot is the ability to trace the derivative chromosome(s). Each segment in the 4th track represents a segment of the derivative chromosome and is linked on both ends either to a centromeric or telomeric end (marked with an open or closed square respective) or a breakend (marked with a track or triangle in the case of foldbacks). Each derivative chromosome can be traced continuously from one telomeric / centromeric end to another (or to a single breakend if one is reached) by following a continuous series of segments and breakends. To make this easier to follow, each time a new segment is connected on a chromosome the segment is offset outwards slightly. Hence the derivative chromosomes can be traced from the inside to the outside of the 4th track of the diagram. A cluster may contain 1 or more derivative chromosomes. In cluster mode each derivative chromosome will be shown in a different colour for ease of viewing (with a maximum of 10 colours after which all derivative chromosomes are shown in black). In chromosome mode, derivative chromosomes will be shown in the same colours, but each cluster is shown in a different colour. Red and green are reserved for simple deletions and tandem duplications respectively. Since there may be many of these on a single chromosome, telomeric and centromeric connectors are not shown for these simple variant types. Light blue is also reserved for LINE clusters which can also be frequent in samples with highly deregulated LINE machinery.

Most of the possible annotations are shown in the LINX visualisation guide (see figure 1B)]. Additionally, 2 types of genomic regions which are frequently disrupted in tumor genomes are indicated using light grey shading on the copy number regions in the 3rd track. For known LINE source elements the green copy number section is shaded and for known fragile sites the red copy number section is shaded light grey.

#### Chromosome view

Since the CIRCOS only represents a part of the genome, the chromosome view is provided to indicate which parts of the genome is shown. Each of the chromosomes included in the cluster(s) shown is displayed. The part of the chromosome that is included in the figure is highlighted in the colour matching the colour used in the outer ring of the CIRCOS. The banding and location of the centromere on each chromosome is also shown.

An example of the chromosome view is shown below indicating the cluster includes a large section of chromosome 7 including the centromere and a small slither of chromosome 15 on the Q arm:

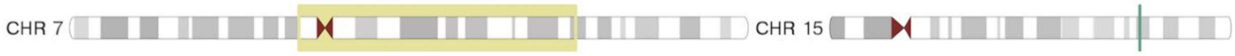


#### Fusion view

The fusion view is added for reportable fusions only in LINX. It's purpose is to show the predicted structure of the fused gene. The fusion includes the fused segments of both the 5' and 3' partner in blue and red and always reads from left to right. The gene representation for each genes and follows the standard conventions of thick bands for coding regions, thinner bands for 5' UTR and 3' UTR exonic regions and thin lines to represent the intronic sections. Protein domains are shown in coloured bands across the exons which they include and are labeled in the accompanying legend. As with the CIRCOS view the lengths of exonic gene segments in the fusion view are scaled by a log scale to improve readability. The intronic segments are set to a fixed segment length regardless of the length.

The fused gene is shown up to and including the breakend on either side that is connected either directly in the case of a simple fusion or via a chain in a chained fusion. If LINX predicts that one or more exons are skipped, then the skipped exonic segments and protein domain sections are faded.

For example in the following TMPRSS2-ERG fusion, exons 3, 4 & 5 are faded and the LDL domain is also faded, indicating that LINX predicts these exons are skipped in order to make a viable in-frame protein, despite the break end occuring after the 5th exon:

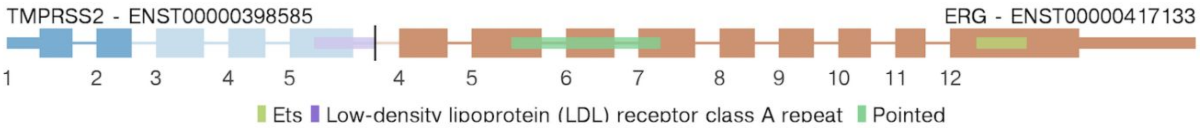


A similar case can be seen in this example where the 5'UTR region of NDRG1 is fused upstream of the 1st exon in PLAG1. Since the 1st exon of a gene has no splice acceptor, LINX predicts the 1st exon is skipped and it is faded on the chart with the fusion connecting to the start of exon 2 which also begins in the 5'UTR region of PLAG1:

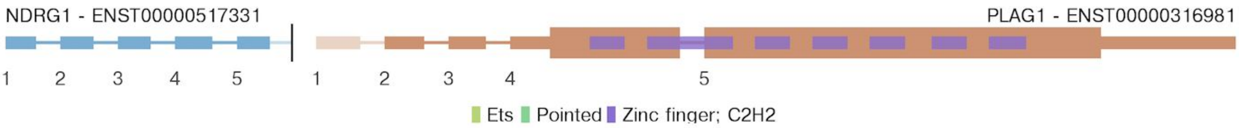


Even when no exons are skipped, a protein domain may be partially disrupted if it extends to an exon the other side of the breakpoint. In the below example, you can see the SH2 protein domain extends before exon 16 and is faded and disrupted:

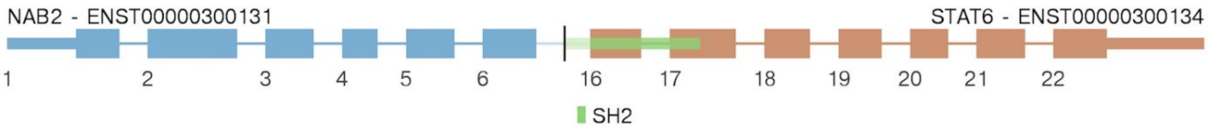


Whilst fusions are normally intronic, rare exonic to exonic fusions do occur. The below figure shows a CIC-FOX04 example where the 2 exons are directly fused

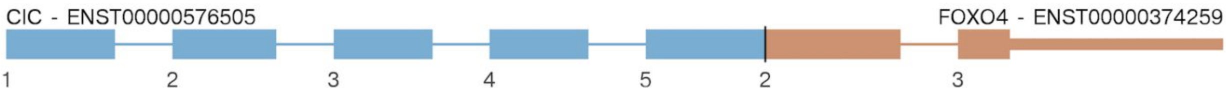
