## Supplementary Note 1 for "Unscrambling cancer genomes via integrated analysis of structural variation and copy number"

### Introduction

COLO829 is a widely studied melanoma tumor-normal paired cell line, that is used as a somatic reference standard for benchmarking in next generation sequencing.  This document uses COLO829 LINX output as an example to give insight into how to interpret a diversity of visualisations in tumor genomes.

### Running LINX on the COLO829T data

Instructions for downloading LINX and it's required resources are provided on github (https://github.com/hartwigmedical/hmftools/blob/master/sv-linx/README.md).  The input files to run LINX on COLO829T are included in supplementary file 2 in the /purple directory.

LINX can be run on the COLO829T data with the following command:

```
java -jar sv-linx.jar
   -sample COLO829T
   -sv_vcf /path_to_purple_vcf/
   -purple_dir /path_to_purple_data_files/
   -fragile_site_file fragile_sites.csv
   -line_element_file line_elements.csv
   -replication_origins_file heli_rep_origins.bed
   -viral_hosts_file viral_host_ref.csv
   -gene_transcripts_dir /path_to_ensembl_data_cache/
   -check_fusions
   -known_fusion_file known_fusion_data.csv
   -check_drivers
   -driver_gene_panel DriverGenePanel.tsv
   -output_dir /output_dir/
   -log_debug
```

### Visualising clusters in COLO829T

LINX identifies 61 clusters in COLO829T which can be found in the linx.clusters.tsv file in the /linx directory. One cluster (Cluster 0) is a low VAF translocation which is filtered as 'ARTIFACT' by LINX. The remaining 60 clusters consist of

- 9 simple tandem duplications
- 35 simple deletions
- 2 synthetic deletions
- 1 synthetic unbalanced translocation
- 1 pair of single breakends that is treated as an implied duplication
- 2 LINE insertions
- 6 complex clusters
- 4 incomplete clusters (all unclustered single or inferred breakends)

Circos must be installed to produce the visualisations. To generate visualisations for each of these clusters you can run the command:

```
java -cp sv-linx.jar com.hartwig.hmftools.linx.visualiser.SvVisualiser
      -gene_transcripts_dir /path_to_ensembl_data_cache/
      -plot_out vis/
      -data_out vis/
      -segment COLO829v003T.linx.vis_segments.tsv
      -link COLO829v003T.linx.vis_sv_data.tsv
      -exon COLO829v003T.linx.vis_gene_exon.tsv
      -cna COLO829v003T.linx.vis_copy_number.tsv
      -circos /data/common/tools/circos_v0.69.6/bin/circos
      -protein_domain COLO829v003T.linx.vis_protein_domain.tsv
      -fusion COLO829v003T.linx.vis_fusion.tsv
      -sample COLO829v003T
```

This produces a picture of each chromosome and also of each non trivial variant cluster (i.e. excluding simple deletion or duplication). Alternatively, the LINX visualiser can be run for a single chromosome or cluster by specifying a -chromsome or -clusterId parameter respectively.

Example plots for each of the types of clusters classified by LINX in COLO829T are described below. The visualization guide in figure 1B of the manuscript can be used to aid interpretation of the figures.

Simple Deletion

Cluster 67 is a simple deletion on chromosome 10 of 12kb that leads to a homozygous loss of exon 6 in PTEN.   Note that simple deletions and tandem duplications will not draw by default unless they affect a gene.

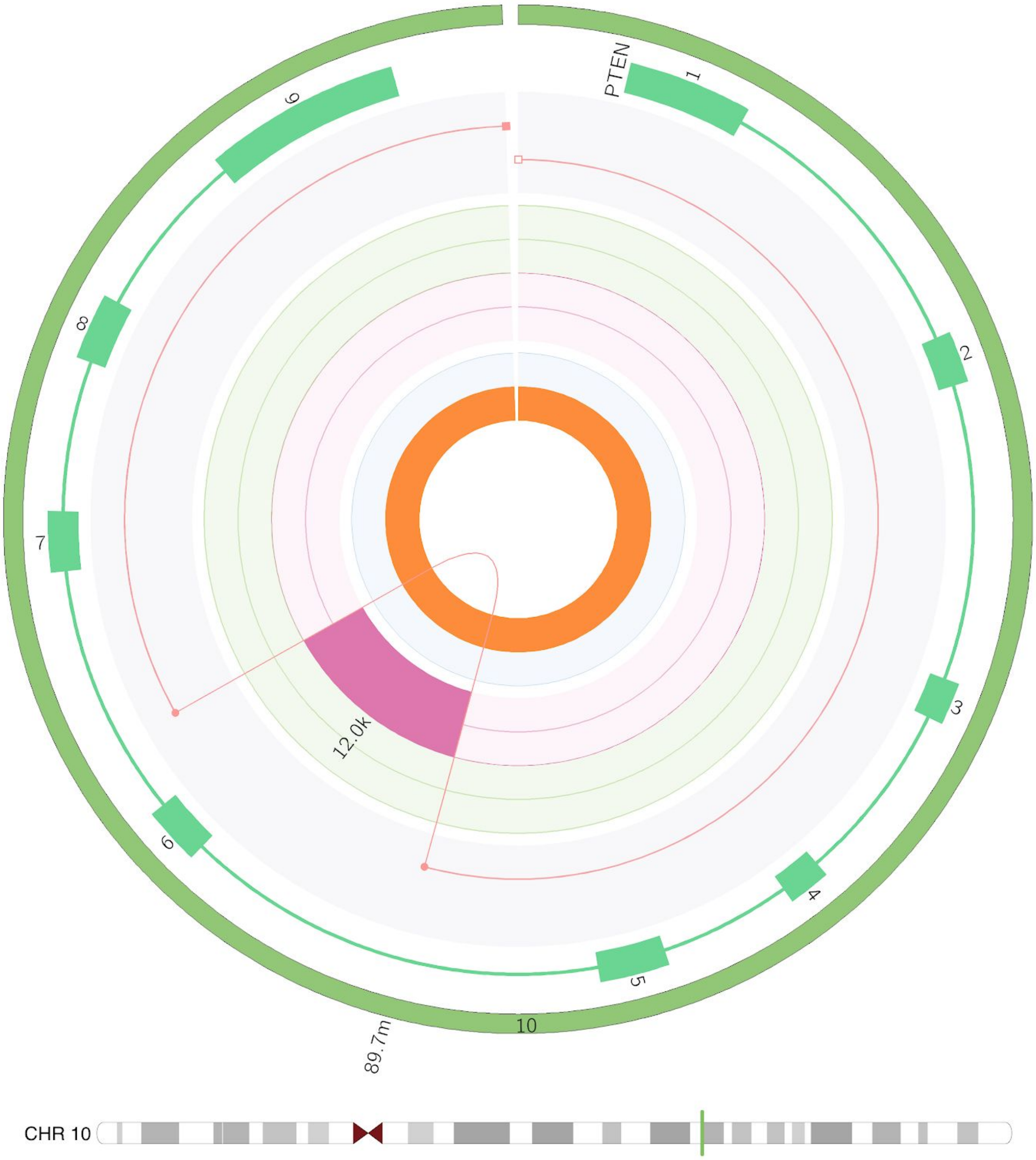

#### Single breakend (incomplete cluster)

Cluster 59 is a single breakend on chromosome 9 with JCN of 3, that causes a LOH from 30.3M to the 9P arm telomere. The other end of the single breakend is unmappable. The LOH contributes to a biallelic driver in CDKN2A which also has a missense variant (not shown)

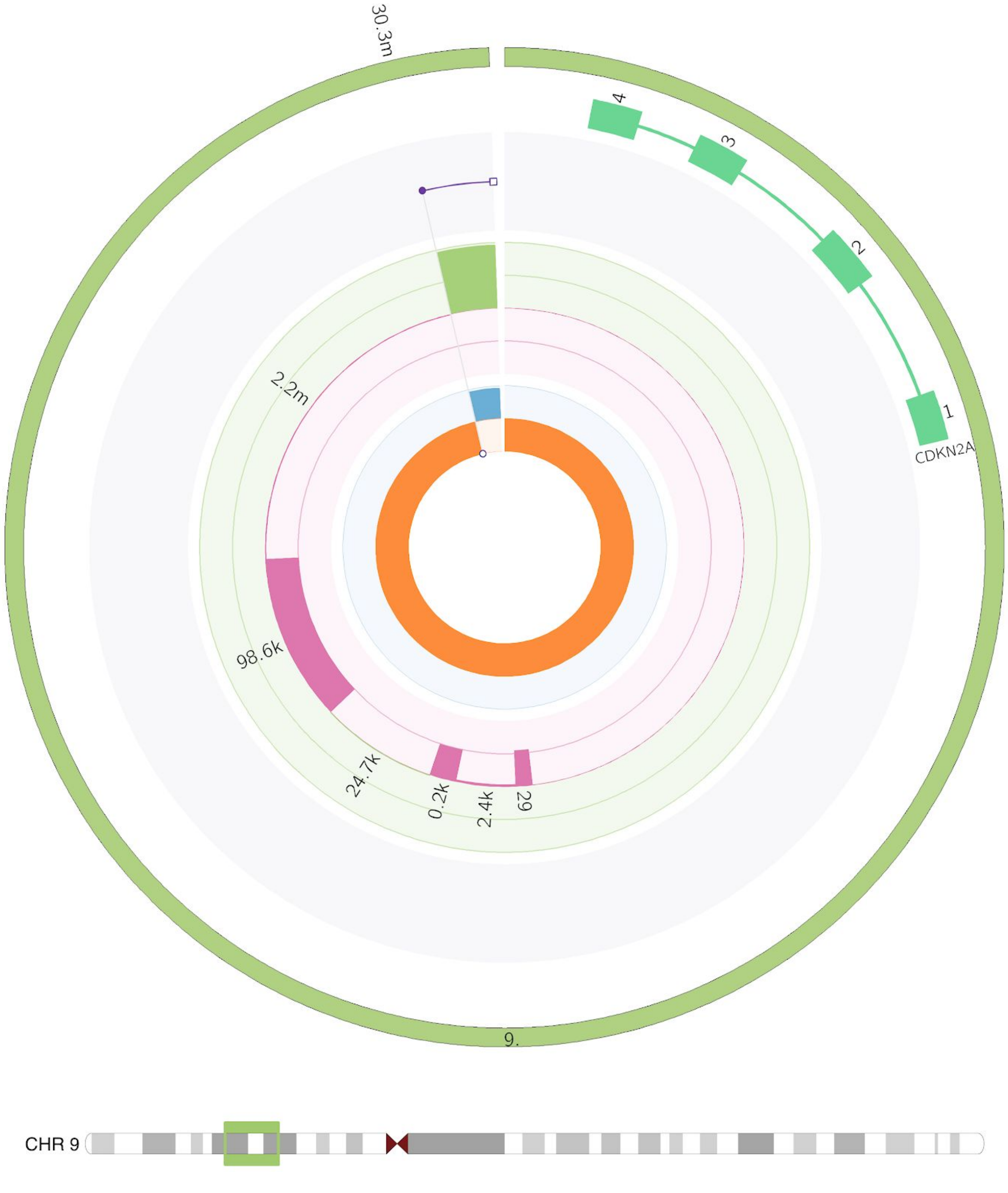

Synthetic Deletion

LINX simplifies variants that form shards of <1kb insertions and will classify as ‘synthetic’ if removal of the shards allows it to be resolved as a 1 or 2 break cluster type. Cluster 13 from COLO829T is an example of a synthetic deletion on chromosome 15 with a 100 base insertion from an otherwise undamaged region of chromosome 6

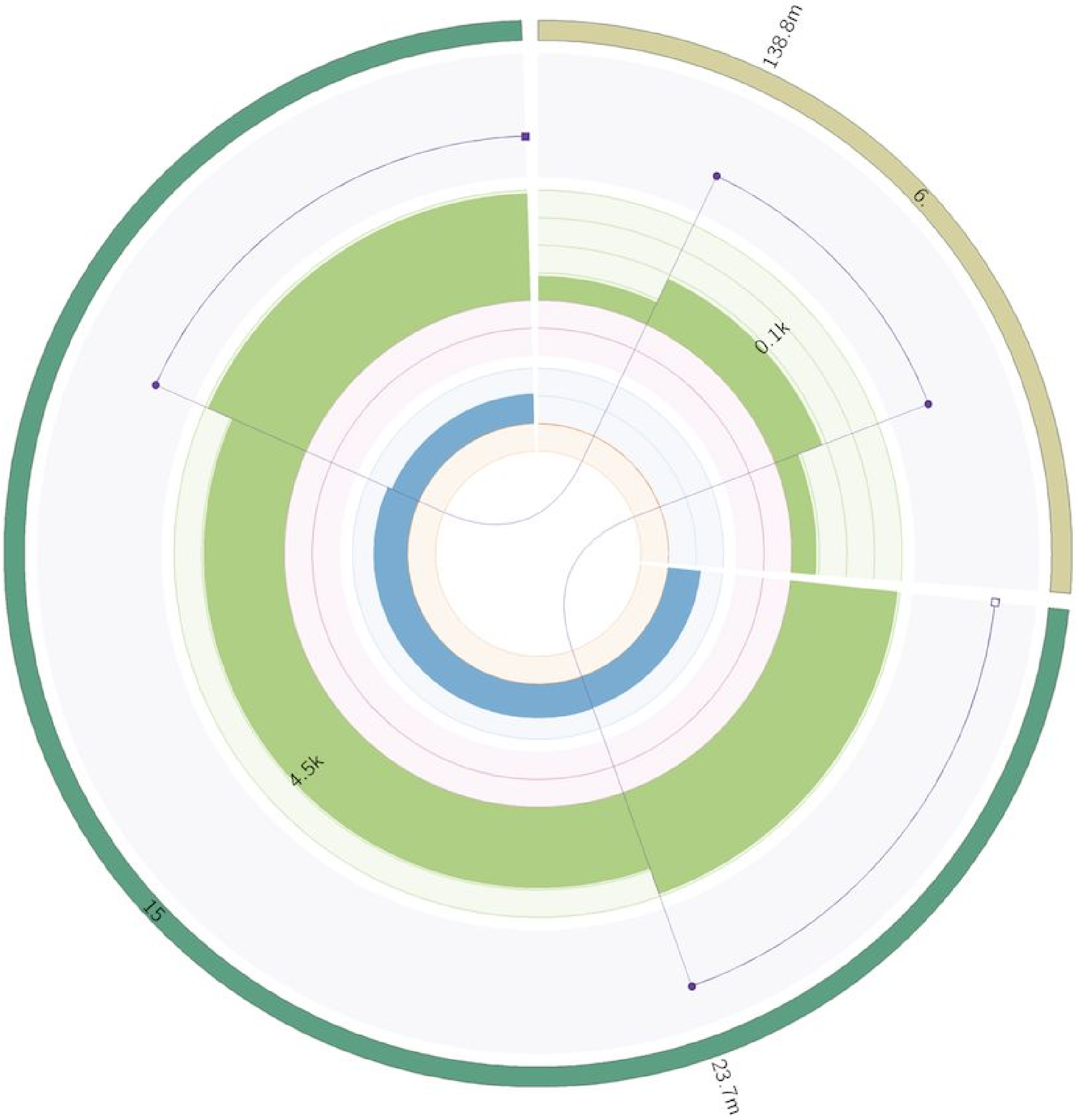

#### Synthetic Unbalanced Translocation

Cluster 64 is an unbalanced translocation has occurred between chromosome 1 and 10 causing a LOH for much of on chromosome 1 and chromosome 10 including the PTEN driver gene (the copy number impact of a homozygous deletion of exon 6 of PTEN can also be seen on the circos in pink, but from an unrelated 2nd hit event).   A small shard of 67 bases has been inserted from chromosome 10 in the unbalanced translocation, so the event is classified as a 'synthetic' unbalanced translocation.

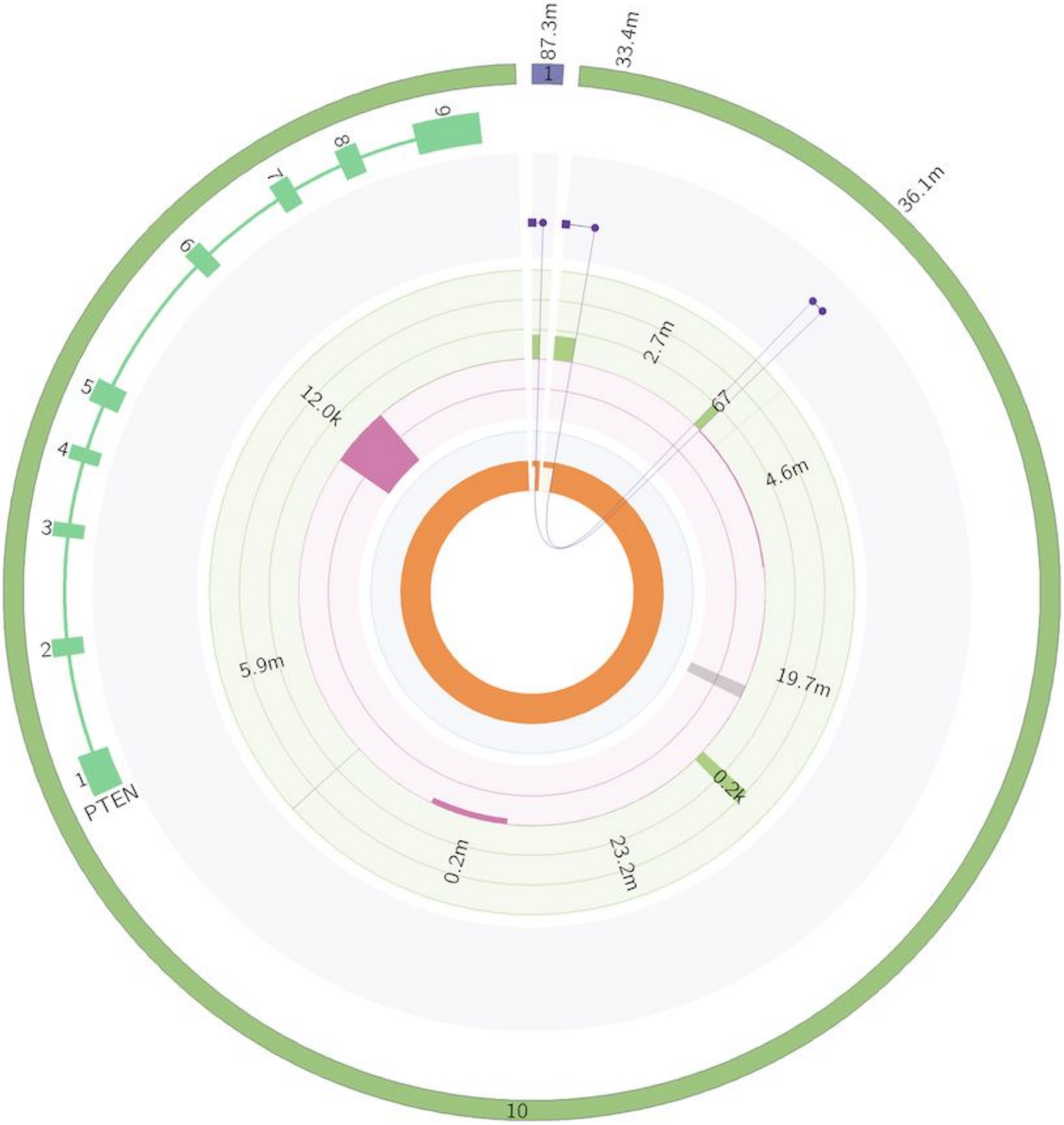

#### LINE Insertion

2 single breakends form the insertion site of a likely LINE insertion on chromosome 12. The LINE source element cannot be uniquely mapped in this case and the LINE classification is made based on the POLYA insertion sequence (not shown). The open circles on the edge of the innermost circle are used by LINX to represent single breakends.

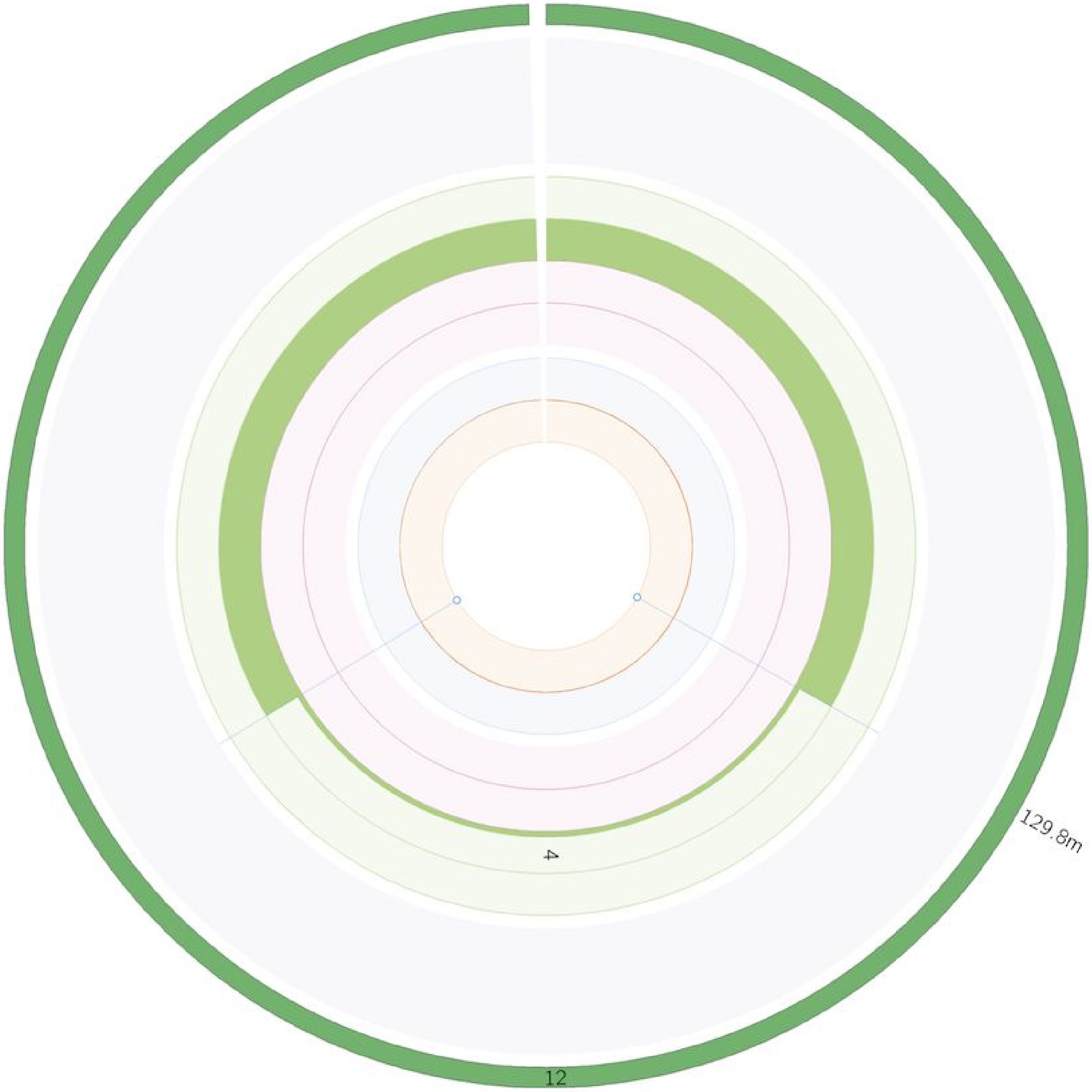

#### COMPLEX Clusters

6 clusters are classified as ‘complex’ by LINX in COLO829T. The most complex rearrangement is a highly amplified structure predominantly on chromosome 3 that includes shards inserted from chromosome 6, 10 and 12 without further DNA damage on those chromosomes. The amplification is caused by 2 simple foldback inversions and 2 chained foldback inversions (with the inserted shards), with foldback breakends identified with a triangle marker in the image. A single breakend at position 25.3M is also clustered and likely resolves the breakage fusion bridge cycle:

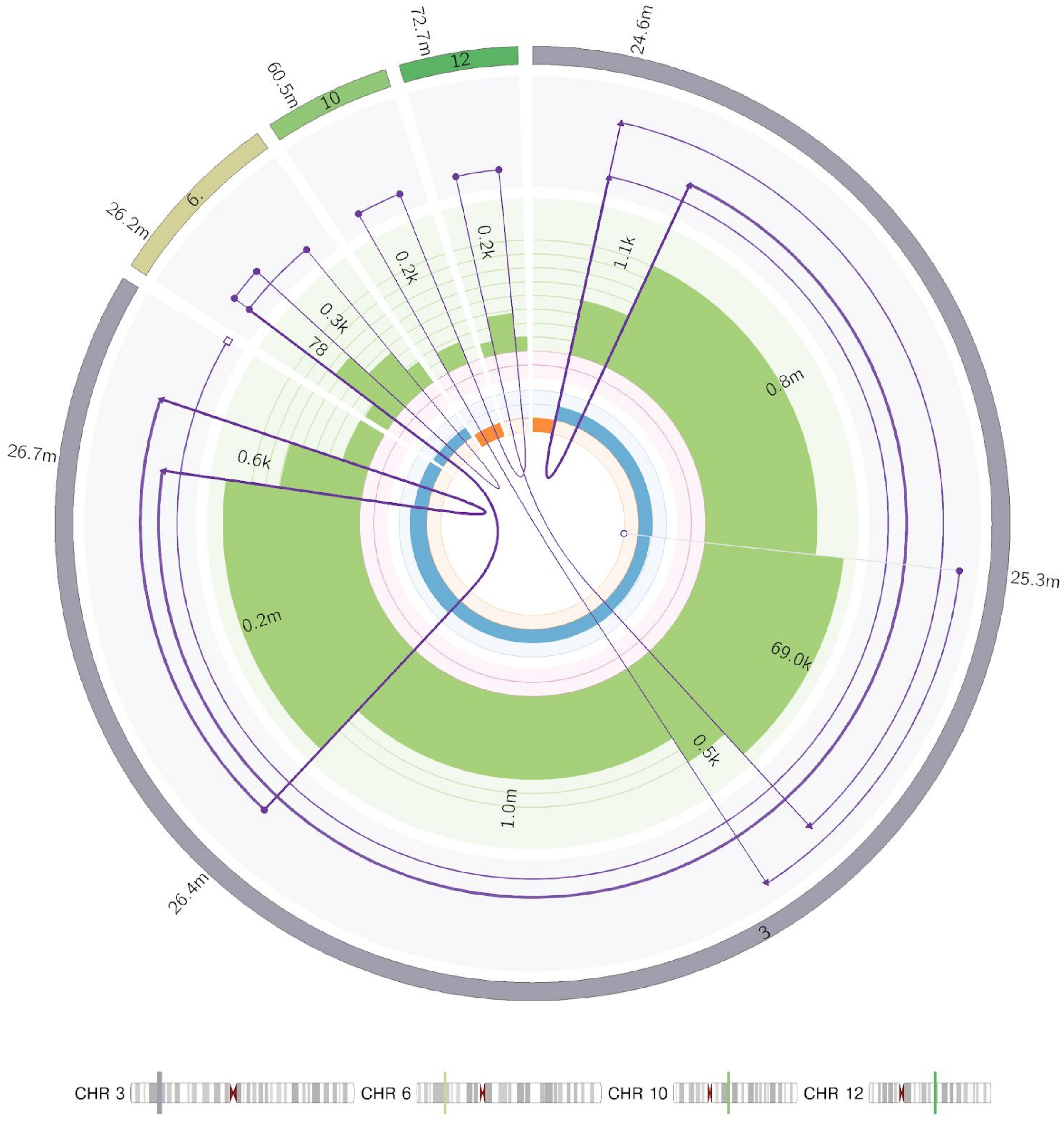

A complex event also occurred on chromosome 9 between 28M and 28.2M and involves 2 inversions and a duplication, likely caused by multiple concurrent double stranded breaks in this region 2 segments are retained and 1 long segment and 2 short gaps are lost.

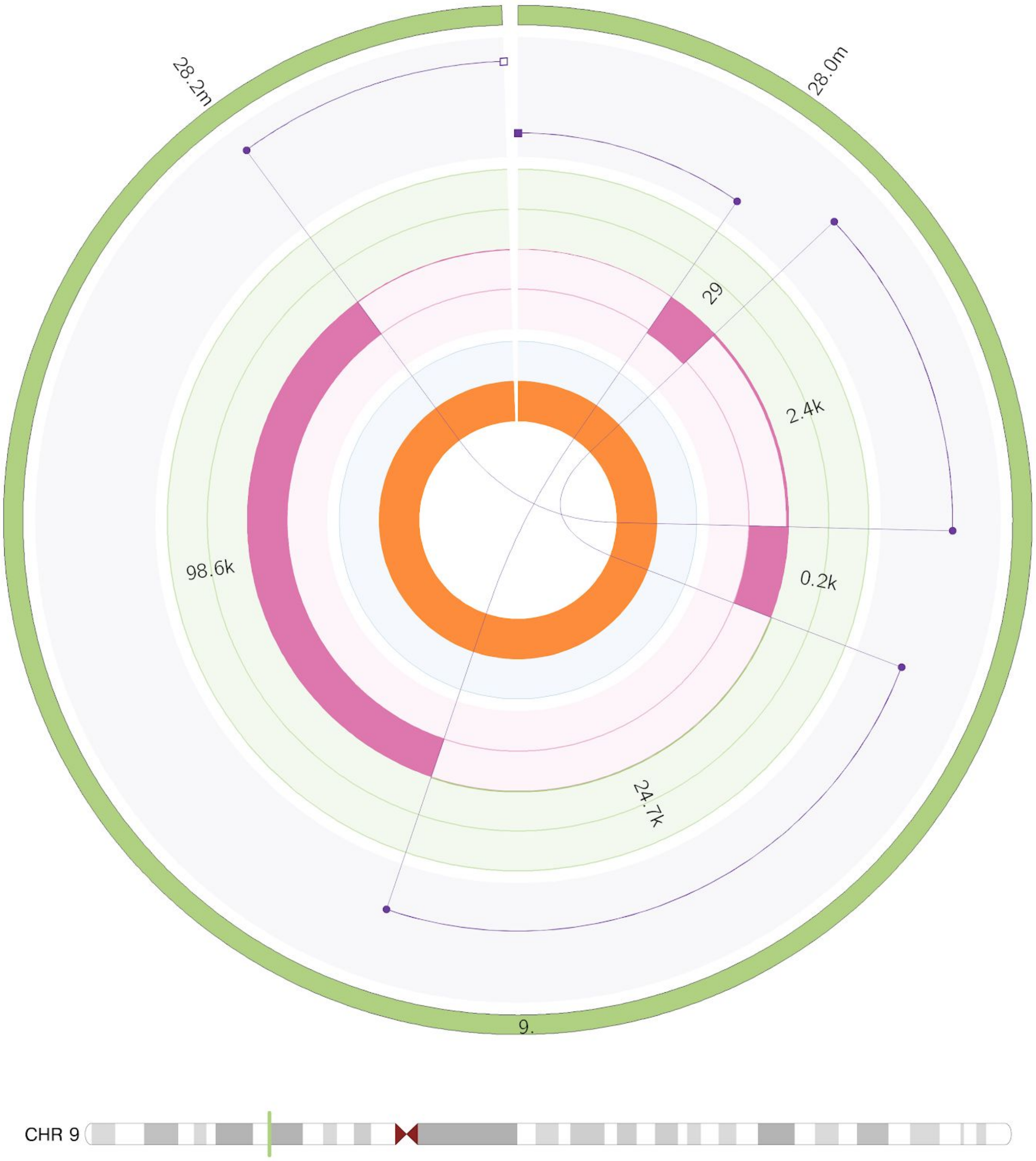

A 3rd complex rearrangement is on chromosome 7 and involves an inversion, a duplication and a single breakend. The rearrangement has caused a LOH in the region from 125.7M to 126.1M and amplification at the telomeric end of the chromosome 7Q arm.  It is unknown where the single breakend connects to in the genome

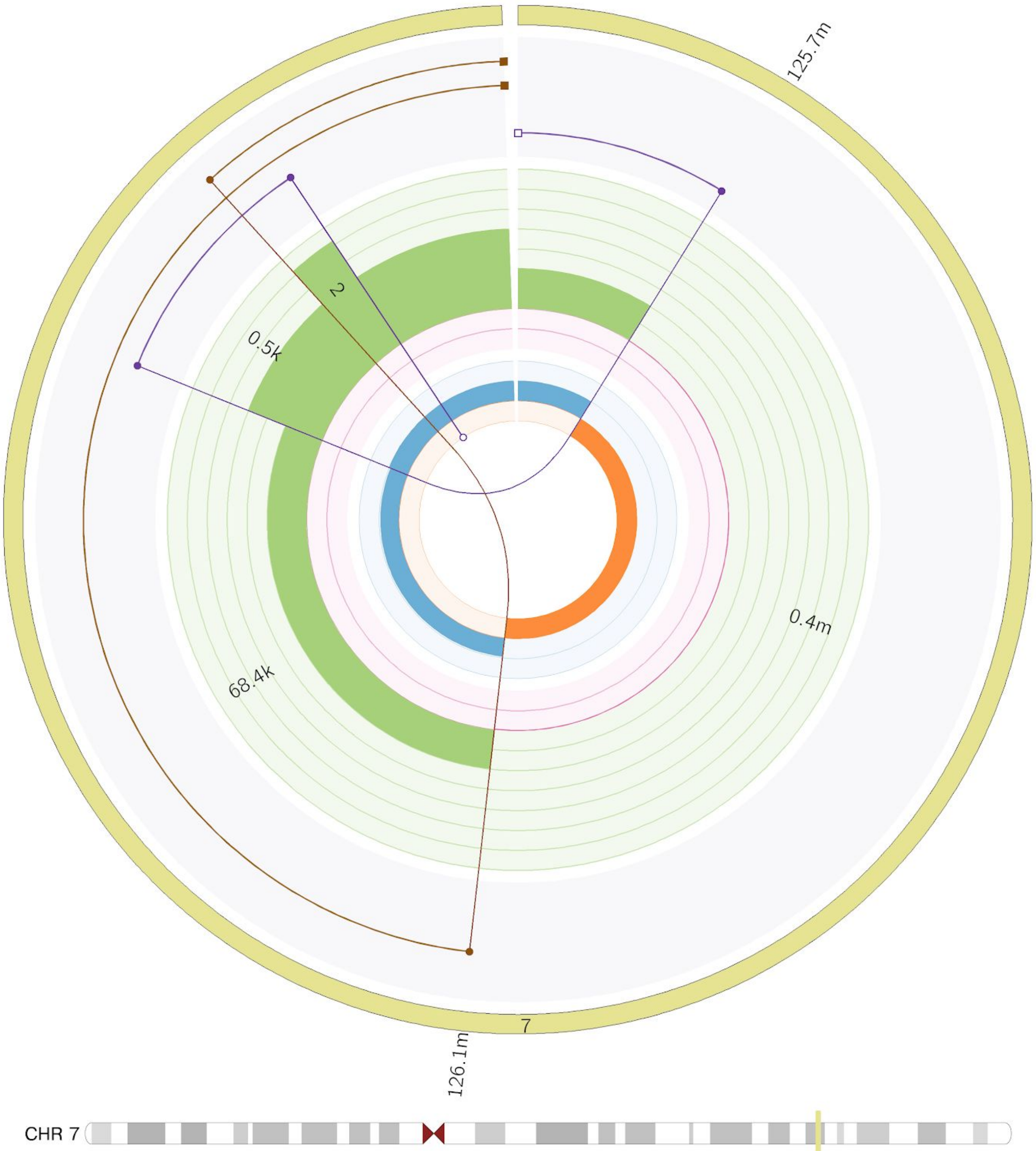

The 4th complex rearrangement involves 3 translocations on 2 distinct derivative chromosomes. The first derivative chromosome inserts a 73.4kb segment of chromosome 10 into a 800 base gap in chromosome 19 and is shown in brown. The 2nd derivative chromosome (purple) is a translocation from chromosome 10 to 18 which has caused amplification of the centromeric end of the 18P arm. The 2 derivative chromosomes events are clustered together due to the LOH deletion bridge in between. An LOH of the first 7.1M bases of chr 10 has also occurred in the same event.

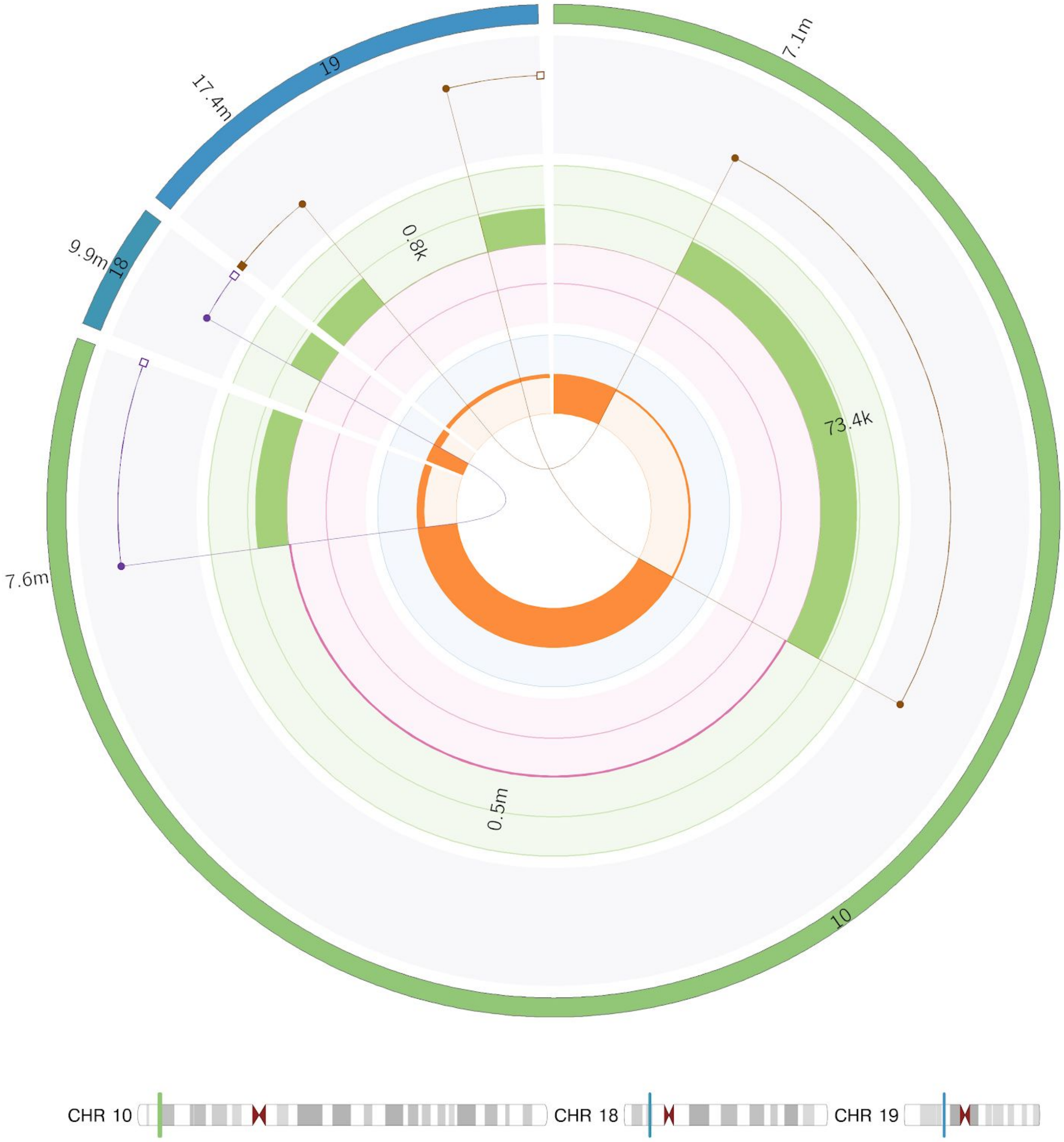

A 5th complex rearrangement primarily involves chromosome 15, causing a LOH on a large segment of chromosome 15 from 24m to 84.8m. The q telomere of chromosome 15 is linked by translocation to the q centromere of chromosome 7.  A pair of facing foldbacks amplify the centromeric region of 15Q.  One of the foldbacks is synthetic with a short shard of 200 bases inserted from chromosome 20.

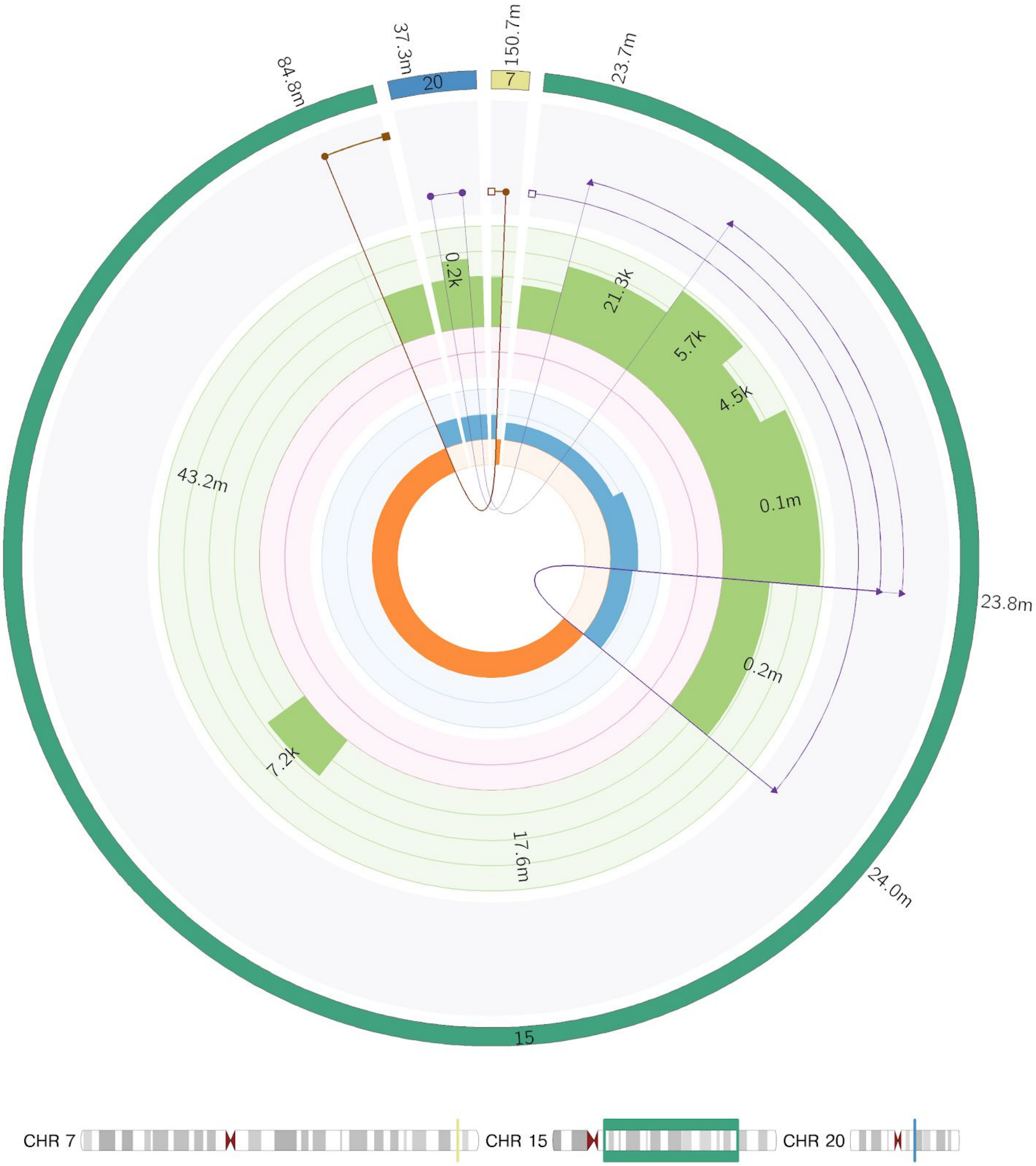


The final complex rearrangement is a subclonal rearrangement of 5 junctions on chromosome 8 that affects 2 regions of the Q arm of chromosome 18 around 78.5M and 112.1M.  At both loci multiple breaks have occurred leading to 3 short chromosomal segments and 1 long segment being rearranged. The damage appears to be localised as the derivative chromosome can be chained consistently from centromere to telomere.

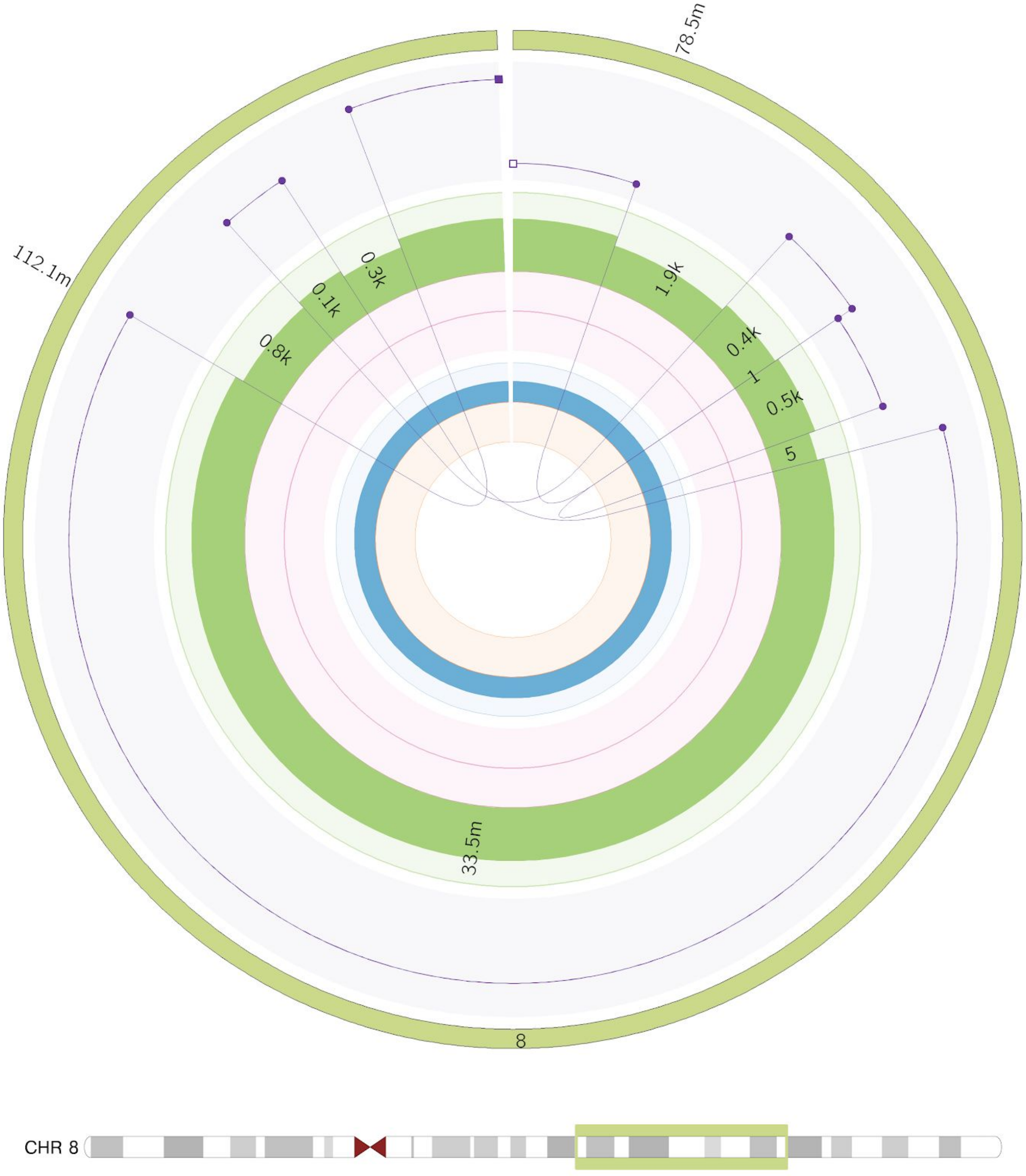

#### Chromosome View

An alternative way to view LINX output is by chromosome view. This may be used to visualise all the clusters with breakends originating from a particular chromosome. The below example shows multiple deletions on Chromosome X. Note, for simple deletions, duplications and insertions, the chromosomal segments are not shown as there can be many in highly rearranged samples. Only the junction is shown with a fixed red colour for deletions, green for duplications and light blue for insertions. Other clusters will each have a distinct colour. Note that for known LINE source elements the green copy number section is shaded and for known fragile sites the red copy number section is shaded light grey. In COLO829, 4 of the 5 deletions on chromosome X are contained within the DMD fragile site as shown.

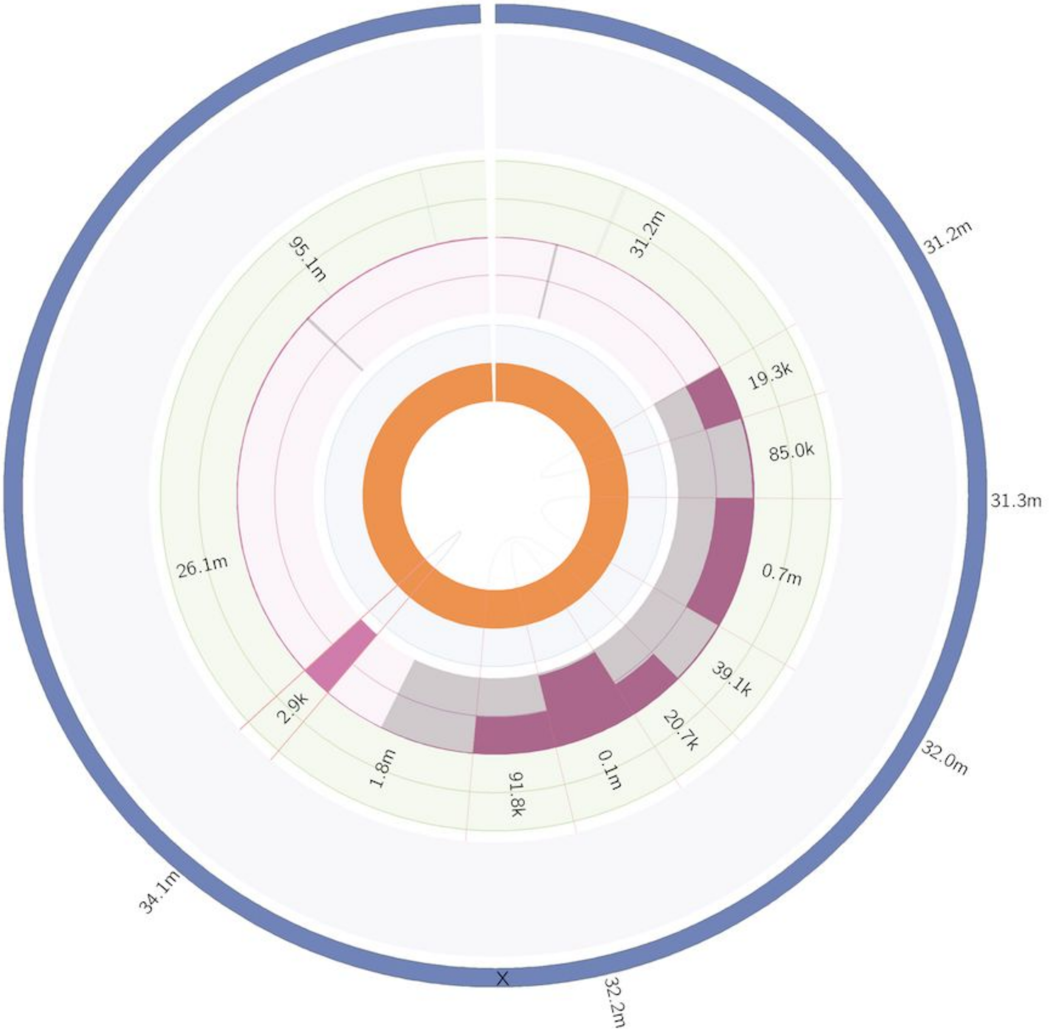
